# Sister chromatid separation determines the proliferative properties upon whole-genome duplication via homologous chromosome arrangement

**DOI:** 10.1101/2025.08.27.672736

**Authors:** Masaya Inoko, Guang Yang, Yuki Tsukada, Ryota Uehara

## Abstract

Whole-genome duplication (WGD) of diploid cells triggers various cell fates, such as cell death, cell cycle arrest, and proliferation with chromosome instability, contributing to broad bioprocesses, including differentiation, tumorigenesis, or aging. However, factors determining the post-WGD cell fates remain largely unknown. In this study, we found that cytokinesis failure (CF) and mitotic slippage (MS), two major routes of WGD induction, differentially affected post-WGD viability and proliferation in human cells. Quantitative live imaging revealed poorer survivability of cells upon multipolar chromosome segregation at the first mitosis after MS than CF. Chromosome-specific labeling showed that the inefficient sister chromatid separation upon MS caused more skewed homologous chromosome distribution than CF. The skewed homologue distribution frequently led to physical isolation (> 10 μm) of the centrosomes from all homologous centromeres, hindering these centrosomes from capturing any of these homologues. The difference in the frequency of this nullisomic chromosome segregation between MS and CF at least partially explained their difference in the viability of the subsequent daughter cells. Moreover, artificial separation of sister chromatids upon MS improved the evenness of homologue distribution, suppressed nullisomic homologue segregation in the following mitosis, and significantly restored the viability of their daughter cells. These results demonstrate the geometric arrangement of homologous chromosomes, defined by the presence or absence of sufficient sister chromatid separation upon WGD, as a key factor determining the proliferative characteristics of subsequent progenies. Our findings would provide a clue to understanding the route-dependent outcomes of WGD in cell fate determination in different bioprocesses.

**Significance:** Whole-genome duplication (WGD), doubling of cellular content through skipping cell division after DNA synthesis, drives cellular diversification in development, aging, tumorigenesis, or evolution. While various mechanisms of WGD are featured in different biological contexts, the potential impacts of differences in WGD mechanisms on resulting cellular properties have been overlooked. Here, we discovered that two major mechanisms of WGD, mitotic slippage and cytokinesis failure, differentially affect proliferative characteristics of post-WGD cells through their contrasting intracellular reorganizations. Mitotic slippage, occurring with inefficient sister chromatid separation, led to a skewed homologous chromosome distribution compared to cytokinesis failure, fueling lethal chromosome loss by non-random chromosome segregation immediately after WGD. Our findings provide insights into the context-dependent preferences for WGD mechanisms.

## Introduction

Whole-genome duplication (WGD) contributes to the progression of broad biological phenomena such as development, aging, cancer, and evolution (1–3). Different cellular processes lead to WGD, causing various cellular outcomes, including cell death, differentiation, senescence, or proliferation with genome instability (4–6). The complexity and diversity of causes and effects of WGD make it challenging to specify key factors affecting the characteristics of WGD cells. Nevertheless, identification of such factors is important for understanding the mechanisms of the different biological phenomena mentioned above.

Cell division failure is a critical cause of WGD in both physiological and pathological processes. There are two main types of cell division failure, cytokinesis failure (CF) and mitotic slippage (MS), leading to WGD (6). CF occurs through defects in dividing the cytoplasm after normal chromosome segregation, forming a binucleated cell that contains two diploid nuclei. Besides spontaneous errors, physiological CF occurs in specific cell types in different tissues, such as hepatocytes, cardiomyocytes, or mammary epithelial cells (7–9). In case of hepatocytes, developmentally programmed suppression of cytokinesis factors mediates the progression of WGD (8). Cancer cells also undergo frequent CF, which is explained, at least in part, by oncogene-mediated suppression of cytokinesis factors (10, 11).

MS is the premature exit from mitosis without chromosome segregation, resulting in a mononucleated cell that contains a single tetraploid nucleus (6). Chromosome capture by spindle microtubules is a stochastic, error-prone process, and spontaneous defects in bipolar spindle formation cause a risk of chromosome segregation failure. Normally, defects in bipolar kinetochore-microtubule attachment block silencing of the spindle assembly checkpoint (SAC), which arrests mitotic progression until the defects are resolved. Suppression of SAC through loss-of-function of its components or attenuation of cdk1-dependent maintenance of the mitotic state lets cells escape mitotic arrest, even without completing bipolar kinetochore-microtubule attachment (12). The premature mitotic exit, if occurring without spindle assembly, results in WGD with a single nucleus (12–14). Various cancer cells manifest altered expression of SAC components, resulting in attenuated SAC function (15–18). Similarly, cancer cells with low cdk1 activity show poor ability to undergo mitotic arrest, which is linked to their proneness to MS upon anticancer drug treatment (19). Moreover, overexpression of v-Src oncogene induces MS by suppressing cdk1 activity (20). In contrast to CF, physiological MS has been found in limited cases, such as megakaryocyte formation or DNA-damage-induced keratinocyte differentiation (21, 22). Therefore, while CF and MS both result in WGD, their cellular causes, geometric features, and prevalence in biological contexts are different. However, it remains largely unknown whether CF and MS differentially affect post-WGD cell fates.

Several factors have been found to determine post-WGD cell fates. The p53 signaling pathway is the most studied mechanism, which is activated upon WGD and mediates cell cycle arrest, senescence, or apoptosis during interphase immediately after WGD (23). Upon WGD, the centrosome number doubles, which induces inhibition or cleavage of Mdm2 to stabilize p53 through the activation of the Hippo signaling pathway or the PIDDosome multiprotein complex, respectively (24, 25). On the other hand, overexpression of cyclin D oncogene allows WGD cells to overcome the p53-mediated cell cycle arrest and proliferate (26, 27). Moreover, proliferative post-WGD subclones established from p53-proficient diploid cell lines frequently manifest altered p53 signaling (28), possibly enabling them to override the WGD-linked anti-proliferative mechanisms.

Once cells evade cell cycle arrest or death immediately after WGD, they enter the first mitosis with 4 centrosomes. A part of these WGD cells cluster the excess centrosomes to form “pseudo-bipolar” spindles to conduct bipolar chromosome segregation (29–31). Moreover, the pseudo-bipolar spindle formation with asymmetric centrosome clustering (i.e., segregating in 1:3 ratio) contributes to efficient removal of the excess centrosomes in their progenies (32). Otherwise, WGD cells form multipolar spindles with the excess centrosomes, driving chromosome segregation into more than 2 daughter cells. Early studies by Boveri proposed that multipolar chromosome segregation occurs in a stochastic fashion, leading to drastic chromosome copy number alterations (33). A substantial proportion of such multipolar daughters are supposed to die due to their lethal karyotypes. On the other hand, the remaining survivors potentially contribute to clonal evolution of WGD progenies with gross aneuploidy, hence driving phenotypic diversification (34–36). Indeed, a frequent multipolar cell division results in the formation of aneuploid progenies after physiological WGD in hepatocytes (37). Therefore, the quality of multipolar chromosome segregation in the early post-WGD phase can be one of the critical factors determining the characteristics of post-WGD cell populations. Importantly, most of the above-mentioned post-WGD alterations (i.e., the excess centrosome gaining, p53 signaling, or multipolar spindle defects) occur in common either after CF or MS (25, 27), having drawn no attention to possible impacts of differences in the route of WGD induction.

In this study, we systematically compared post-WGD cell proliferation after MS or CF using HCT116, a near-diploid human cell line commonly used in ploidy comparative studies (38).

## Results

### MS-derived WGD cells are less proliferative than CF-derived ones

To gain insight into potential differences in properties of WGD cells derived from different types of cell division defects, we compared cell proliferation following pharmacological induction of MS or CF in HCT116. MS-derived WGD (MS-WGD, hereafter) was induced by arresting cells at prometaphase by nocodazole shake-off, followed by co-treating nocodazole and a cdk1 inhibitor RO3306 for mitotic exit without spindle formation (Fig. 1A). CF-derived WGD (CF-WGD) was induced by removing nocodazole from the above prometaphase-arrested cells, and subsequently treating cytochalasin B (Fig. 1A). These compounds were thoroughly washed out after 2-h treatment. MS- and CF-WGD induction resulted in equivalent increases in DNA content (SI Appendix, Fig. S1A and B). Efficiency of MS- or CF-WGD induction, assessed by possession of the excess centrosomes, was 85% or 82%, respectively (SI Appendix, Fig. S1B and C). As expected, most MS- or CF-WGD cells had a single nucleus or two nuclei, respectively (Fig. 1B and C).

**Figure 1.**
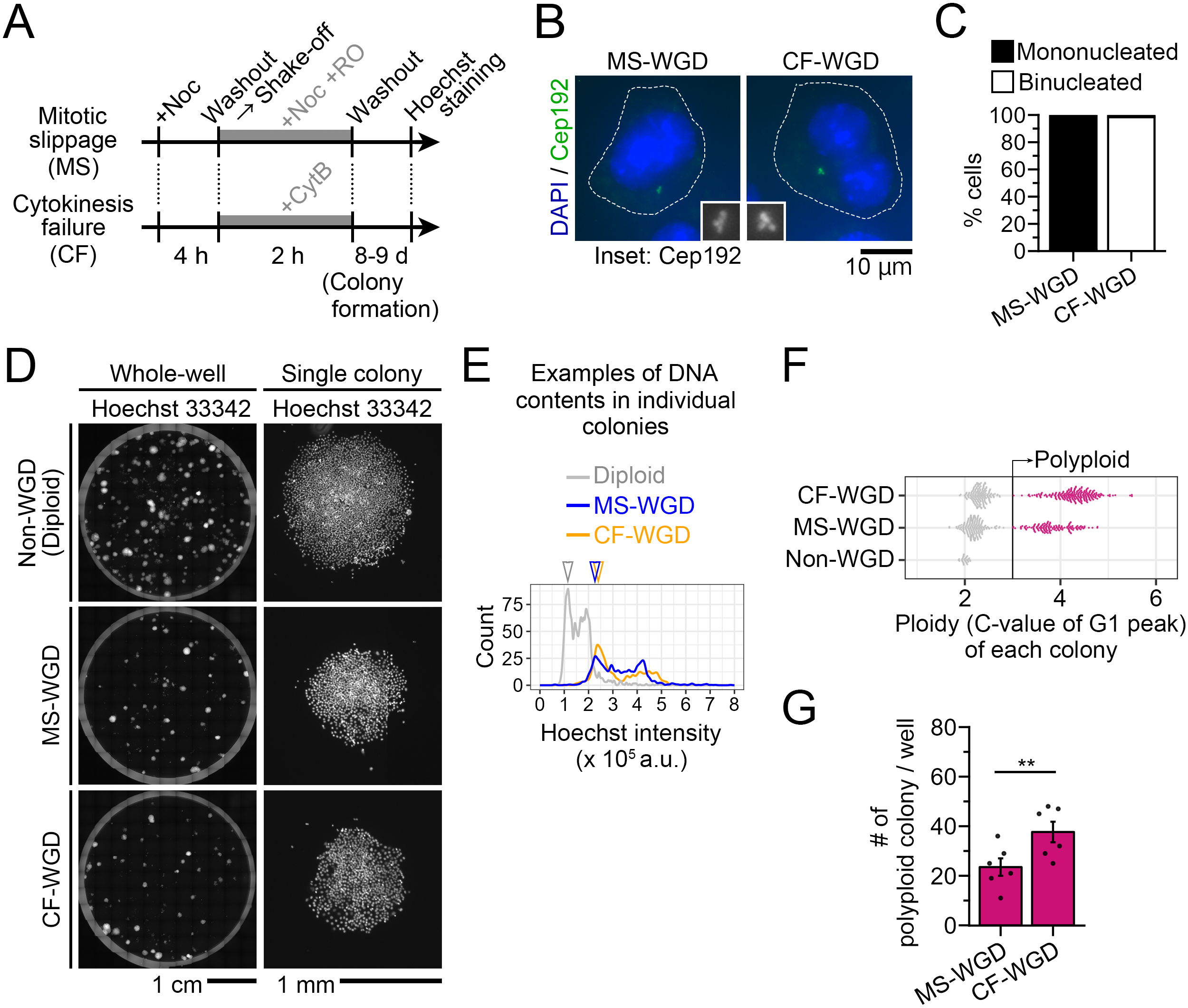
Less efficient cell proliferation after MS-WGD than CF-WGD. **(A)** A scheme of WGD induction. **(B)** An MS- or CF-WGD-induced cell, fixed at 4 h after WGD induction. The centrosomes or DNA were stained with anti-Cep192 or DAPI, respectively. Insets show 3x enlarged images of the centrosomes. Broken lines indicate cell contours. **(C)** Frequency of nuclear numbers in WGD-induced cells at the first cell cycle. Pooled data from at least 235 cells from three independent experiments. **(D)** Hoechst 33342-stained colonies formed after WGD induction. The entire well views (left) or single colony images (right). Mock treatment control without WGD induction on top (non-WGD). **(E)** Examples of DNA content distribution, quantified from Hoechst 33342 signal, in single colonies for each condition in D. Arrowheads indicate the estimated G1 DNA content values in the exemplified colonies. Note that the DNA content of G1 populations varied among colonies, reflecting their ploidy levels. See Supplemental Material S1 for histograms taken from all colonies in each condition. **(F)** Ploidy levels of individual colonies, calculated by normalizing their G1 DNA contents such that the average G1 DNA content of the diploid control was set to 2. Each data point represents a single colony. At least 355 colonies from six replicates from three independent experiments were analyzed for MS- or CF-WGD samples. Thirty-two colonies from three independent experiments for diploid control. Colonies with normalized G1 DNA content of > 3 C were defined as “polyploid.” **(G)** Number of polyploid colonies per well in each WGD condition. Means ± s.e. of six replicates from three independent experiments (\*\**p* < 0.01, the Brunner-Munzel test).

We first attempted to evaluate the survival and proliferation of post-WGD cells using flow cytometry after bulk cell culture. However, with this approach, the initially contaminated non-WGD diploid population rapidly dominated the culture within several days, precluding precise evaluation of polyploid cell growth (SI Appendix, Fig. S1D). To avoid this issue, we adopted the colony-counting assay under limited dilution after WGD induction (Fig. 1D). For this, the WGD-induced cells were seeded at 2,000 cells/well in a 6-well culture dish. Cells forming colonies after 8- or 9-d culture were stained with Hoechst 33342 to quantify DNA content distributions in individual colonies using fluorescence microscopy (Fig. 1D and E). Histograms of DNA content in most colonies showed typical cell cycle distributions with G1 and G2/M peaks, indicating their proliferative state (Fig. 1E and Supplemental material S1). Ploidy levels estimated from G1 DNA contents of individual subclones (normalized by the average DNA content in diploid control colonies) distributed in a bimodal fashion, peaking at either near-diploid or polyploid (> 3 C) (Fig. 1F). For the following reason, these near-diploid subclones were likely originated from the cells that escaped WGD induction: Based on the WGD induction rates (SI Appendix, Fig. S1C), about 50-300 out of 2,000 cells were assumed to remain diploid (i.e., escaping WGD induction) when seeded after WGD induction. When equivalent numbers of untreated diploids were seeded, we obtained colony counts roughly matching those after the aforementioned WGD induction (SI Appendix, Fig. S1E). Therefore, we excluded these near-diploid colonies and quantified the number of polyploid colonies to assess post-WGD cell proliferation (Fig. 1G). We found that CF-WGD cells reproducibly formed significantly more polyploid colonies compared to MS-WGD cells (Fig. 1G), indicating that the two WGD routes have distinct effects on subsequent cell proliferation. Since this “context-dependent” difference in post-WGD cell proliferation was novel, we decided to address the cellular processes differentially affected between MS- and CF-WGD cells.

### Poorer survivability of MS-WGD cells through multipolar chromosome segregation at the first mitosis

The significantly lower number of colonies formed after MS-WGD than CF-WGD indicates that earlier processes of cell proliferation were more severely compromised in the former. We addressed this possibility by live-imaging histone H2B-mCherry as a chromosome marker to investigate the progression and fates of subsequent interphase and mitosis after MS- or CF-WGD (Fig. 2A and B). A substantial proportion of cells completed the first interphase and entered mitosis after MS- or CF-WGD induction (81% or 80%, respectively), while the remaining cells stayed in interphase or died (SI Appendix, Fig. S2A). The relatively efficient mitotic entry indicates that the p53-mediated mechanism, which arrests the cell cycle after WGD or cytochalasin treatment (24, 27, 39), is less stringent in the HCT116 cellular background. Indeed, unlike non-cancer p53-proficient hTERT-RPE1, HCT116 failed in timely upregulating p21, the primary mediator of cell cycle arrest in the p53 pathway, despite an increase in p53 expression after WGD (SI Appendix, Fig. S2B, C, and Supporting text 1). Moreover, the mitotic stopwatch, which blocks cell cycle depending on the degree of the previous mitotic delay (40), was inactive in HCT116, resulting in no difference in first cell cycle length between MS- and CF-WGD cells (SI Appendix, Fig. S2D-I, and Supporting text 1). These data excluded the progression of the first interphase as a possible cause of the difference in proliferative capacity between MS- and CF-WGD.

**Figure 2.**
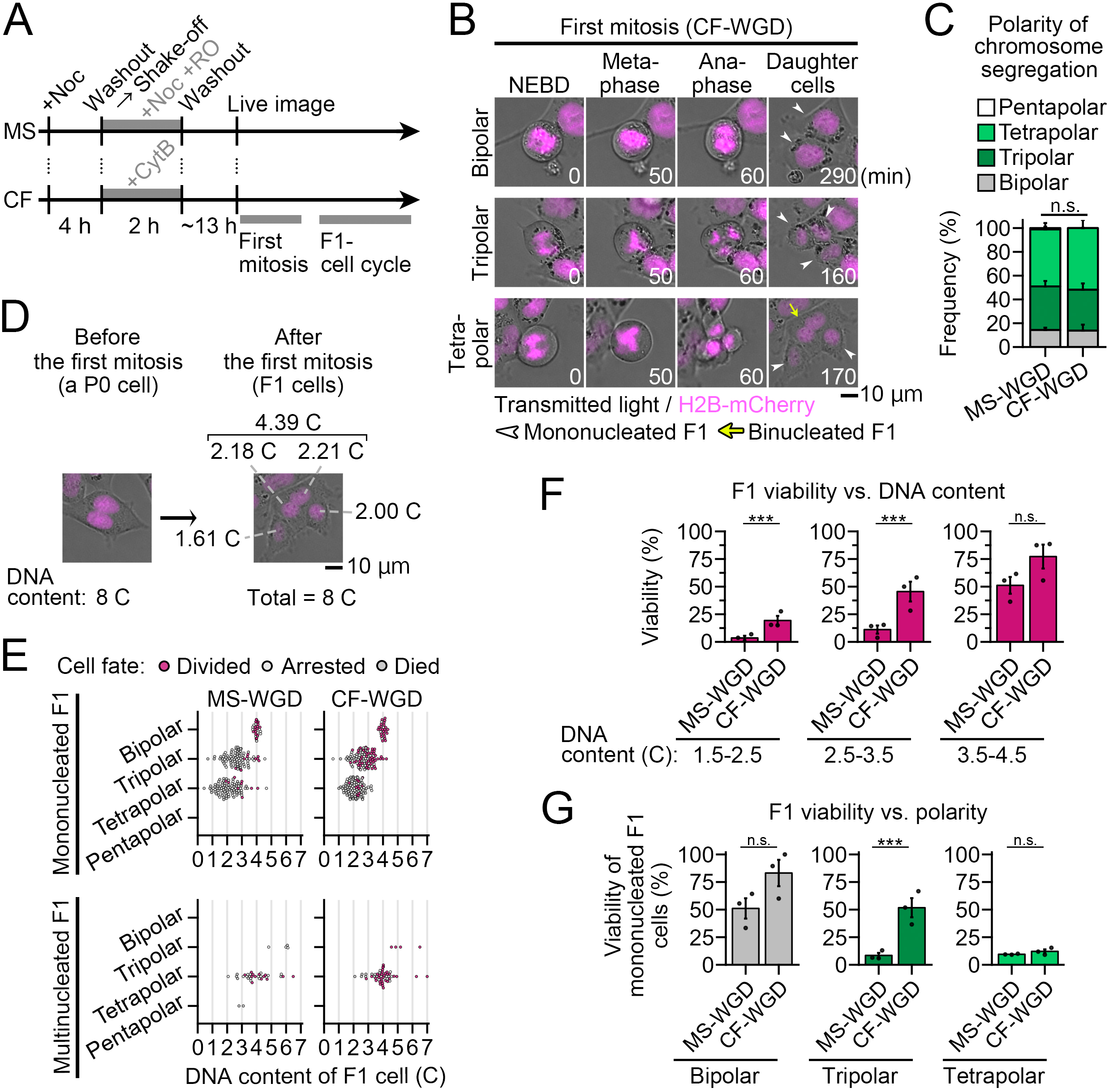
Poorer survival of MS-WGD cells through multipolar chromosome segregation at the first mitosis. **(A)** A scheme of WGD induction and live imaging. **(B)** Time-lapse of H2B-mCherry cells during the first mitosis after CF-WGD. Images were taken at 10-min intervals. The NEBD timing is set to 0 min. White arrowheads indicate “mononucleated” daughter cells. The yellow arrow indicates a binucleated cell formed through incomplete cytokinesis between two daughter cells. SI Appendix, Fig. S3B shows MS-WGD cells. **(C)** Frequency of chromosome segregation polarity in Fig. 2B and SI Appendix, Fig. S3B. Means ± s.e. of three independent experiments. At least 108 cells were analyzed in each condition (*p* = 0.95, the Fisher exact test). **(D)** An example of the estimation of daughter cell DNA content based on the segregation ratio of H2B-mCherry signal. For the demonstration, an identical cell to the “tetrapolar” cell in B is shown. **(E)** Estimated DNA content (C-value) in individual F1 cells in the live imaging. Data are sorted according to the prolife of the first mitosis (chromosome segregation polarity and the presence or absence of daughter cell multinucleation) that formed the corresponding F1 cells. At least 257 cells from three independent experiments. **(F, G)** Viability (proportion of “divided” cells) of F1 cells, sorted by DNA content of individual F1 cells (F) or by the polarity of chromosome segregation (G). Means ± s.e. of three independent experiments (n.s.: not significant, \*\*\**p* < 0.001, the Brunner-Munzel test). At least 55 or 28 cells were analyzed in each bin (F) or polarity (G), respectively. Only mononucleated F1 cells were used for the quantification in G.

We next investigated possible differences in the progression of the first mitosis. In the above live-imaging, diploids that escaped WGD induction were distinguished from WGD cells based on nuclear area immediately before mitotic entry and excluded from the analyses (SI Appendix, Fig. S3A). Because of the excess centrosomes gained upon WGD, either MS- or CF-WGD cells suffered an equivalently high frequency of multipolar chromosome segregation with no difference in mitotic duration (89% and 86%, respectively; Fig. 2B, C, SI Appendix, Fig. S3B, and C). We analyzed patterns of these multipolar chromosome segregations. At each event of the first mitosis, the nuclear H2B-mCherry signal was segregated into daughter cells (F1 cells, hereafter) at the ratio that recapitulated those measured by Hoechst 33342 (SI Appendix, Fig. S3D and E). Therefore, assuming that P0 cells had 8 C DNA content at the first mitosis after WGD, we estimated the DNA content of each F1 cell using the H2B-mCherry segregation ratio (Fig. 2D). We noticed that a substantial F1 cell proportion became multinucleated presumably because of incomplete cytokinesis (Fig. 2B, SI Appendix, Fig. S3B, and F), consistent with a previous report (36). Therefore, we separately analyzed these multinucleated F1 populations from the mononucleated ones. Reflecting the stochastic nature of multipolar division, DNA contents were heterogeneous among individual cells (Fig. 2E). DNA contents in mononucleated F1 cells from bipolar, tripolar, or tetrapolar chromosome segregation were on average 4.0, 2.7, or 2.1 C, respectively, both in MS- and CF-WGD (Fig. 2E). DNA contents in F1 cells multinucleated after tripolar or tetrapolar chromosome segregation were on average >5.0, or 3.9 C, both in MS- and CF-WGD (Fig. 2E). Therefore, either reduction in the segregation polarity or multinucleation contributed to increasing DNA contents in F1 cells.

Next, we analyzed F1 cell fates, categorized as “divided” (entered the subsequent mitosis), “arrested” (stayed alive without cell division for more than 36 h), or “died” (underwent cell death during the F1 cell cycle; Fig. 2E, SI Appendix, Fig. S3G, and H). MS-WGD produced significantly fewer “divided” F1 progenies than CF-WGD (SI Appendix, Fig. S3H). CF-WGD F1 cells gained a significantly higher chance of viability (judged by the frequency of “divided” cells) when they obtained higher DNA contents (Fig. 2F). Consistent with this trend, either the reduction in chromosome segregation polarity or daughter cell multinucleation resulted in higher F1 cell viability (Fig. 2G and SI Appendix, Fig. S3I). Whereas the viability of MS-WGD F1 cells also increased with higher DNA contents, that with 1.5-3.5 C DNA contents was significantly lower in MS-WGD than in CF-WGD (Fig. 2F). Similarly, the viability of tripolar chromosome segregation-derived MS-WGD F1 cells was significantly lower than that of CF-WGD F1 cells (Fig. 2G). This result indicates that cells are less tolerant of a reduction in DNA content following multipolar chromosome segregation after MS-WGD than after CF-WGD.

### Non-random chromosome segregation causes frequent nullisomy after MS-WGD

We next addressed the reason for the poorer viability of MS-WGD cells than CF-WGD cells after multipolar chromosome segregation at the first mitosis. For this, we compared patterns of multipolar chromosome segregation at the first mitosis by labeling specific homologous chromosomes using dCas9-EGFP with a specific single-guide RNA (sgRNA) targeting chromosome 9 pericentromere loci (C9-1) (41) (Fig. 3A). The C9-1 labeling was in good agreement with that of a chromosome 9 pericentromere-specific FISH probe (SI Appendix, Fig. S4A). We also labeled chromosomes and the centrosomes with H2B-mCherry and EGFP-GCP3, respectively (Fig. 3A and SI Appendix, Fig. S4B; C9-1 and the centrosomes were readily distinguished by their size and relative location to chromosomes).

**Figure 3.**
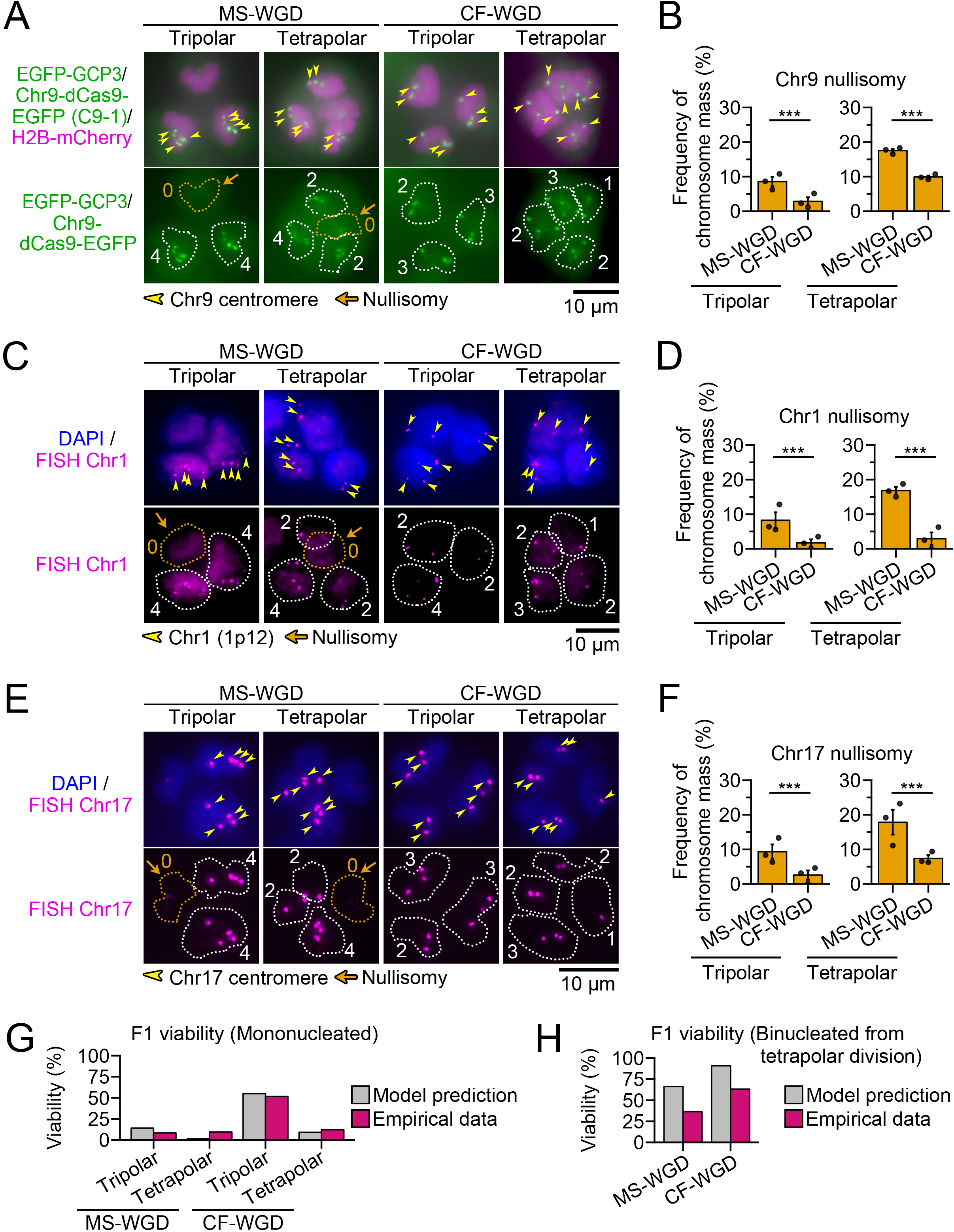
More frequent nullisomic chromosome segregation occurs at the first mitosis after MS-WGD than CF-WGD. **(A, C, E)** Anaphase cells at the first mitosis after MS- or CF-WGD. The centrosomes, centromeres of chromosome 9, and chromosomes were labeled by EGFP-GCP3, dCas9-EGFP with specific sgRNA (C9-1), and H2B-mCherry, respectively, in A. Identical images with indications of the centrosome positions are shown in SI Appendix, Fig. S4B. Genome loci on chromosome 1 (C) or chromosome 17 (E) were stained with specific FISH probes. Chromosomes were stained with DAPI in C and E. Yellow arrowheads indicate C9-1 foci or FISH signals. Broken lines indicate the contours of chromosome masses. Orange arrows indicate chromosome masses devoid of the labeled homologous chromosomes (nullisomy). The number of C9-1 foci or FISH signals in each chromosome mass is indicated in the bottom panels. **(B, D, F)** Frequency of the segregated chromosome masses devoid of any signals for chromosome 9 (B), 1 (D), or 17 (F). Means ± s.e. of three independent experiments (\*\*\**p* < 0.001, the Brunner-Munzel test). At least 252, 116, or 172 segregated chromosome masses (presumptive daughter nuclei) were analyzed in each polarity in B, D, or F, respectively. **(G, H)** Comparison between the model-predicted and empirical viability of mononucleated (G) or binucleated (H) F1 cells. See *Materials and methods* for details of the model prediction. The empirical data in G are identical to those in Fig. 2G (shown for direct comparison), and those in H were obtained from at least 33 cells from three independent experiments.

We found a prominent variation in the number of C9-1 foci per chromosome mass during multipolar chromosome segregation in both WGD conditions (SI Appendix, Fig. S4C). However, we observed nullisomic chromosome masses devoid of C9-1 focus significantly more frequently after MS-WGD than after CF-WGD (8.6% or 2.9% upon tripolar segregation and 18% or 9.9% upon tetrapolar segregation after MS- or CF-WGD, respectively; Fig. 3B and SI Appendix, Fig. S4C). Nullisomic chromosome masses were also more frequent after MS-WGD than CF-WGD, when chromosome 1 or 17 was labeled by FISH (Fig. 3C-F). Moreover, we observed a similar extent of increase in nullisomic segregation when MS-WGD was induced by the transient co-inhibition of a mitotic kinesin, Eg5, and Mps1 kinase (SI Appendix, Fig. S4D; to suppress SAC with monopolar spindle formation). A similar trend of MS-WGD-linked increase in nullisomic segregation was also observed in hTERT-RPE1 cells (SI Appendix, Fig. S4E-G; p53 was depleted by RNAi to allow entry into the first mitosis after WGD). Therefore, MS-WGD led to more frequent nullisomic segregation regardless of chromosome species, the means of MS induction, or the cellular backgrounds we tested.

Nullisomy of any chromosome species most likely results in death in the F1 generation. Therefore, we theoretically estimated F1 cell viability as the frequency of cells with at least one copy of all chromosome species (Fig. 3G and H). In this estimation, we assumed that the probability of nullisomy formation for all chromosome species was equivalent to that for chromosome 9 (Fig. 3B; see *Discussion* on limitations of the simplified assumption). We estimated F1 viability in either situation i) where the observed chromosome mass was segregated into a mononucleated daughter (Fig. 3G), or ii) where two masses were segregated into a binucleated daughter (Fig. 3H; see Materials and methods for details). The estimated viability of mononucleated F1 cells was in good agreement with the empirical value in the live-imaging analysis for both MS-WGD and CF-WGD (Fig. 3G). The theoretical estimation also recapitulated the drastic increase in F1 cell viability upon their binucleation, though it tended to overestimate the trend (Fig. 3H). These results suggest that the proneness of MS-WGD-derived F1 cells to nullisomy, at least in part, explains their poorer survivability after multipolar chromosome segregation. In a previous theoretical estimation in human WGD cells, 75% or 23% of F1 cells avoided nullisomy of any homologs upon tripolar or tetrapolar segregation, respectively, assuming that all chromosomes were randomly segregated (32). The estimated values were closer to those observed after CF-WGD (Fig. 3G), indicating that the homologous chromosome segregation occurred in a non-random manner after MS-WGD.

### MS-WGD causes skewed homologous chromosome distribution at the entry into the first mitosis

We next addressed why nullisomic chromosome segregation occurred more frequently at the first mitosis after MS-WGD than CF-WGD. MS- and CF-WGD cells showed lagging chromosomes or chromosome bridges to the same extent (SI Appendix, Fig. S4H-M), ruling out differences in chromosome segregation fidelity as the cause of the difference in nullisomy frequency between these conditions. As another possible factor affecting nullisomy frequency, we next compared the spatial distribution of homologous chromosomes upon the entry into the first mitosis between MS- and CF-WGD cells by imaging C9-1 along with chromosomes, and the centrosomes (Fig. 4A). At this mitotic stage, most WGD cells possessed four C9-1 foci, as sister chromatids had not yet been separated. In CF-WGD cells, two C9-1 foci in each of two nuclei tended to be located distal to each other, resulting in relatively even distributions of the four C9-1 foci (with an average inter-centromere distance of 9.1 μm; Fig. 4A, B, and SI Appendix, Fig. S5A). In sharp contrast, in MS-WGD cells, four C9-1 foci from the single nucleus were often distributed pairwise (Fig. 4A). Accordingly, the frequency of “paired C9-1 foci,” defined as those with the inter-centromere distance of < 2 μm, was significantly higher after MS-WGD than CF-WGD (Fig. 4B, C, SI Appendix, Fig. S5A, and B). These C9-1 pairs were occasionally located proximally from each other, resulting in an overall shorter inter-centromere distance among all four C9-1 foci in MS-WGD cells (on average 6.1 μm; Fig. 4B and SI Appendix, Fig. S5A). We also quantified the angles between each two C9-1 foci with regard to the centroid of the whole-chromosome mass (Fig. 4D). The angles were significantly smaller in MS-WGD than in CF-WGD cells (Fig. 4D), reflecting more skewed configurations of homologous centromeres in the former condition.

**Figure 4.**
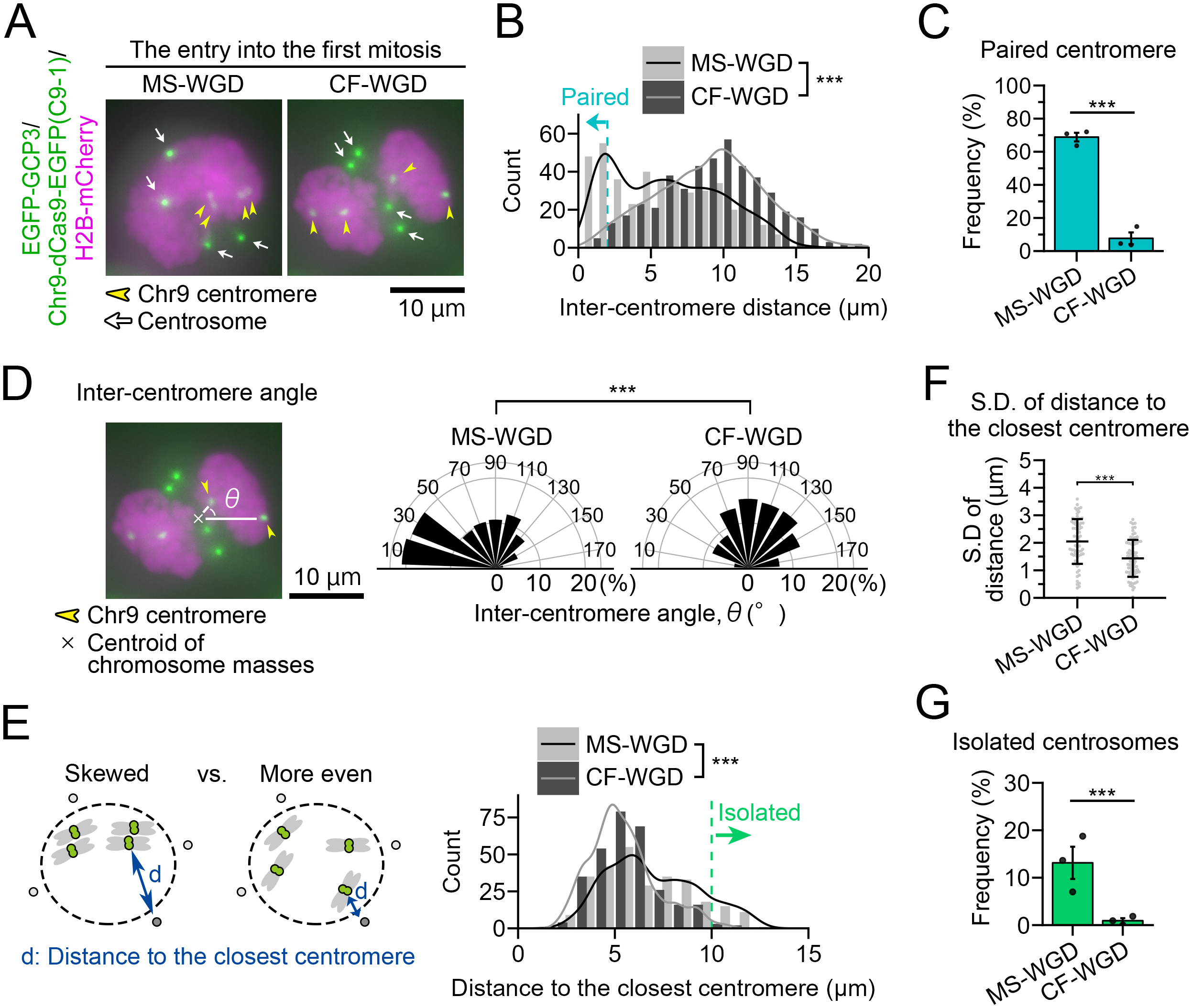
Skewed distribution of homologous centromeres in the first mitosis after MS-WGD. **(A)** Cells at the entry into the first mitosis after MS- or CF-WGD. The centrosomes, centromeres of chromosome 9, and chromosomes were labeled by EGFP-GCP3, dCas9-EGFP with specific sgRNA (C9-1), and H2B-mCherry, respectively. **(B)** Histogram of inter-centromere distance in A. Six inter-centromere distances among the 4 C9-1 foci were obtained from each cell. At least 414 C9-1 centromere combinations from at least 69 cells from three independent experiments (\*\*\**p* < 0.001, the Welch’s t-test). C9-1 combinations with their distance less than 2 μm (defined as “paired centromeres”) are indicated by the blue arrow. **(C)** Frequency of cells with paired C9-1 centromeres in A. Means ± s.e. of three independent experiments (\*\*\**p* < 0.001, the Brunner-Munzel test). **(D)** Left: An example of the inter-centromere angle quantification. For the demonstration, the identical CF-WGD cell in A is shown. Right: Histogram of the angles between two C9-1 foci with regard to the centroid of the whole-chromosome mass in A. At least 354 C9-1 combinations from at least 59 cells from three independent experiments were analyzed (\*\*\**p* < 0.001, the Welch’s t-test). **(E)** Left: Schematic images of the closest centrosome-centromere distance with skewed or more even homologous centromere distributions. Right: Histogram of the distance from each centrosome to its closest C9-1 focus in A. Four centrosome-centromere distances were obtained from an individual cell. The centrosome-centromere combinations whose distances exceeded 10 μm (defined as “isolated centrosomes”) are indicated by the green arrow. At least 276 centrosome-centromere combinations from at least 69 cells from three independent experiments were analyzed (\*\*\**p* < 0.001, the Welch’s t-test). **(F)** Standard deviation of the closest centrosome-centromere distances in each cell in A. Asterisks indicate statistically significant difference between conditions (\*\*\**p* < 0.001, Welch’s t-test). **(G)** Frequency of isolated centrosomes in A. Means ± s.e. of three independent experiments (\*\*\**p* < 0.001, the Brunner-Munzel test).

Using the above images, we also analyzed spatial coordination between each C9-1 foci and the centrosomes. Four centrosomes had already been separated in most MS- and CF-WGD cells around NEBD, often flanking the chromosome masses from their opposite sides (Fig. 4A and SI Appendix, Fig. S5C). Reflecting the difference in the skewedness of C9-1 distribution, the distance from each centrosome to the closest C9-1 focus was significantly larger with higher deviation in MS-WGD than in CF-WGD cells (7.0 μm or 5.4 μm in MS- or CF-WGD cells, respectively; Fig. 4E, F, and SI Appendix, Fig. S5D). Accordingly, we observed “isolated centrosomes” located extremely far (> 10 μm) from any of C9-1 foci more frequently in MS-WGD than in CF-WGD cells (Fig. 4E and G). We also observed the skewed centromere distribution with the frequent centrosome isolation in MS-WGD induced by Eg5 and Mps1 co-inhibition (SI Appendix, Fig. S5A-F). These data collectively demonstrate that MS-WGD and CF-WGD differentially impact the geometry of the first mitosis, with the former introducing more skewed homologous chromosome distributions relative to the centrosomes.

### Geometric isolation of centrosomes from all homologues drives nullisomic chromosome segregation after MS-WGD

Distally placed centrosomes may be less likely to capture any homologous centromeres, possibly explaining the frequent nullisomy after MS-WGD. To test this idea, we traced C9-1 foci and the centrosomes from entry into the first mitosis through anaphase after MS-WGD using live imaging (Fig. 5A and SI Appendix, Movie S1). Following the centrosome separation starting before NEBD (SI Appendix, Fig. S6A), spatial coordination between C9-1 foci and the centrosomes drastically changed around NEBD (Fig. 5A). As a result, the distance between each C9-1 focus and each centrosome became highly deviated around NEBD (SI Appendix, Fig. S6B), which was consistent with the above observation (Fig. 4E, F, and SI Appendix, Fig. S5D). During chromosome congression at prometaphase, the initial positions of C9-1 foci around the NEBD were well transmitted to their positions within the aligned metaphase plate (Fig. 5A and SI Appendix, Fig. S6C), consistent with a previous observation in normal mitosis (42). At anaphase onset, each C9-1 focus split into two sister foci. Each of these sister foci was segregated toward one of the four centrosomes, enabling us to identify the centrosomes that captured individual sister chromatids in the early mitotic phase (Fig. 5A and SI Appendix, Fig. S6B). Using this information, we compared distances from C9-1 foci to the centrosomes that captured these foci with those to the centrosomes that did not capture these foci (Fig. 5B). The former were significantly shorter than the latter from NEBD to the late prometaphase (Fig. 5B). We also analyzed relationship between the C9-1-centrosome distance at NEBD with the frequency of the subsequent capturing of the C9-1 focus by the centrosome (Fig. 5C). A centrosome captured a C9-1 focus at the frequency of >75% when their distance was ≤ 10 μm at NEBD, while the frequency became less than 50% when the distance was > 10 μm (Fig. 5C). Therefore, the initial proximity of the centrosomes to each centromere was linked to a higher chance of their capturing, potentially determining the pattern of subsequent homologue segregation.

**Figure 5.**
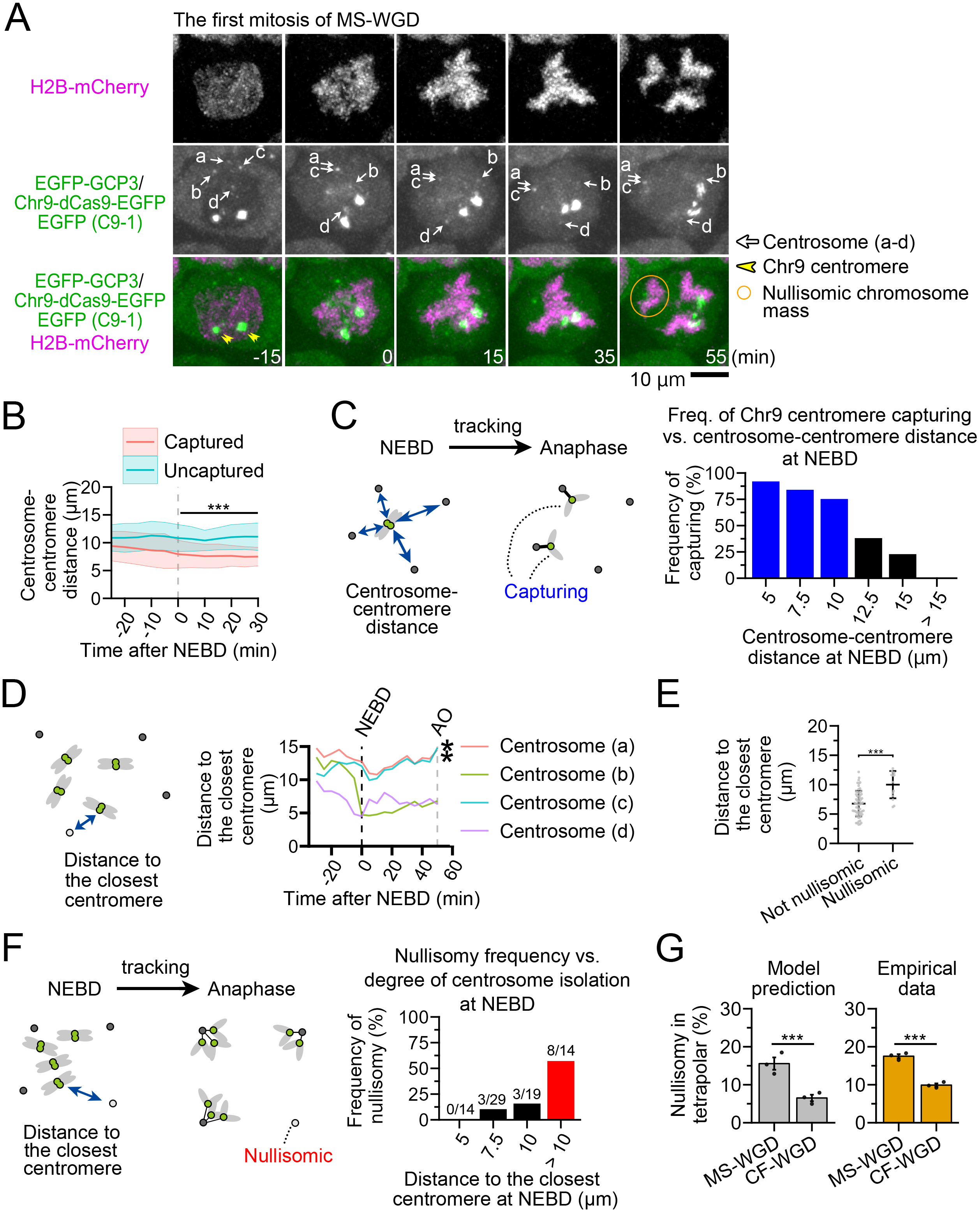
Geometric isolation of centrosomes drives nullisomic chromosome segregation after MS-WGD. **(A)** Time-lapse images of H2B-mCherry, EGFP-GCP3, and dCas9-EGFP with C9-1 sgRNA throughout the first mitosis after MS-WGD. Images were taken at 5-min intervals. The NEBD timing is set to 0 min. The yellow circle indicates a chromosome mass devoid of any C9-1 foci. Individual centrosomes were distinguished by labeling different letters (a-d). **(B)** The centrosome-centromere distance of captured or uncaptured pairs in A. Means ± s.d. of at least 114 centrosome-centromere pairs from 19 cells from five independent experiments. Asterisks indicate statistically significant differences between conditions throughout mitosis (0-30 min after NEBD; \*\*\**p* < 0.001, the Welch’s t-test). **(C)** The frequency of centrosome-centromere capturing, which was calculated for each bin of the initial centrosome-centromere distance (schematized on the left). Three hundred four centrosome-centromere pairs from 19 cells from five independent experiments were analyzed. **(D)** The time course of the distance from each centrosome to its closest C9-1 focus (schematized on left) in the cell shown in A. The asterisk indicates a centrosome that failed to capture any of the C9-1 foci. **(E)** The initial distance (at NEBD) to the closest C9-1 focus from centrosomes that captured at least one C9-1 focus (non-nullisomic) or those that did not capture any foci (nullisomic). Means ± s.d. of at least 14 centrosomes from 12 cells from five independent experiments (\*\*\**p* < 0.001, the Welch’s t-test). **(F)** The frequency of the centrosomes that failed to capture any C9-1, which was calculated for each bin of the closest centrosome-centromere distance at NEBD (schematized on the left). At least 76 centrosomes from 19 cells from five independent experiments were analyzed. The number of centrosomes failing to capture any C9-1 per total centrosomes in each bin is shown on top of each graph bar. **(G)** Comparison between the model-predicted and empirical frequency of segregated chromosome masses devoid of any C9-1 foci in tetrapolar chromosome segregation at the first mitosis after MS- or CF-WGD. See *Materials and methods* for details of the model prediction. The empirical data are identical to those in Fig. 3B (for the direct comparison). Means ± s.e. of three independent experiments (\*\*\**p* < 0.001, the Brunner-Munzel test).

To gain further insight into the possible relationship between early centrosome-centromere arrangement and the subsequent formation of nullisomy, we analyzed the time course of the distance from each centrosome to its closest C9-1 focus (Fig. 5D). The centrosomes that did not capture any C9-1 foci located significantly more distant from their closest C9-1 focus (on average 10 µm, at NEBD) than those capturing at least one C9-1 focus (6.8 μm; Fig. 5D and E). The frequency for a centrosome to fail to capture any C9-1 foci (nullisomic segregation) drastically increased when the distance from the centrosome to the closest C9-1 focus was more than 10 μm at NEBD (Fig. 5F). The same trend was also observed in a live imaging analysis of CF-WGD cells (SI Appendix, Fig. S6D). Consistent with these quantitative data, visual comparisons of C9-1 and centrosome locations between NEBD and anaphase showed the trend that the centrosomes initially located at the area devoid of C9-1 focus failed to capture any C9-1 foci at the subsequent anaphase (Fig. 5A and SI Appendix, Fig. S6C). These results suggest that the initial centrosome-centromere distance is a key determinant of subsequent nullisomy formation.

We reasoned that, if the above assumption was correct, the frequency of nullisomic homologue segregation could be well predicted by the centrosome-centromere distance around NEBD (Fig. 5F). Indeed, the frequency of the nullisomic homologue segregation, which was predicted based on the distance between each centrosome and its closest C9-1 focus around NEBD, well matched with those obtained by experiments either in MS- or CF-WGD (Fig. 5G, see also Fig. 3B; Materials and methods for details). These data collectively support the idea that the proneness of MS-WGD cells to nullisomic homologue segregation is mainly attributed to the skewed homologue distribution at the entry into the first mitosis.

### Forced sister chromatid separation in the MS-WGD time window suppresses subsequent nullisomic chromosome segregation

We wished to further address the causality between the skewed homologue distribution and the frequent nullisomic homologue segregation after MS-WGD. To investigate this, we traced C9-1 foci during MS-WGD to determine how homologue distribution became skewed (SI Appendix, Fig. S7A). Live-imaging revealed that, while < 22% (6 out of 28) sister foci separated by a short distance (> 2 μm) during MS-WGD, the rest (22 out of 28) failed to separate to a detectable degree (SI Appendix, Fig. S7A and B). As a result, the distance between sister C9-1 foci remained on average 1.3 μm at the time the cells exited mitosis (SI Appendix, Fig. S7C).

Inefficient chromatid separation during MS-WGD may skew homologue distribution and limit the evenness of their capturing during the subsequent mitosis. To directly test this idea, we attempted to induce precocious sister chromatid separation by depleting RAD21, a cohesin subunit, during MS-WGD. To avoid pleiotropic damage caused by long-term loss-of-function of cohesin, we employed the auxin-inducible degron 2 (AID2) technology to deplete RAD21 only for a short time window prior to MS-WGD (Fig. 6A) (43). We treated 5-Ph-IAA in RAD21-mAID-Clover (RAD21-mAC) HCT116 expressing OsTIR1(F74G) (43) for 2 h (Fig. 6A), which efficiently degraded RAD21-mAC by the time of MS-WGD induction (SI Appendix, Fig. S7D-F). RAD21-mAC degradation induced precocious sister chromatid disjunction with a significant increase in inter-centromere distance during prometaphase arrest and caused further separation during MS-WGD (Fig. 6B, C, SI Appendix, Fig. S7G-J). As a result, distances between sister C9-1 foci became significantly longer after mitotic exit in RAD21-degraded cells compared to mock-treated control (on average 2.8 μm or 1.5 μm in 5-Ph-IAA- or DMSO-treated condition, respectively; Fig. 6B and C).

**Figure 6.**
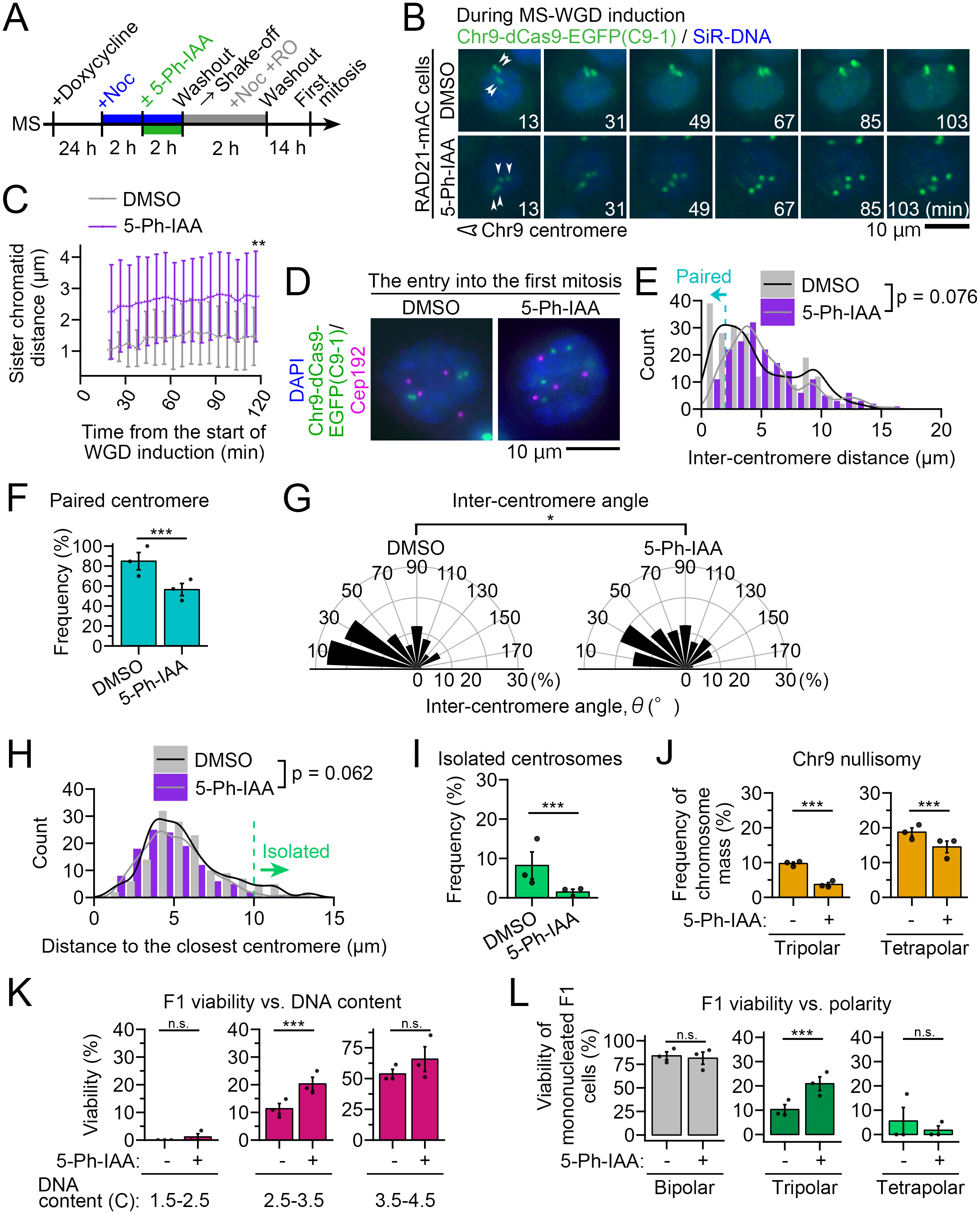
Suppression of nullisomic homologue segregation at the first mitosis by artificial sister chromatid separation upon MS-WGD. **(A)** The schedule for AID2-mediated transient depletion of RAD21 before MS-WGD induction. **(B)** Time-lapse of dCas9-EGFP with C9-1 sgRNA and chromosomes stained with SiR-DNA during MS-WGD in RAD21-mAC cells treated with or without 5-Ph-IAA. Images were taken at 6-min intervals. The time after the start of WGD induction was indicated. **(C)** Time course of inter-sister chromatid distance in B. Means ± s.d. of at least 22 sister chromatid pairs from at least 11 cells from two independent experiments (\*\**p* < 0.01, the Welch’s t-test; tested at the last time point). **(D)** Cells at the entry into the first mitosis after MS-WGD in RAD21-mAC cells treated with or without 5-Ph-IAA. The centrosomes, C9-1 centromeres, and chromosomes were labeled by immunostaining of Cep192, expression of dCas9-EGFP with a targeting sgRNA, and DAPI staining, respectively. **(E, H)** Histogram of inter-centromere distance (E) or the distance from each centrosome to its closest C9-1 focus (H). At least 168 C9-1 combinations (E) or at least 112 centrosome-C9-1 combinations (H) from at least 28 cells from three independent experiments were analyzed (*p* values of the Welch’s t-test are shown). **(F, I)** Frequency of cells with paired C9-1 centromeres (F) or frequency of isolated centrosomes (I) in D. Means ± s.e. of three independent experiments (\*\*\**p* < 0.001, the Brunner-Munzel test). **(G)** Angles between each two C9-1 foci with regard to the centroid of the whole-chromosome mass. At least 168 C9-1 combinations from at least 28 cells from three independent experiments were analyzed (\**p* < 0.05, the Welch’s t-test). **(J)** Frequency of C9-1 nullisomic chromosome segregation during anaphase at the first mitosis after MS-WGD with or without the transient RAD21-mAC depletion. Means ± s.e. of three independent experiments (\*\*\**p* < 0.001, the Brunner-Munzel test). At least 150 segregated chromosome masses from at least 50 cells from three independent experiments were analyzed in each polarity. **(K, L)** Viability of F1 cells in live imaging analysis after MS-WGD with or without the transient RAD21 depletion. Data are sorted by ploidy of individual F1 cells (K) or the polarity of chromosome segregation at the first mitosis (L). Means ± s.e. of three independent experiments (n.s.: not significant, \*\*\**p* < 0.001, the Brunner-Munzel test). At least 59 or 30 cells were analyzed in each bin (K) or polarity (L), respectively. Only mononucleated F1 cells were used for the quantification in L.

We then removed 5-Ph-IAA, allowing complete restoration of RAD21-mAC expression in the subsequent cell cycle (SI Appendix, Fig. S7E and F). Chromosome spreading showed complete restoration of sister chromatid cohesion at the first mitosis after removing 5-Ph-IAA (SI Appendix, Fig. S7I and J), demonstrating that precocious sister chromatid disjunction was induced only before MS-WGD. Additionally, “diplochromosomes,” which form when the sister chromatids do not separate during the previous cell cycle (44), were not prevalent, even in the mock-treated control (SI Appendix, Fig. S7I and Supplemental material S2).

We next tested the effect of the transient RAD21-mAC depletion before MS-WGD on the subsequent homologue distribution at the first mitosis (Fig. 6D). Along with C9-1, chromosomes and the centrosomes were visualized by DAPI and anti-Cep192 immunostaining, respectively. Distance among C9-1 foci tended to be longer with a significant reduction in “paired C9-1 foci” by the transient RAD21-mAC depletion (Fig. 6D-F). The transient RAD21-mAC depletion also significantly increased the inter-C9-1 foci angles, indicating dispersion of homologous centromeres (Fig. 6G). The distance from each centrosome to its closest C9-1 focus tended to be shorter, with a significant reduction in “isolated centrosomes” by the transient RAD21-mAC depletion (Fig. 6H and I). Moreover, the transient RAD21-mAC depletion significantly reduced the frequency of nullisomic segregation of C9-1 foci, both in tripolar and tetrapolar chromosome segregation (Fig. 6J).

We also induced precocious sister chromatid separation during MS-WGD by RNAi-mediated depletion of Sororin, a cohesin-stabilizing factor (SI Appendix, Fig. S8A-C) (45, 46). Similar to RAD21-mAC depletion, precocious sister chromatid separation by Sororin depletion led to substantial dispersion of C9-1 foci compared to mock-depleted control at the first mitosis (SI Appendix, Fig. S8D-I), with a significant reduction in the frequency of nullisomic segregation in the subsequent anaphase (SI Appendix, Fig. S8J). The above data collectively demonstrate that incomplete sister chromatid separation during MS-WGD is the primary cause of uneven distribution of homologous chromosomes after WGD, facilitating the nullisomy-prone multipolar chromosome segregation in the subsequent mitosis.

### Forced sister chromatid separation in the MS-WGD time window improved the viability of the F1 generation

Finally, we determined the causality between inefficient sister chromatid separation during MS-WGD and poorer cell survival in the subsequent mitosis. For this, we tested the effects of precocious sister chromatid separation during MS-WGD on subsequent F1 cell viability. Although we initially tried RNAi-mediated Sororin depletion, it resulted in frequent cell cycle arrest before entry into the first mitosis (SI Appendix, Fig. S8K). This result presumably reflects the deleterious effects of consecutive Sororin depletion on the progression of the first cell cycle after MS-WGD, precluding us from using this condition to assess F1 cell viability. Therefore, we next tried transient RAD21-mAC depletion, which avoided the consecutive loss of function of cohesin (SI Appendix, Fig. S7E, F, I, and J). We compared F1 cell viability after MS-WGD with or without the transient RAD21-mAC depletion by live imaging. The transient RAD21-mAC depletion did not affect the efficiency of entry into the first mitosis (SI Appendix, Fig. S9A), suggesting minimal damage during the cell cycle after MS-WGD. The polarity of chromosome segregation at the first mitosis was also unaffected by the transient RAD21-mAC depletion (SI Appendix, Fig. S9B). However, compared to the mock-depleted MS-WGD control, the transient RAD21-mAC depletion significantly increased viable F1 cells with DNA content of 2.5-3.5 C or those generated through tripolar chromosome segregation (Fig. 6K, L, and SI Appendix, Fig. S9C). Total viability of F1 cells was also significantly improved in the transient RAD21-mAC-depleted sample than in the mock-depleted control (SI Appendix, Fig. S9D). These results demonstrate that inefficient sister chromatid separation during MS-WGD limits the viability of their progeny, at least in part, through skewing the distribution of chromosome territories and increasing the chance of nullisomic homologue segregation in the subsequent multipolar cell division.

## Discussion

While WGD occurs through multiple cellular processes, it has been unclear whether differences in the WGD route affect the characteristics of the post-WGD cell. We found that the efficiency of sister chromatid separation during WGD defined the spatial arrangement of homologous chromosomes in the subsequent cell cycle. Homologous chromosome arrangement then determined the patterns of multipolar chromosome segregation at the first mitosis, critically affecting the viability of resulting daughter cells. Our findings demonstrate that differences in geometric properties between MS- and CF-WGD cells have a crucial influence on the proliferative characteristics of their progeny.

We propose the following mechanism of the route-dependent difference in post-WGD cell viability (see SI Appendix, Fig. S9E for a model). Due to the inefficient sister chromatid separation, MS-WGD causes more skewed homologue distribution in the subsequent interphase than CF-WGD, often with all four homologous chromosomes located on one side of the nucleus (SI Appendix, Fig. S9E-i). The skewed homologue distribution persists until the entry into the first mitosis, when the separated four centrosomes surround the chromosome mass from all sides (SI Appendix, Fig. S9E-ii). This intracellular configuration causes a higher deviation in the centrosome-homologous centromere distance after MS-WGD than CF-WGD, with some centrosomes located particularly distal from any of the homologous centromeres. Our live-imaging suggests that this heterogeneity in the distance from each centrosome to the homologous centromeres has a crucial impact on the efficiency of its homologue capturing: A centrosome placed further than 10 μm from any homologous centromeres at NEBD has a drastically higher chance of nullisomic segregation (Fig. 5F and SI Appendix, Fig. S6D). Interestingly, this threshold distance corresponds well with the length that spindle microtubules can reach during prometaphase in mammalian cells (47). Therefore, a centrosome is more likely to lose to other centrosomes in the “competition” for homologous chromosomes when it is initially isolated from homologous centromeres further than its microtubules can reach (SI Appendix, Fig. S9E-iii). We found that the degree of the initial centrosomal isolation well explained the frequency of subsequent nullisomic segregation (Fig. 5F and G). Moreover, the artificial sister chromatid separation in MS-WGD significantly alleviated the skewed chromosome distribution and reduced nullisomic segregation (Fig. 6 and SI Appendix, Fig. S8). These results demonstrate that the skewed homologous chromosome arrangement resulting from inefficient sister chromatid separation is the crucial cause of the higher prevalence of nullisomic chromosome segregation after MS-WGD (SI Appendix, Fig. S9E-iv). In other words, sister chromatid separation contributes to “evening out” the intranuclear distribution of homologous chromosomes, facilitating efficient centrosome-chromosome interactions at mitosis in the next cell cycle, whose significance becomes evident in the geometric condition of post-WGD cells.

The low prevalence of diplochromosomes in the first mitosis after MS-WGD (SI Appendix, Fig. S7I) suggests that sister chromatids are mostly disjoined upon MS-WGD, as reported in other animal cells (48). However, live imaging revealed that they failed to separate efficiently by a distance (Fig. 6C and SI Appendix, Fig. S7C), in contrast to more efficient chromosome separation in the absence of a spindle (so-called “C-anaphase”) in plant cells (49). We found that precocious sister chromatid disjunction induced their separation even during prometaphase, resulting in a further increase in their distance during the subsequent MS-WGD (SI Appendix, Fig. S7H and 6C). Although the mechanisms underlying the pre-anaphase separation remain unknown, we speculate that this precocious separation helps further chromatid segregation observed during MS-WGD in RAD21- or Sororin-depleted cells.

While our results suggest that the frequent nullisomy caused by the skewed homologous chromosome distribution is an important factor that limits the F1 cell viability after MS-WGD, the following facts indicate the contributions of other unknown factors to limiting viability of MS-WGD cells: I) While the transient RAD21-mAC depletion reduced the frequency of nullisomy after MS-WGD to the equivalent level as that after CF-WGD (particularly in case of tripolar segregation; Fig. 6J), the F1 viability after MS-WGD with the transient RAD21-mAC depletion was still lower than that after CF-WGD (Fig. 6K, L, 2F, and G). II) Moreover, CF-WGD progenies showed better survivability than MS-WGD progenies even after bipolar chromosome segregation (Fig. 2G; supposed to be devoid of frequent nullisomy). In this context, it would be beneficial to consider other issues arising from MS-WGD beyond the aggravation of nullisomy. A recent study in C. elegans embryos reported that artificially altering the developmentally programmed CF-WGD in intestinal cells to MS-WGD compromised the expression of genes required for embryo growth (50). The study proposed that differences in nuclear surface-to-volume ratio may have altered gene regulation between MS- and CF-WGD. Another recent study reported MS-specific alterations in chromatin organization, attributed to precocious mitotic exit and an abnormally high level of mitotic-specific histone H3 phosphorylation (51). Therefore, several independent principles, including the one found here, may critically affect the more dysregulated characteristics of MS-derived cells.

Fates of post-WGD progenies beyond the F1 generation would be more complicated. We found that the F1 population, whether from MS- or CF-WGD, included mononucleated and multinucleated polyploids (Fig. 2E). Therefore, their intranuclear organization is expected to be more heterogeneous and potentially cluttered than that immediately after WGD. Therefore, WGD route differences would have direct effects only on the intranuclear organization and chromosome segregation patterns in the first cell cycle. However, F1 cells derived from MS- and CF-WGD cells differed not only in the frequency of nullisomy, but also in that of other types of copy number abnormalities, such as monosomy or trisomy (SI Appendix, Fig. S4C). Therefore, the initial differences between MS- and CF-WGD cells would be transmitted and have a long-term influence on the characteristics of their progeny in the form of basal karyotypic profiles. Related to this point, it should be noted that the frequency of copy number alterations may differ across chromosome species, whereas at least three chromosome species (chromosomes 1, 9, and 17) with different sizes showed a similar trend in our results. To accurately understand genome-wide trends in karyotypic alterations across different WGD routes, single-cell genome-wide copy-number analysis would be a powerful approach in future studies.

The impacts of differences in WGD routes on subsequent cell viability may also provide an important clue to the control of post-WGD cell populations in cancer therapeutics. Recently, it has been pointed out that anticancer drug treatment often causes artificial induction of WGD in tumors, potentially causing the risk of recurrence through subsequent proliferation of these post-WGD progenies (2). Since sister chromatid separation is a key factor affecting post-WGD cell proliferation, suppressing it, for example, through topoisomerase II inhibitors (52) could be a potential approach to reduce the risk of post-WGD cell proliferation. Our findings indicate the importance of understanding the differential effects of WGD processes on subsequent cell fates for accurate comprehension of principles of WGD-driven biological processes and their precise control.

## Materials and methods

A detailed description of Materials and methods is provided in Supporting Text 2 in the SI Appendix.

### Cell culture and reagents

Cell lines, plasmid vectors, or compounds and antibodies used in this study are listed in Dataset S1, S2, or S3, respectively. Cell culture, nucleic acid transfection, WGD induction, or biochemical analyses were conducted as detailed in SI Appendix, Supporting Text 2.

### Imaging

All cell images were acquired using a TE2000, Ti, or Ti2 microscope (Nikon). Detailed information on sample preparation, microscope equipment, or quantitative image analysis is provided in SI Appendix, Supporting Text 2.

### Statistical analysis

Analyses for significant differences between or among conditions were conducted as detailed in SI Appendix, Supporting Text 2. Statistical significance was set at *p* < 0.05. *P*-values are indicated in figures or the corresponding figure legends.

## Acknowledgment

We are grateful to Masato Kanemaki, Gohta Goshima, and Kinya Yoda for reagents, Sarah McClelland, Koya Yoshizawa, Saori Takahashi and Ichiro Hiratani for valuable comments, and the NIC at Hokkaido University for microscopes. MI and GY are supported by JST SPRING, Grant Number JPMJSP2119. This work was supported by JSPS KAKENHI (Grant Numbers JP19KK0181, JP19H05413, JP19H03219, JPJSBP120193801, JP21K19244, JP22H04926, JP24K21956, JP24KK0139, and JP24K02017 to R.U.), the Princess Takamatsu Cancer Research Fund, the Kato Memorial Bioscience Foundation, the Orange Foundation, the Smoking Research Foundation, Daiichi Sankyo Foundation of Life Science, the Akiyama Life Science Foundation, the Hoansha Foundation, Sumitomo Electric Group CSR Foundation, and the Terumo Life Science Foundation to R.U. The authors declare no competing financial interests.

## Author Contributions

Conceptualization, M.I., and R.U.; Methodology, M.I., G.Y., Y.T., and R.U.; Investigation, M.I.; Formal Analysis, M.I., and R.U.; Resources, M.I., G.Y., Y.T., and R.U.; Supervision, R.U.; Writing – Original Draft, R.U.; Writing – Review & Editing, M.I., and R.U.; Funding Acquisition, M.I., G.Y., Y.T., and R.U.

