## Supplementary material for "Sister chromatid separation determines the proliferative properties upon whole-genome duplication via homologous chromosome arrangement": SI appendix: SI251223final_260218_BioRxiv.pdf

**This PDF file includes:**

Supporting text
Figures S1 to S10
Legends for Movie S1
Legends for Supplemental materials S1 to S3
SI References

**Other supporting materials for this manuscript include the following:**

Dataset S1 to S4
Movies S1
Supplemental materials S1 to S3

#### Supporting Information Text

##### Supporting text 1: Assessment of mitotic defect-responding signaling in HCT116 cells

Previous studies using non-cancer p53-proficient cell lines reported p53-dependent cell cycle arrest after WGD induction or cytochalasin treatment (1-3). In contrast, although HCT116 cells had wild-type p53, they frequently progressed through the cell cycle and entered mitosis after WGD induction in our live-imaging analysis (Fig. S2A). This observation indicates that p53-dependent cell cycle regulation under these perturbations is not stringent in HCT116 cells. To address this possibility, we compared p53 signaling in HCT116 and hTERT-RPE1 (a p53-proficient control cell line) at 4 or 15 h after release from mitotic synchronization, with or without subsequent WGD induction (Fig. S2B and C). To assess p53 signaling status, we quantified the expression levels of p53 and p21, which is responsible for cell cycle arrest under p53-mediated transcriptional regulation (4, 5), using immunoblotting. At 4 h after WGD induction, either in HCT116 or hTERT-RPE1, p53 remained unchanged in its amount from that in the non-treated asynchronous control but showed an evident band upshift representing its phosphorylation (Fig. S2B and C). At 15 h after WGD induction, p53 increased drastically in HCT116, whereas it did not drastically change from the untreated control, with an evident band upshift in hTERT-RPE1 (Fig. S2B and C). These data indicate that the responsiveness of p53 expression or phosphorylation is preserved in HCT116. Consistent with this idea, we also found that p53 expression in HCT116 was drastically upregulated, with an evident band upshift to a level equivalent to that in hTERT-RPE1 upon doxorubicin treatment (Fig. S2B).

However, we found a striking difference in the dynamics of p21 expression between HCT116 and hTERT-RPE1. In hTERT-RPE1 cells, p21 increased substantially as early as 4 h and reached an even higher level at 15 h after the treatments (Fig. S2B and C). On the other hand, in HCT116 cells, p21 expression was faint at 4 h and remained at a much lower level than in hTERT-RPE1 at 15 h after the treatments (Fig. S2B and C). Therefore, despite the upregulation of p53 protein, HCT116 failed to timely induce p21 to suppress cell cycle progression immediately after WGD induction or cytochalasin treatment.

Besides the above mechanism, prolonged mitosis also induces cell cycle arrest by forming the mitotic stopwatch complex in p53-proficient cells, with effects that are more pronounced when mitotic arrest is prolonged (6). Therefore, if cells spend different durations in mitosis upon MS- or CF-WGD induction, the mitotic stopwatch complex may have different impacts on the progression of the subsequent cell cycle. To address this possibility, we compared the progression of G1 entry after MS- or CF-WGD induction using live imaging of FUCCI-G1 (mKO2-hCdt1 (30-120)) in HCT116 (Fig. S2D) (7). There was no significant difference in the kinetics of FUCCI-G1 signal increase between MS- and CF-WGD conditions (Fig. S2E and F), suggesting that mitotic duration in MS- or CF-WGD was equivalent in our experimental conditions. We also analyzed the extent of mitotic stopwatch complex formation in HCT116 at 15 h after release from mitotic synchronization, with or without subsequent WGD induction (assessed by detecting p53 in immunoprecipitates obtained with anti-53BP1 antibodies; Fig. S2G and H). p53 content in anti-53BP1 immunoprecipitants in HCT116 cells after these treatments did not increase compared to that in the non-treated control (Fig. S2G and H), indicating that the mitotic stopwatch mechanism is inactive in HCT116, as previously reported (6). Therefore, MS- and CF-WGD cells experienced an equivalent degree of mitotic arrest upon WGD induction and failed to efficiently activate the mitotic stopwatch in the HCT116 background, which is unlikely to explain the proliferation difference between these conditions. Consistent with this idea, live-imaging analysis revealed that MS- and CF-WGD cells entered the first mitosis at equivalent times (Fig. S2I), indicating no difference in the progression of the first cell cycle.

While it remains unclear why p53-mediated cell cycle block is less stringent in HCT116, its mutation background may compromise p53 function. For example, HCT116 harbors a KRAS G13D mutation (8), which may affect p53 stability through the AKT-MDM2 axis (9). HCT116 also harbors a gain-of-function mutation in PPM1D/Wip1, which inactivates p53 signaling by dephosphorylating p53 (10). Given the relatively normal increase in p53 expression and phosphorylation upon WGD in HCT116 (Fig. S2C), we may also need to consider potential abnormalities in other aspects of the p53 pathway in the cell line. Since the MS-linked aggravation of nullisomic chromosome segregation was also observed in hTERT-RPE1 cells (Fig. S4F and G), the route-dependent outcomes of WGD cells are not specific to the HCT116 cellular background. However, given the impact of WGD on broader biological phenomena, it would be important in the future to further validate the generality of MS-driven nullisomy across more diverse organisms and cellular contexts.

#### Supporting text 2: Materials and methods

##### Cell culture and cell line establishment

HCT116 cells (RRID: CVCL_0291 from Riken BRC) or hTERT-RPE1 cells (RRID: CVCL_4388, a kind gift from Gohta Goshima) were cultured in McCoy's 5A (Wako) or DMEM/Ham's F-12 (Wako), respectively, supplemented with 10% fetal bovine serum (FBS) and 1× antibiotic-antimycotic solution (AA; Sigma-Aldrich) at 37°C with 5% $CO_2$. Transgenic cell lines were established by transfecting HCT116 cells with plasmids encoding the corresponding transgenes using JETPEI (Polyplus-transfection), followed by selection of positive clones with the appropriate antibiotics. The RAD21-mAC HCT116 cell line expressing OsTIR1(F74G) (11) was a kind gift from Masato Kanemaki (National Institute of Genetics, Japan). Cell lines used in this study are listed in Dataset S1. All experiments were performed with mycoplasma-free cells (tested using the MycoStrip mycoplasma detection kit; InvivoGen).

##### Plasmids, compounds, and antibodies

The plasmid vectors constructed or used in this study are listed with the primer information in Dataset S2. pSpCas9(BB)-2A-GFP (PX458)(12), CDNA3.1-dCas9-2xNLS-EGFP (13), and mEOS2-H2B-6 (14) were purchased from Addgene (plasmid #48138, #74710, and #57384, respectively). Compounds and antibodies used in this study are listed in Dataset S3. Anti-CENP-C antibody was a kind gift from Kinya Yoda (15). The siRNAs used in this study are 5'-GCCUAGGUGUCCUUGAGCUtt-3' (Sororin; DNA in lowercase), 5'-CGUACGCGGAAUACUUCGAtt-3' (luciferase), and siGENOME TP53 siRNA (D-003329-26, Dharmacon). siRNA transfection was performed using Lipofectamine RNAiMAX (Thermo Fisher Scientific).

##### WGD induction

To induce WGD, cells were first arrested at prometaphase by treating with 40 ng/mL nocodazole for 4 h, washing 3 times with supplemented culture medium, and then shaking off. For CF-WGD induction, these shaken-off mitotic cells were treated with 5 μg/mL cytochalasin B for 2 h. For MS-WGD induction, the shaken-off mitotic cells were co-treated with 40 ng/mL nocodazole and 10 μM RO3306 for 2 h. As an alternative way to induce MS-WGD, cells were arrested at prometaphase by treating with 50 μM monastrol for 4 h, washed 3 times with supplemented culture medium, shaken off, and co-treated with 50 μM monastrol and 5 μM reversine for 2 h. Throughout the manuscript, the time of WGD induction was defined as the time point at which these WGD-induction inhibitors were added to cell cultures. After 2-h WGD induction, cells were washed 3 times with supplemented culture medium. The efficiency of WGD induction was evaluated by the presence of excess centrosomes or by increases in nuclear area.

For MS-WGD induction in mock- or Sororin-depleted cells, siRNA was transfected at 50 h prior to the induction of MS-WGD. In this case, cells were once arrested at prometaphase by 4-h nocodazole treatment, washed 3 times with supplemented culture medium, and then additionally arrested at metaphase by 2-h MG132 treatment before inducing MS-WGD. For MS-WGD induction in RAD21-mAC cells, 1μg/mL doxycycline and 1 μM 5-Ph-IAA were sequentially administered to the cell culture at 26 h and 2 h prior to MS-WGD induction, respectively. Doxycycline and 5-Ph-IAA were removed by washing mitotically arrested cells with supplemented culture medium before shake-off.

To analyze the first mitosis after WGD in hTERT-RPE1 cells, cells were transfected with p53-targeting siRNA at 26 h prior to WGD induction to override p53-mediated cell cycle arrest.

Flow cytometry

Flow cytometric DNA content analyses were performed using a JSAN desktop cell sorter (Bay Bioscience). To analyze DNA content immediately after WGD induction, cells were trypsinized using 0.05% trypsin-EDTA (Wako) and stained with 10 µg/ml Hoechst 33342 (Dojindo) for 1 h at 37°C. To analyze DNA content in cell populations at 8 d after WGD induction, cells were trypsinized, fixed using 70% ethanol overnight at 4°C, and stained with 1 µg/ml propidium iodide (Dojinbo) for 2 h at 37°C. The stained cells were filtered through a cell strainer and applied to the cell sorter ($1.5 \times 10^4$ cells were counted for each analysis). Data analyses were conducted using FlowJo software (BD Sciences).

Colony formation assay and DNA content quantification

For the colony formation assay, the number of mitotically synchronized cells was counted using a hemocytometer right after the mitotic shake-off for WGD induction. Then, 2,000 cells were seeded into each well of 6-well dishes to induce either CF-WGD or MS-WGD. After WGD induction, cells were washed 3 times with supplemented medium and further incubated for 8-9 d. Cells forming colonies were live-stained with 10 µg/mL Hoechst 33342 for 30 min at 37°C, subjected to medium change with supplemented phenol red-free McCoy's 5A, and imaged under a Ti2 microscope.

The tile images covering the entire wells were acquired using the Grid/Collection Stitching plugin in µManager software (16), followed by stitching these tiles using the Image Stitching plugin in FIJI software (NIH). The images of individual colonies were then cropped from the stitched well image and subjected to automated single-nucleus segmentation using the StarDist plugin to quantify nuclear Hoechst fluorescence intensities of individual cells. The ImageJ macro files used for the automated segmentation are provided as Supplemental material S3. The nuclear Hoechst intensities obtained from an individual colony were plotted into a histogram and subjected to Kernel density estimation to estimate the Hoechst intensity value corresponding to the G1 or G2/M peak. The average G1 Hoechst intensity in diploid control colonies was set to 2C, and the G1 Hoechst intensities of all colonies in WGD samples were normalized to this value (Fig. 1F). Then, colonies with their G1 Hoechst intensities of more than 3C were categorized as "polyploid." A single colony consisting of a mixture of diploid and polyploid cell populations was also categorized as polyploid, and its polyploid G1 value was used for the analysis.

Live imaging

For live imaging of the first mitosis after WGD induction, we seeded WGD-induced cells on an 8-well cover glass-bottom chamber (zell-kontakt GmbH), replaced culture media with supplemented phenol red-free McCoy's 5A (Cytiva) at 2 h or 15 h after WGD induction, and started cell imaging from 15 h or 16 h after WGD induction, respectively. For visualizing chromosomes in RAD21-mAC live imaging, 100 nM SiR-DNA (Cytoskeleton, Inc.) was treated 1 h before the start of live imaging. For live imaging of C9-1 foci during MS-WGD induction, the cells arrested at prometaphase were directly subjected to WGD induction without shake-off on an 8-well cover glass-bottom chamber. For analyzing cell cycle entry immediately after WGD induction, prometaphase-arrested cells expressing FUCCI biosensors (7) were treated either with 40 ng/mL nocodazole and 10 µM RO3306 (to induce MS-WGD) or 5 µg/mL cytochalasin B (to induce CF-WGD), and imaging was initiated from 15 min after the compound administration without removing these compounds.

Immunostaining

Cells were fixed with 100% methanol at -20°C for 10 min, treated with BSA blocking buffer (150 mM NaCl, 10 mM Tris-HCl, pH 7.5, 5% BSA, and 0.1% Tween 20) for 30 min at 25°C, incubated with the first antibodies overnight at 4°C, and with fluorescence-conjugated secondaries overnight at 4°C. DNA was stained with 1 µg/mL DAPI. Following each treatment, cells were washed 3 times with phosphate-buffered saline (PBS).

Fluorescence in situ hybridization (FISH)

We conducted FISH using ZytoLight FISH-Cytology Implementation kit (ZytoVision GmbH) following the manufacturer's instructions. Cells were treated with the pepsin solution (Cytology Pepsin Solution, ES2, ZytoVision GmbH) or 0.05% pepsin in 10 mM HCl for 10 min at 37°C and fixed with 1% PFA in 1x $MgCl_2$ solution (PT4, ZytoVision GmbH; 1:9 mixture of 10x $MgCl_2$ solution and PBS). The fixed cells were dehydrated with an ethanol series (70, 90, and 100%; for 1 min, each) and air-dried. Samples were then treated with 1x ZytoLight FISH probe solution (ZytoLight SPEC 1p12 Probe, ZytoLight CEN 9 Probe, or ZytoLight CEN 17 Probe), denatured for 5 min at 72°C, incubated overnight at 37°C, then washed once with 1x Cytology Stringency Wash Buffer SSC (WB7, ZytoVision GmbH) for 2 min at 70°C, and washed once with 1x Cytology Wash Buffer SSC (WB8, ZytoVision GmbH) for 1 min at 25°C. DNA was stained with 1x DAPI/DuraTect-Solution (MT7, ZytoVision GmbH).

Chromosome spread

Cells were treated with 40 ng/mL nocodazole for 4 h at 37°C, trypsinized for 5 min at 37°C, washed twice with PBS, and suspended in 0.056 M KCl. After incubating for 20 min at 25°C, the swollen cells were spun down and fixed with 5 mL of a methanol-glacial acetic acid (3:1) solution for 20 min on ice. The fixed cell suspension was applied dropwise onto slide glasses from approximately 10 cm above the slides, air dried, and mounted using Fluoromount (Diagnostic BioSystems). Chromosome spreads were stained with 167 nM SYTOX Green (Thermo Fisher Scientific).

Quantification of DNA contents in live imaging

To distinguish WGD-induced tetraploids and contaminated (non-WGD-induced) diploids, nuclear areas of individual cells right before the entry to the first mitosis were quantified using automated single-nucleus segmentation by FIJI. Based on the difference in nuclear area between stable diploid and tetraploid control cells (Fig. S3A), we set the threshold of nuclear area between contaminated diploids and WGD-induced tetraploids at 160 $\mu m^2$.

To evaluate the validity of F1 cell DNA content estimation based on H2B-mCherry signals, we live-stained MS- or CF-WGD-induced H2B-mCherry cells with 100 ng/mL Hoechst 33342 for 15 min at 37°C and subjected them to live imaging under a Ti2 microscope. Nuclear signals of H2B-mCherry or Hoechst in F1 cells were obtained using automated segmentation by FIJI at the time point when these cells became attached to the dish bottom after the first mitosis. After subtracting the non-cell area background intensities, the DNA content of an individual F1 cell, $D_i$, was calculated using the equation below, with an assumption that the DNA contents of all sister F1 cells from a single mother cell sum up to 8C:

$$D_i = \frac{8C \times I_i}{\sum I_i}$$

where $I_i$ is the corrected intensity of H2B-mCherry or Hoechst signal in an individual F1 cell. Since the ploidy values estimated by H2B-mCherry signals recapitulated those by Hoechst (Fig. S3E), we used the H2B-mCherry signal as an indicator of F1 cell ploidy level in the live imaging analyses. In live imaging of RAD21-mAC cells, the SiR-DNA signal was used to estimate ploidy using the above equation.

Quantitative analyses of the centrosome and C9-1 centromere positions in the first mitosis

To analyze the centrosome and C9-1 centromere positions, EGFP-GCP3/dCas9-EGFP (C9-1)/H2B-mCherry cells were imaged either by fixed or live cell microscopy. For fixed-cell imaging, we fixed the WGD-induced cells at 16 h after WGD induction and distinguished the phases of the first mitosis based on the chromosome configuration labeled with H2B-mCherry. The positions of the centrosomes and C9-1 foci were manually measured using the cell counter tool in FIJI. The centroid of the mitotic chromosome masses was measured using FIJI, from the auto-segmented regions of H2B-mCherry signals on the middle slice of Z-sections covering the entire chromosome masses. In live-imaging data, the three-dimensional positions of the centrosomes and C9-1 were manually measured using the cell counter tool in FIJI.

Microscope

Fixed cells were observed under either of the following microscopes: A TE2000 microscope (Nikon) equipped with a ×100 1.45 NA Plan-Apochromatic, a CSU-X1 confocal unit (Yokogawa), and an iXon3 electron multiplier-charge coupled device (EMCCD) camera (Andor): A Ti2 microscope (Nikon) with a ×100 1.45 NA Plan-Apochromatic, and Zyla4.2 sCMOS camera (Andor) or ZL41 Cell sCMOS camera (Andor): A Ti microscope (Nikon) with ×100 1.4 NA Plan-Apochromatic, a CSU10 confocal unit (Yokogawa), and an ImagEM (EMCCD) camera (Hamamatsu).

Live cell imaging was conducted under either of the following microscopes at 37°C with 5% $CO_2$: A TE2000 microscope with x60 1.4 NA Plan-Apochromatic, CSU-X1, and iXon3: A Ti2 microscope with ×20 0.75 NA Plan-Apochromatic, and Zyla4.2: A Ti microscope with a x60 1.4 NA Plan-Apochromatic, CSU10, and ImagEM. Post-WGD cell colonies were observed under a Ti2 microscope with x4 0.13 NA Plan-Fluor and Zyla4.2. Image acquisition was controlled by µManager.

Theoretical modeling of F1 cell viability

F1 cell viability was estimated based on the empirically obtained frequency of nullisomic chromosome segregation of C9-1 at the first mitosis, with the assumptions that 1) nullisomic segregation occurred independently with the same frequency as chromosome 9 for all other chromosome species, and 2) F1 cells were viable when they possessed at least one copy of all chromosome species. Then, the probability of viable F1 cell formation, $p_{viable}$, was modeled as

$$p_{viable} = \left(1 - p_{nullisomy}\right)^{23},$$

where $p_{nullisomy}$ is the probability of nullisomy of a chromosome species, estimated based on the frequency of nullisomic segregation for chromosome 9. The nullisomy frequency in mononucleated F1 cells was estimated based on the assumption that each chromosome mass analyzed in Fig. 3B and S4C was segregated into a separate daughter cell. To estimate the nullisomy frequency in binucleated F1 cells, we first quantified the number of C9-1 foci in all possible combinations of two chromosome masses in Fig. 3B and S4C. Then, the binucleated nullisomy frequency was estimated as the proportion of the combinations possessing no C9-1 foci in all combinations. In Fig. 3H, the viability of only binucleated cells generated via tetrapolar chromosome segregation was compared, since those via tripolar chromosome segregation were rare and not suitable for quantification. The estimated F1 viability was then expressed as a percentage for comparison with the empirical values obtained by live imaging analysis. Numerical values assigned to the model are listed in Dataset S4.

Theoretical modeling of nullisomic segregation frequency

Based on the quantitative analysis of live imaging of C9-1 dynamics during the first mitosis (Fig. 5F), we modeled the relationship between the centrosome-centromere distance at NEBD and the frequency of nullisomic homologue segregation at the subsequent anaphase, $f_{nullisomy\ model}$ as

$$f_{nullisomy\ model} = \frac{\sum(N_i \cdot f_i)}{\sum N_i}$$

where $N_i$ is the number of pairs of centrosomes and their closest centromere that falls in each distance bin defined in Fig. 5F, and $f_i$ is the frequency of nullisomic segregation in the corresponding distance bin estimated by the empirical data in Fig. 5F. Model-predicted frequency of nullisomic segregation was then expressed as a percentage for comparison with the empirical values obtained by imaging analysis. Numerical values assigned to the model are listed in Dataset S4.

Immunoblotting and immunoprecipitation

For immunoblotting, cells were lysed in SDS/PAGE sample buffer (1.125% SDS, 35 mM Tris-HCl, pH 6.8, 11.25% glycerol, 5% 2-mercaptoethanol), boiled for 5 min, and subjected to SDS/PAGE. Separated proteins were transferred onto an Immun-Blot PVDF membrane (Bio-Rad). The blotted membranes were blocked with 0.3% skim milk in Tween Tris-buffered saline (TTBS; 50 mM Tris, 138 mM NaCl, 2.7 mM KCl, and 0.1% Tween 20), incubated with the first antibodies for 1 or 2 h at 25°C, or overnight at 4°C, and incubated with horseradish peroxidase (HRP)-conjugated secondary antibodies for 1 or 2 h at 25°C. Each step was followed by 3 washes with TTBS. Signals were detected using the ezWestLumi plus ECL Substrate (ATTO, Tokyo, Japan) with a LuminoGraph II chemiluminescent imaging system (ATTO).

For immunoprecipitation, control or WGD-induced cells were extracted with RIPA buffer (50 mM Tris-HCl (pH 8.0), 150 mM NaCl, 1% NP-40, 0.5% sodium deoxycholate, 0.1% SDS, 1 μM dithiothreitol, 1×protease inhibitor cocktail (cOmplete, Roche) and 1×PhosSTOP (Roche)) for 15 min on ice, precleaned with Protein G Sepharose 4 Fast Flow beads (17061805, Cytiva) for 1 h, incubated with 200 ng anti-53BP1 antibody for 1 h, and then with Protein G Sepharose beads for 4 h. Immunoprecipitants were washed with RIPA buffer three times, lysed in SDS/PAGE sample buffer and subjected to immunoblotting. All procedures were conducted at 4°C after extraction.

Statistical analysis

Analyses for significant differences between the two groups were conducted using the two-tailed Welch t-test, the Fisher exact test, or the Brunner-Munzel test in R software (The R Foundation). Multiple group analyses were conducted using the Fisher exact test with the Benjamini-Hochberg multiple testing correction, the Steel-Dwass test, or the Dwass-Steel-Critchlow-Fligner (DSCF) test in R software. Statistical significance was set at $p < 0.05$. P-values are indicated in figures or the corresponding figure legends.

Supplementary figure 1

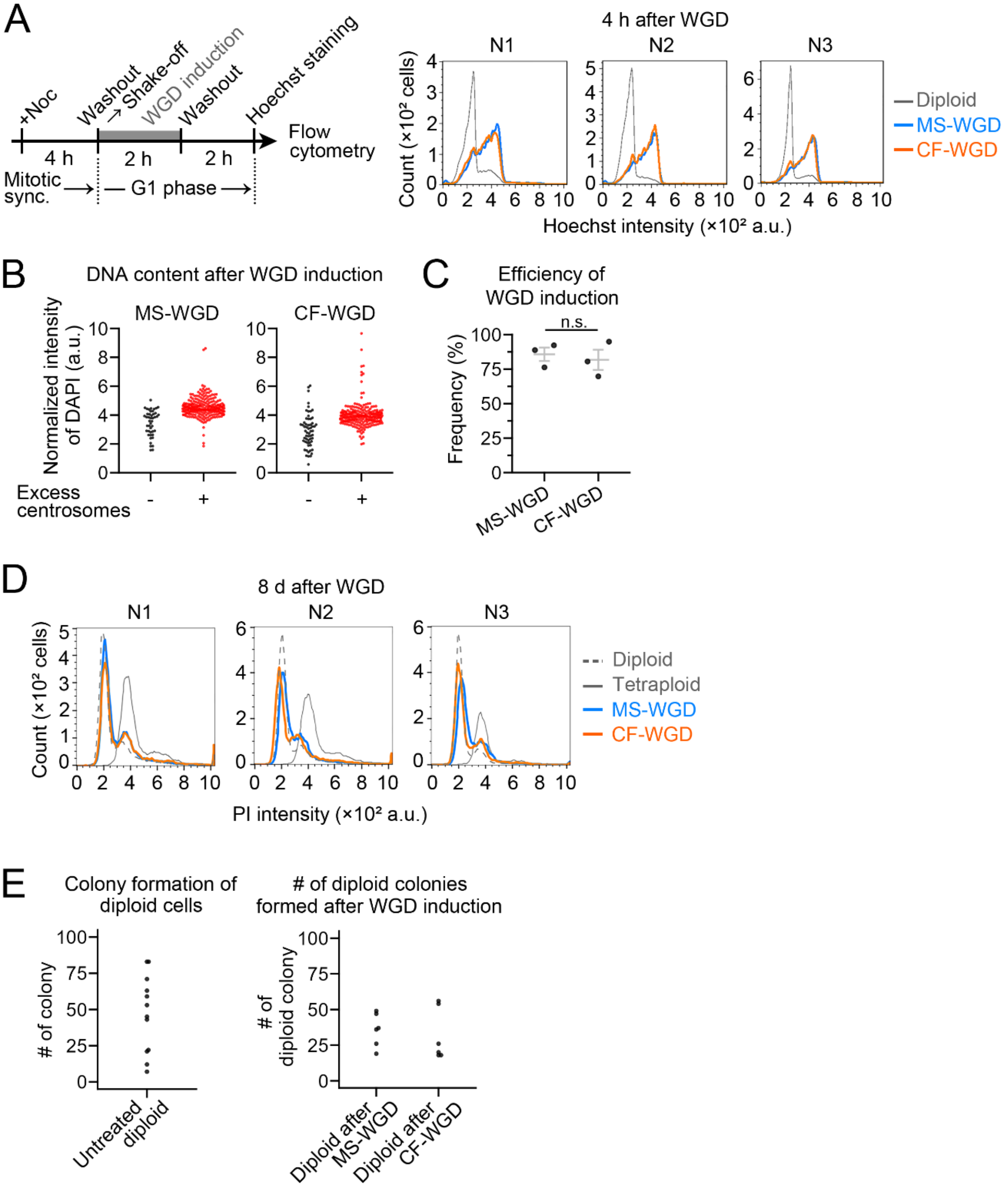

**Figure S1. Evaluation of the efficiency of WGD induction**
**(A)** Flow cytometric analysis of DNA content in DMSO-treated diploid control, MS-, or CF-WGD-induced cell culture. Cells were sampled at 4 h after the initiation of WGD induction (schematized on the left). DNA was stained with Hoechst 33342. Results from three independent experiments are shown. **(B)** Quantification of DNA content and the possession of excess (>2) centrosomes in WGD-induced cell cultures. WGD cells were defined based on the possession of excess centrosomes (red dots; the centrosomes were marked by Cep192 as depicted in Fig. 1B). Note that the substantial overlap in DNA contents between diploids (black dots) and WGD cells is presumably due to the asynchronous cell cycle stage distributions in the former. **(C)** Efficiency of WGD induction in B. Means ± s.e. of three independent experiments. At least 292 cells were analyzed in each condition. There is no statistically significant difference between conditions ($p$ = 0.87, the Brunner-Munzel test). **(D)** Flow cytometric analysis of DNA content in DMSO-treated diploid control, MS-, or CF-WGD-induced cell culture. Cells were cultured in bulk populations for 8 d after WGD induction and then fixed. DNA was stained with propidium iodide. DNA content in a stable tetraploid HCT116 line was also measured as a reference. Results from three independent experiments are shown. **(E)** A comparison of the number of colonies formed from untreated diploids (left) and that of diploid cell colonies formed after MS- or CF-WGD induction in Fig. 1D (right). The number of untreated diploids initially seeded was determined based on the estimated number of contaminated diploids in C. Data were obtained from three independent experiments, each with two replicates.

### Supplementary figure 2

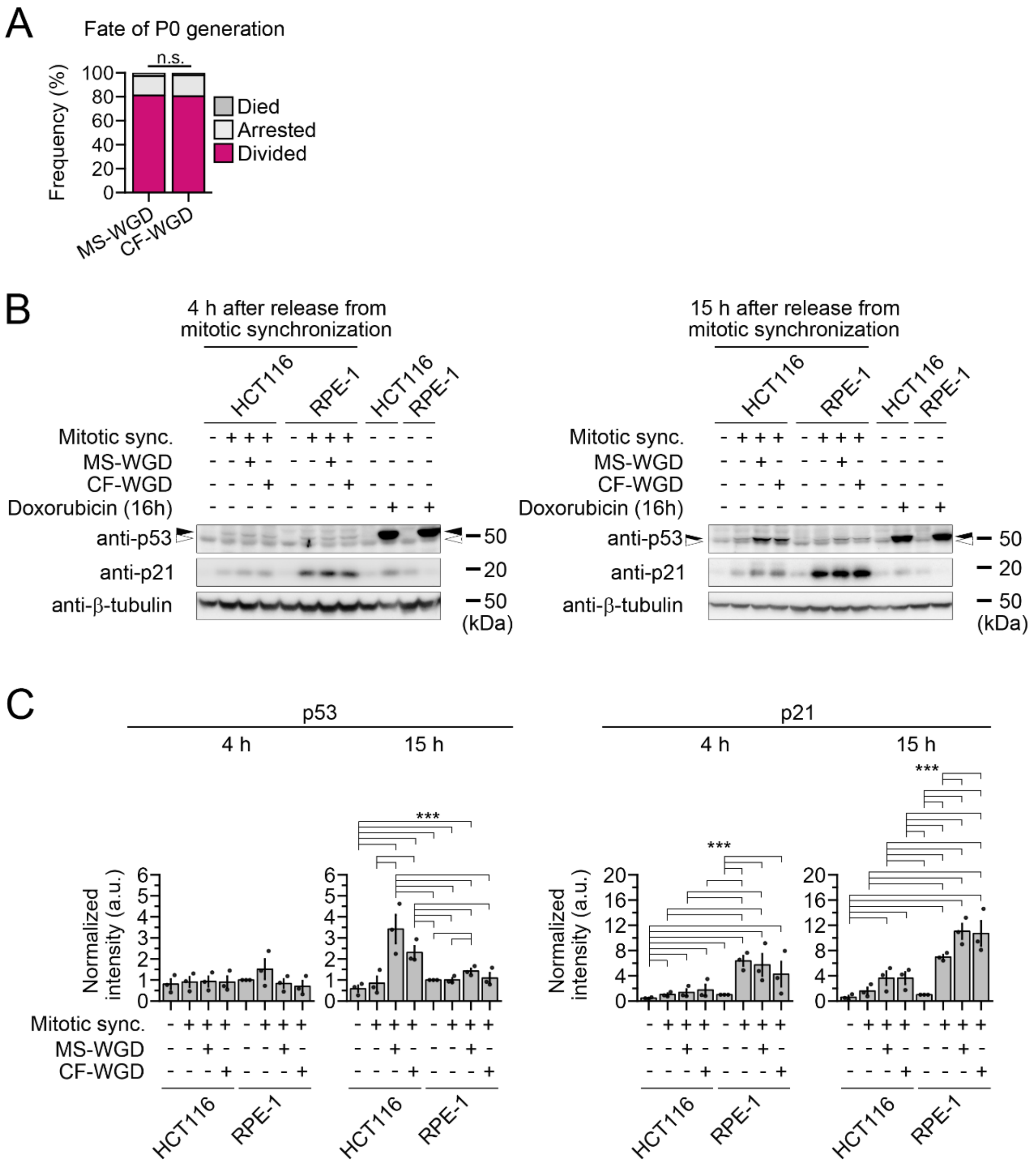

### Supplementary figure 2 (continued)

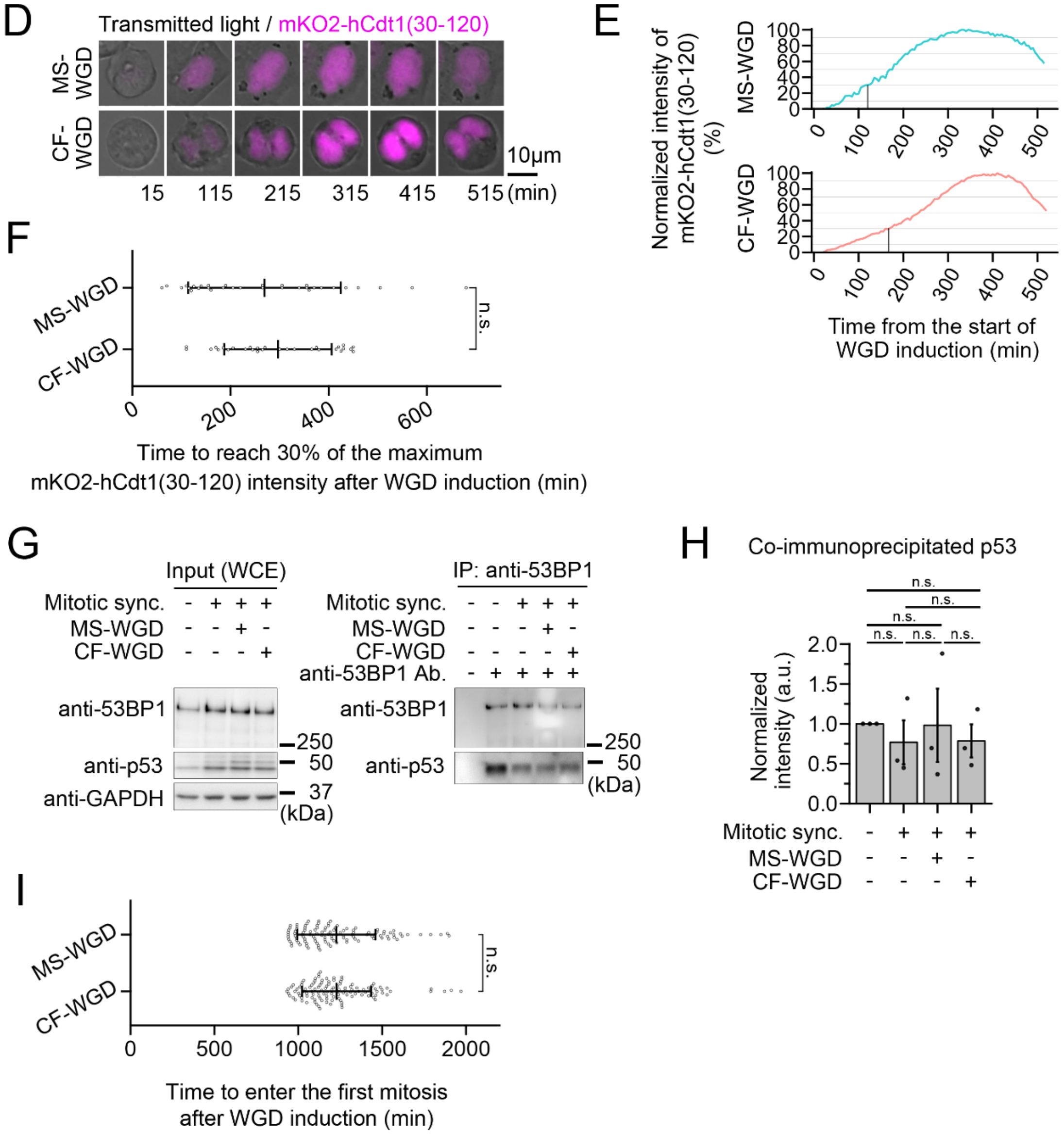

**Figure S2. p53 signaling during the first cell cycle after WGD in HCT116 cells**
**(A)** Fates of P0 cells (the first cell cycle) after MS- or CF-WGD induction. At least 161 cells from three independent experiments were analyzed. There is no statistically significant difference between conditions ($p = 0.83$, the Fisher exact test). **(B)** Immunoblotting of p53 and p21 in HCT116 or hTERT-RPE1 cells at 4 or 15 h after the release from mitotic synchronization with or without subsequent WGD induction. Asynchronous culture was analyzed as a non-treated control. Open or closed arrowheads indicate unshifted or upshifted p53 bands, respectively. β-tubulin was detected as a loading control. **(C)** Quantification of the relative expression of p53 or p21 in B. Protein loading differences were corrected based on β-tubulin signals. Mean ± s.e. of three independent experiments. Asterisks indicate statistically significant differences among conditions (***$p < 0.001$, the Steel-Dwass test). **(D)** Time-lapse images of FUCCI-G1 (mKO2-hCdt1(30-120)) upon MS- or CF-WGD induction. Images were taken at 5-min intervals. The time after the start of WGD induction (the release from the prior mitotic synchronization) was indicated. **(E)** Examples of the time course of FUCCI-G1 intensity in D, expressed as percentages relative to the maximum intensities. Black lines indicate the time points at which FUCCI-G1 intensities reach 30% maximum. **(F)** Time at which FUCCI-G1 intensity reached 30% maximum after WGD induction. Means ± s.d. of at least 27 cells from two independent experiments. There is no statistically significant difference between conditions ($p = 0.43$, the Welch's t-test). **(G)** Immunoblotting of p53 or 53BP1 in anti-53BP1 immunoprecipitant from HCT116 at 15 h after the release from mitotic synchronization with or without subsequent WGD induction. The immunoprecipitant from asynchronous culture was analyzed as a non-treated control. Immunoblots of whole-cell extracts are shown as inputs (GAPDH was detected as a loading control). **(H)** Quantification of the relative p53 content in the immunoprecipitants in G. Mean ± s.e. of three independent experiments. There are no statistically significant differences among conditions (n.s.: not significant, the Steel-Dwass test). **(I)** Time at which cells entered the first mitosis after WGD induction. Means ± s.d. of at least 108 cells from three independent experiments. There is no statistically significant difference between conditions ($p = 0.97$, the Welch's t-test).

### Supplementary figure 3

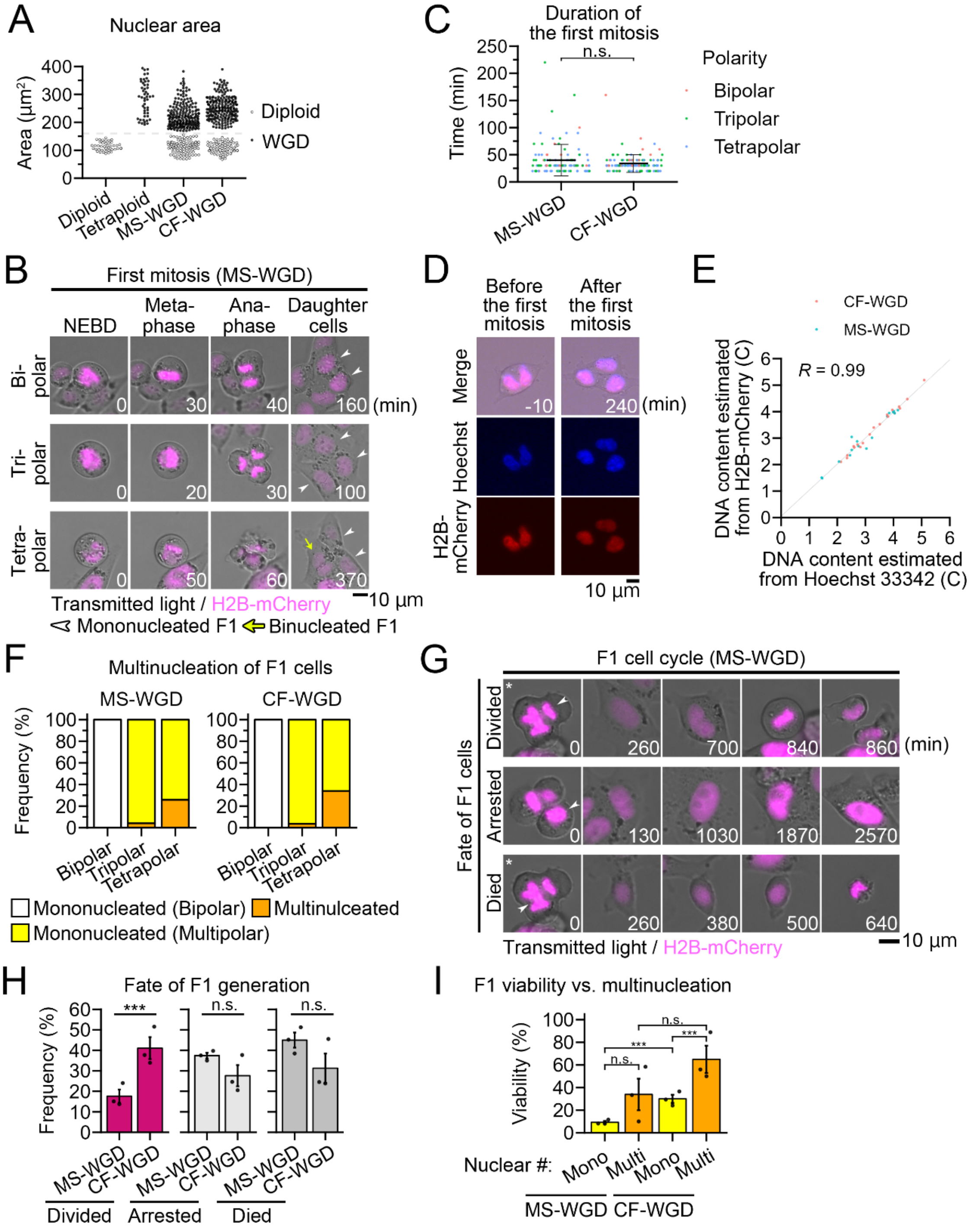

**Figure S3. Live imaging of the first mitosis and F1 cell cycle after MS- or CF-WGD**
**(A)** Quantification of nuclear areas in diploid, stable tetraploids, or MS- or CF-WGD-induced cells immediately before the entry into mitosis (the first mitosis in case of the WGD-induced cells). Diploid and tetraploid cells were clearly distinguished based on the nuclear area. Therefore, cells with nuclear areas of >160 $\mu m^2$ (the broken line) were defined as WGD-induced cells in the MS- or CF-WGD condition. At least 41 cells from at least two independent experiments were analyzed. **(B)** Time-lapse images of H2B-mCherry cells during the first mitosis after MS-WGD. Images were taken at 10-min intervals. The NEBD timing is set to 0 min. White arrowheads indicate "mononucleated" daughter cells formed through the first mitosis. The yellow arrow indicates a binucleated daughter cell. **(C)** Mitotic duration from NEBD to anaphase onset at the first mitosis after MS- or CF-WGD in Fig. 2B and S3B. Means ± s.d. of at least 108 cells from three independent experiments. There is no statistically significant difference between conditions ($p$ = 0.054, the Welch's t-test). **(D)** Fluorescence microscopy of a CF-WGD-induced H2B-mCherry cell stained with Hoechst 33342 (at 16 h after WGD-induction). **(E)** Relationship between the DNA content of F1 daughter cells estimated based on the signal segregation ratio of Hoechst or H2B-mCherry. DNA content estimation was conducted as depicted in Fig. 2D. The correlation coefficient between the variables is shown. **(F)** Frequency of daughter cell multinucleation at the first mitosis after MS- or CF-WGD in Fig. 2B and S3B. At least 28 cells from three independent experiments were analyzed in each polarity. **(G)** Time-lapse images of H2B-mCherry cells during the F1 cell cycle after MS-WGD. Images were taken at 10-min intervals. Dividing daughter cells indicated by arrowheads are traced in the subsequent time frames. The timing of anaphase onset is set to 0 min. The panels with asterisks show an identical dividing mother cell, with its different daughters traced in subsequent time frames in different rows. **(H, I)** Frequency of F1 cell fate (H) or viability of F1 generation that was sorted based on the presence or absence of daughter cell multinucleation after multipolarization (I) in the live imaging in Fig. 2B and S3B. Means ± s.e. of three independent experiments. At least 55 cells (H) or 28 cells (I) were analyzed in each category. Asterisks indicate statistically significant difference between conditions (n.s.: not significant, ***$p$ < 0.001, the Brunner-Munzel test in H, or the Steel-Dwass test in I).

#### Supplementary figure 4

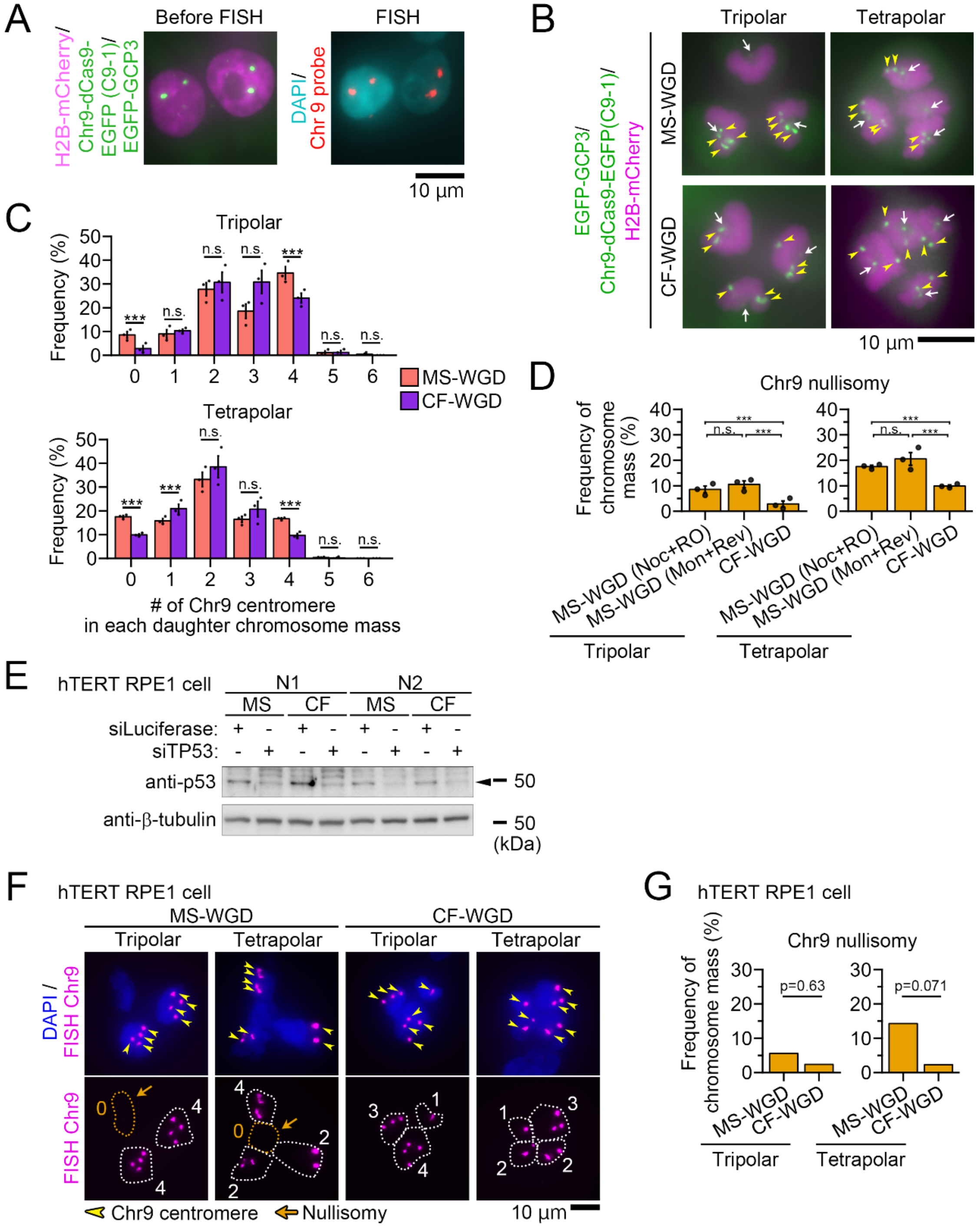

#### Supplementary figure 4 (continued)

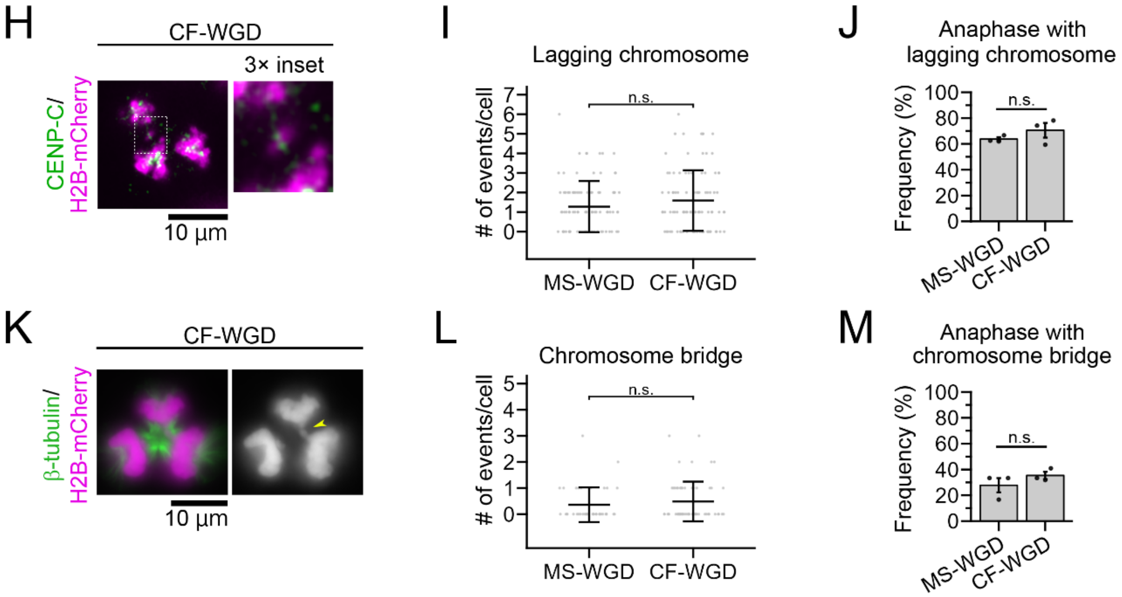

**Figure S4. Imaging analyses for multipolar chromosome segregation patterns after WGD**
**(A)** Comparison of chromosome 9 centromere labeling with dCas9-EGFP (left; imaged before cell fixation) and that with a specific FISH probe (right; imaged after cell fixation) in diploid interphase cells. Fluorescence microscopy of identical cells before and after cell fixation. DNA was stained with DAPI after fixation. Note that dCas9-EGFP, EGFP-GCP3, and H2B-mCherry signals observed in the pre-fixed sample were breached in the process of cell fixation for FISH staining. **(B)** Fluorescence microscopy of anaphase cells at the first mitosis after MS- or CF-WGD. The centrosomes (white arrows), centromeres of chromosome 9 (yellow arrowheads), and chromosomes were labeled by EGFP-GCP3, dCas9-EGFP with specific sgRNA (C9-1), and H2B-mCherry, respectively. For the demonstration, identical cells are shown in Fig. 3A. **(C)** Histograms of C9-1 foci numbers detected in segregated chromosome masses in anaphase in Fig. 3A. Means ± s.e. of three independent experiments. Asterisks indicate statistically significant difference between MS- and CF-WGD in each category (n.s.: not significant, ***$p < 0.001$, the Brunner-Munzel test). **(D)** The proportion of segregated chromosome masses devoid of any C9-1 focus. Means ± s.e. of three independent experiments. At least 252 segregated chromosome masses (presumptive daughter nuclei) from 84 cells were analyzed in each condition in C and D. Asterisks indicate statistically significant difference between conditions (n.s.: not significant, ***$p < 0.001$, the Steel-Dwass test). Data identical to those in Fig. 3B are shown for MS-WGD (Noc+RO) and CF-WGD, for comparison with the MS-WGD (Mon+Rev) condition. **(E)** Immunoblotting of p53 in mock- or p53-depleted hTERT-RPE1 cells sampled at 18 h after WGD induction. β-tubulin was detected as a loading control. **(F)** Fluorescence microscopy of anaphase cells at the first mitosis after MS- or CF-WGD in p53-depleted hTERT-RPE1 cells. Centromeres of chromosome 9 were stained with specific FISH probes. Chromosomes were stained with DAPI. Yellow arrowheads indicate the centromeres. Orange arrows indicate chromosome masses devoid of the labeled homologous centromeres (nullisomy). Broken lines indicate the contours of chromosome masses. The number of centromere foci in each chromosome mass is indicated in the bottom panels. **(G)** Frequency of the segregated chromosome masses devoid of any centromere signals for chromosome 9 in F. Pooled data from three independent experiments. At least 28 segregated chromosome masses (presumptive daughter nuclei) were analyzed in each polarity. Asterisks indicate statistically significant difference between conditions ($p$-values of the Fisher exact test are shown). **(H, K)** Fluorescence microscopy of H2B-mCherry and the kinetochores (marked by CENP-C immunostaining in H) or spindle microtubules (β-tubulin immunostaining in K) in anaphase at the first mitosis after CF-WGD. The right panel in H shows a 3x enlarged image of a lagging chromosome. The yellow arrowhead indicates a chromosome bridge in K. **(I, L)** Numbers of lagging chromosomes (I) or chromosome bridges (L) found in each cell in H or K, respectively. There is no statistically significant difference between conditions in I or L ($p = 0.14$ or $p = 0.39$, respectively, the Welch's t-test). At least 39 cells from three independent experiments were analyzed in each condition. **(J, M)** Frequency of anaphase cells with at least one lagging chromosome (J) or chromosome bridge (M) in H or K, respectively. Means ± s.e. of three independent experiments. At least 39 cells were analyzed in each condition. There is no statistically significant difference between conditions in J or M ($p = 0.67$ or $p = 0.55$, respectively, the Brunner-Munzel test).

### Supplementary figure 5

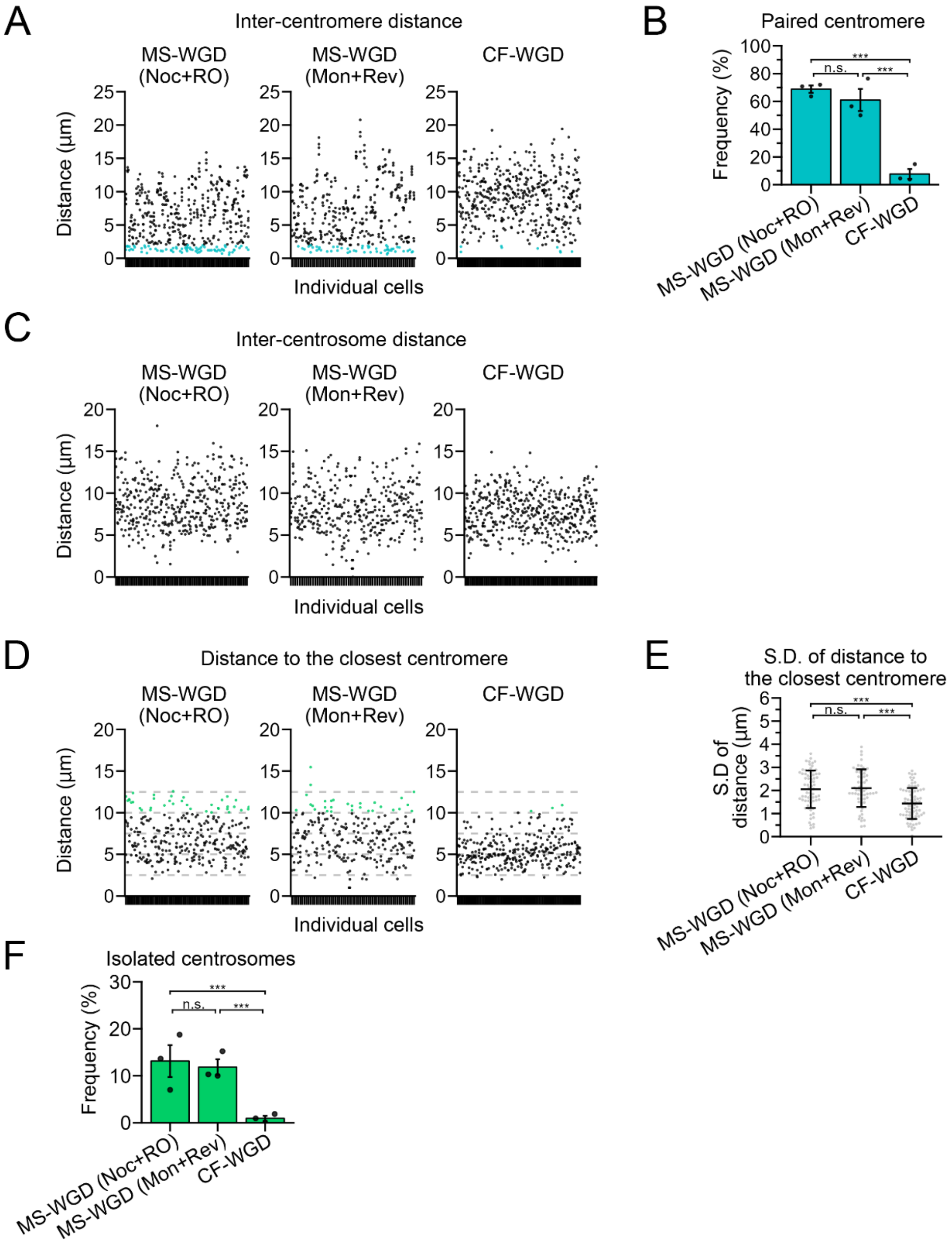

**Figure S5. Homologue distribution is commonly skewed after MS-WGD induced by different methods**
**(A, C, D)** Quantification of inter-centromere distance (A), inter-centrosome distance (C), or the distance from each centrosome to its closest centromere (D) at the entry into the first mitosis after MS- or CF-WGD. Each column represents the distances obtained in an individual cell. Paired centromeres or isolated centrosomes are colored in blue (A) or green (D), respectively. At least 60 cells from three independent experiments were analyzed. **(B, F)** Frequency of cells with paired C9-1 centromeres (B) or frequency of isolated centrosomes (F) in A or D, respectively. Means ± s.e. of three independent experiments. Asterisks indicate statistically significant difference between conditions (n.s.: not significant, ***$p < 0.001$, the Steel-Dwass test). **(E)** Standard deviation of the distances in each cell in D. Asterisks indicate statistically significant difference between conditions (n.s.: not significant, ***$p < 0.001$, the DSCF test). Identical data to those in Fig. 4C, F, or G are shown for MS-WGD (Noc+RO) and CF-WGD in B, E, or F, respectively, for comparison with the MS-WGD (Mon+Rev) condition.

### Supplementary figure 6

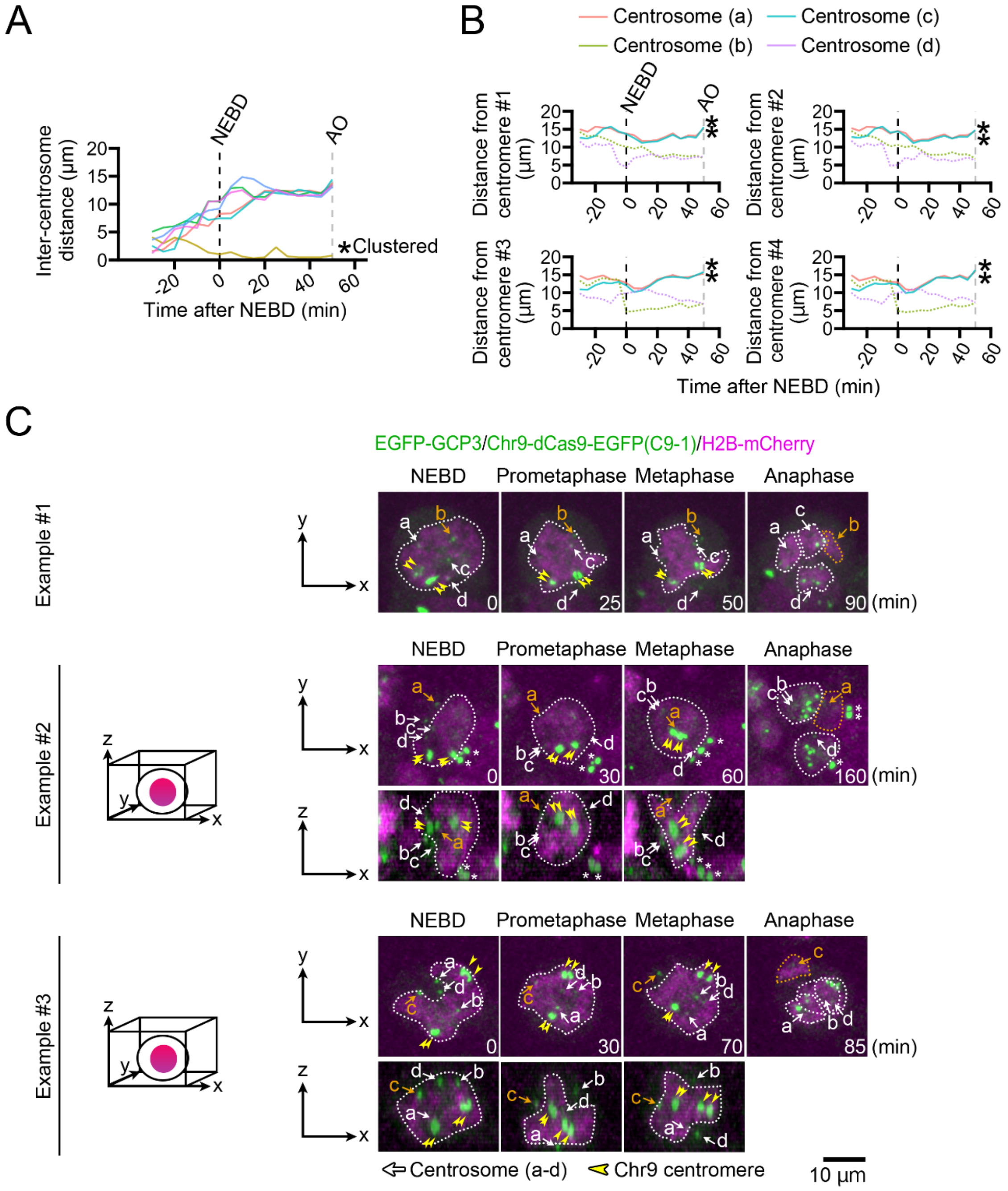

#### Supplementary figure 6 (continued)

D

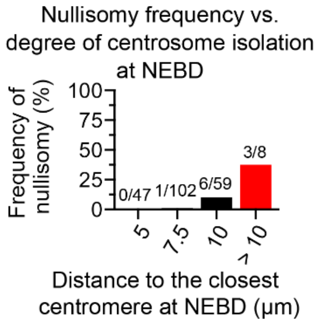

**Figure S6. Quantitative analysis of centromere and centrosome dynamics during the first mitosis after MS-WGD**

**(A)** Examples of the time course of inter-centrosome distance in Fig. 5A. Broken lines indicate the timing of NEBD or anaphase onset (black or gray, respectively). The asterisk indicates a clustered centrosomal pair. **(B)** Examples of the time course of distance between each centrosome-centromere combination in Fig. 5A. Each graph represents the distances from a single C9-1 focus to all 4 centrosomes (a-d). Asterisks indicate the centrosomes that did not capture the corresponding centromere. **(C)** Time-lapse images of H2B-mCherry, EGFP-GCP3, and dCas9-EGFP with C9-1 sgRNA throughout the first mitosis after MS-WGD. Yellow arrowheads indicate C9-1 foci. White arrows or orange arrows indicate centrosomes that captured or did not capture C9-1 foci, respectively, during anaphase. To improve the visibility of complex shapes of mitotic chromosome masses (highlighted by broken lines), top and side views (XY and XZ planes, respectively) of three-dimensional reconstructions are shown for examples #2 and 3. Note that the initial positions of C9-1 foci tended to be reflected in their positions on the multipolar metaphase plates and those in anaphase chromosome masses. Asterisks indicate C9-1 foci in neighbor cells. **(D)** The frequency of the centrosomes that failed to capture any C9-1, which was calculated for each bin of the closest centrosome-centromere distance at NEBD. Two hundred sixteen centrosomes from 54 CF-WGD cells from nine independent live-imaging experiments were analyzed. The number of centrosomes failing to capture any C9-1 per total centrosomes in each bin is shown on top of each graph bar.

### Supplementary figure 7

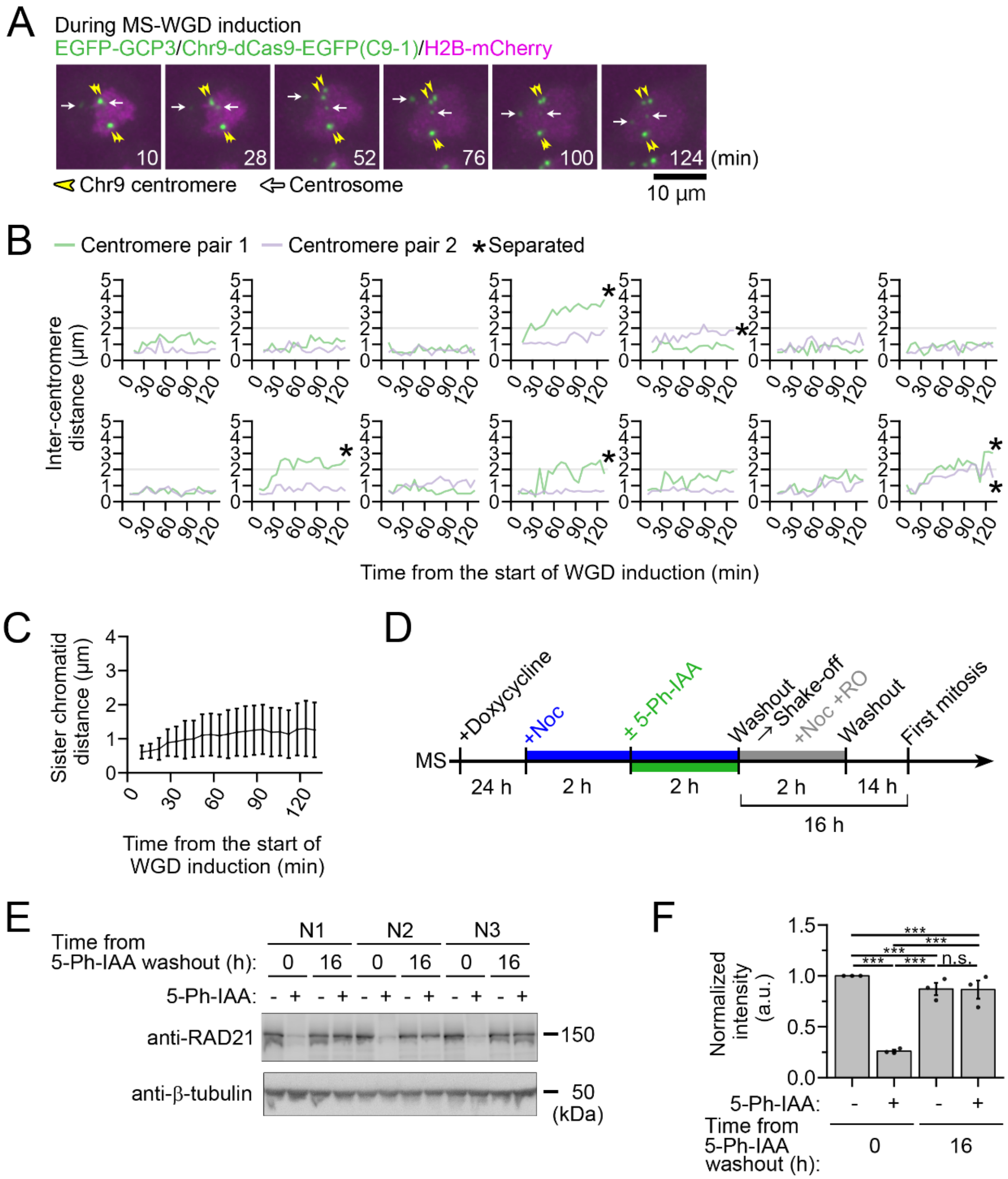

### Supplementary figure 7 (continued)

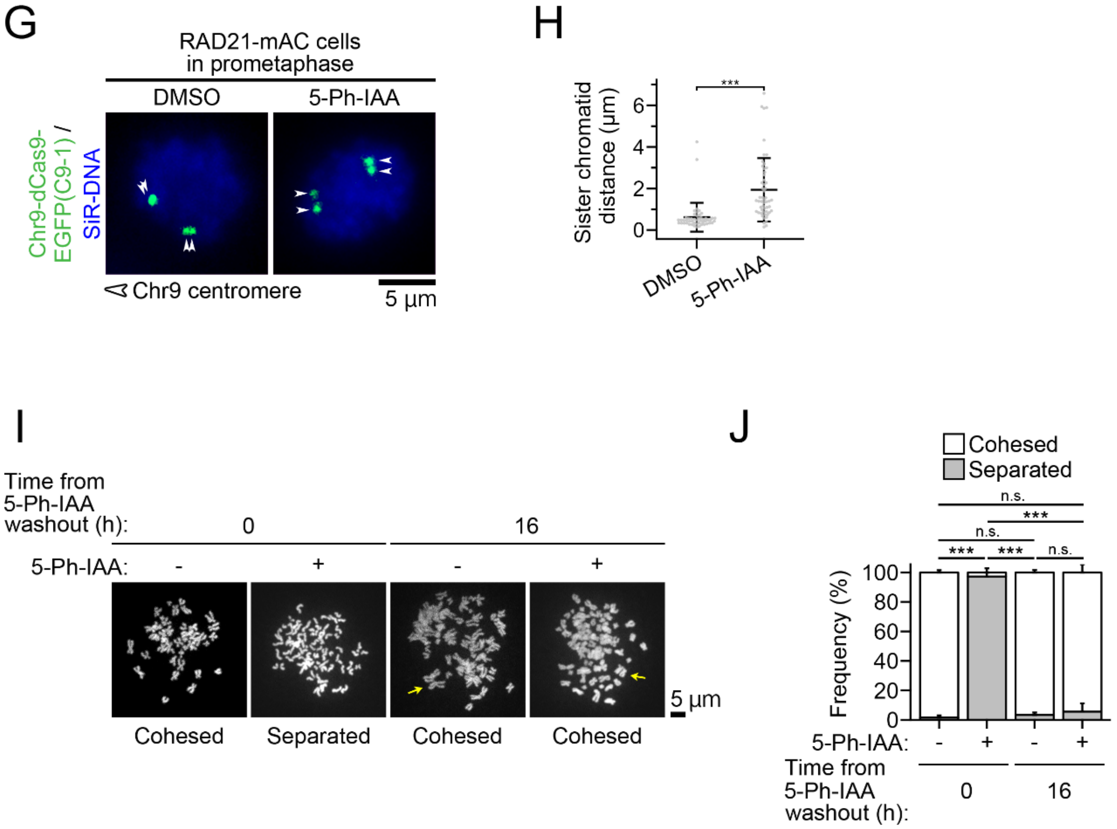

**Figure S7. Artificial induction of sister chromatid separation in a short time window upon MS-WGD by transient RAD21-mAC depletion**

**(A)** Time-lapse images of H2B-mCherry, EGFP-GCP3, and dCas9-EGFP with C9-1 sgRNA during MS-WGD. Images were taken at 6-min intervals. The timing of starting WGD induction was set to 0 min. White arrows and yellow arrowheads indicate the centrosomes and C9-1 centromeres, respectively. **(B, C)** Time-course of inter-sister chromatid distance during MS-WGD induction in A. Time-course data from individual cells in B. Asterisks indicate centromere pairs whose distances exceeded 2 μm at least once during the live-imaging analysis. Means ± s.d. of 28 sister chromatid pairs from 14 cells from two independent experiments in C. **(D)** A schematic image of the schedule for AID2-mediated transient depletion of RAD21 before MS-WGD induction. **(E)** Immunoblotting of RAD21 in RAD21-mAC cells treated with or without 5-Ph-IAA. RAD21 expression before or 16 h after washout of 5-Ph-IAA was tested. β-tubulin was detected as a loading control. **(F)** Quantification of the relative intensity of RAD21 in E. Protein loading differences were corrected based on β-tubulin signals. Mean ± s.e. of three independent experiments. Asterisks indicate statistically significant differences among conditions (n.s.: not significant, ***$p < 0.001$, the Steel-Dwass test). **(G)** Fluorescence microscopy of RAD21-mAC cells at prometaphase after 2-h nocodazole treatment with or without 5-Ph-IAA. The chromosome 9 centromeres and chromosomes were labeled by dCas9-EGFP expression with the C9-1 sgRNA and SiR-DNA staining, respectively. White arrowheads indicate C9-1 foci. **(H)** Inter-sister C9-1 foci distance in G. Means ± s.d. of at least 52 sister chromatid pairs from 26 cells from two independent experiments. Asterisks indicate statistically significant difference between conditions (***$p<0.001$, the Welch's t-test). **(I)** Chromosome spreads that were prepared from the cells undergoing MS-WGD (0 h) or the first mitosis after MS-WGD (16 h). While AID2-mediated depletion of RAD21 resulted in precocious separation of sister chromatids during MS-WGD, subsequent removal of 5-Ph-IAA allowed sister chromatid cohesion to be restored and maintained at the prometaphase in the first mitosis after MS-WGD. DNA was stained with SYTOX green. Note that diplochromosomes (marked by yellow arrows) are not prevalent in these spreads. All spread images are provided as Supplemental material S2. **(J)** Frequency of chromosome spreads with separated or cohesed sister chromatid pairs in I. Mean ± s.e. of three independent experiments. At least 53 cells were analyzed in each condition. Asterisks indicate statistically significant differences among conditions (n.s.: not significant, ***$p < 0.001$, the Fisher exact test).

### Supplementary figure 8

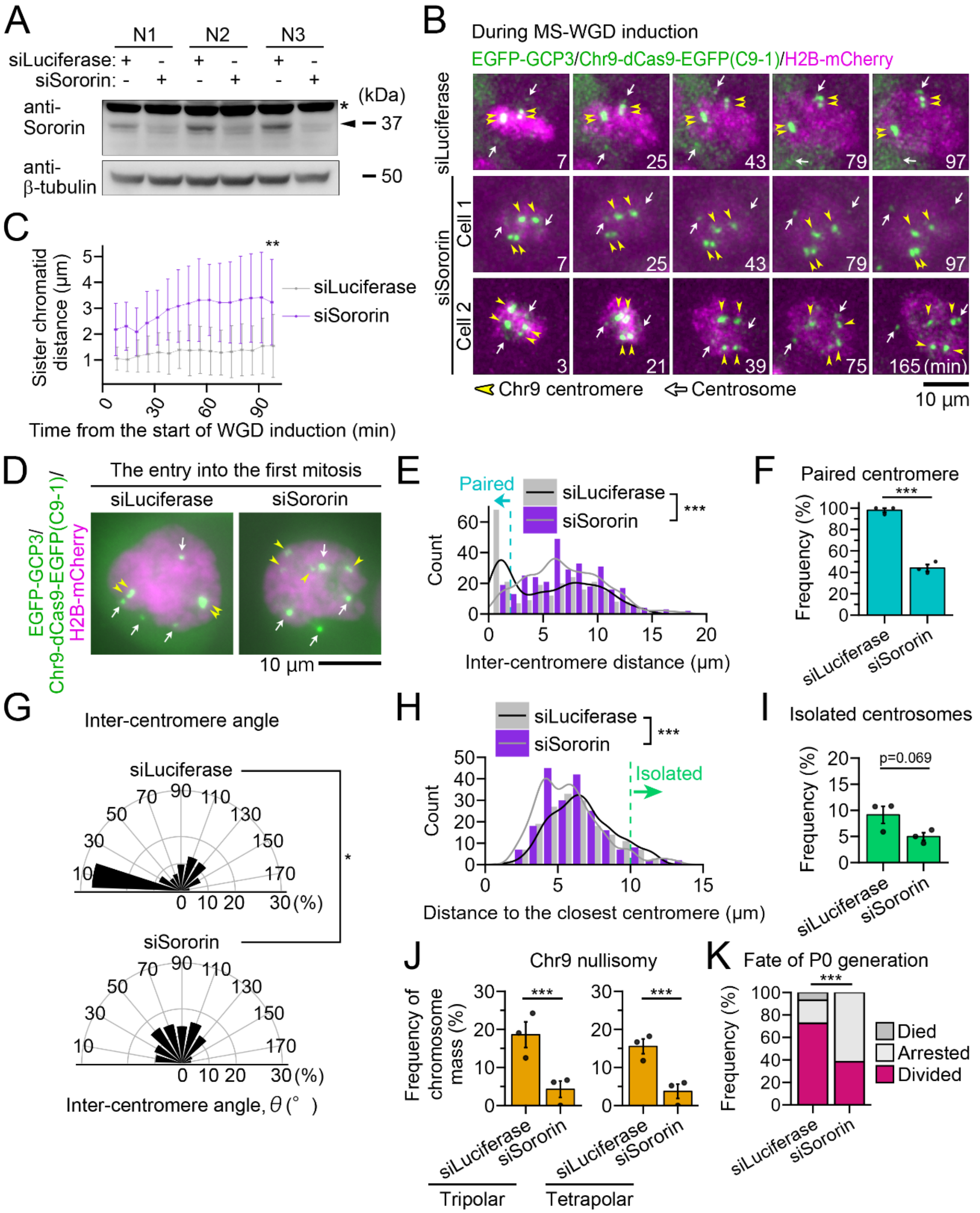

**Figure S8. Artificial induction of sister chromatid separation by RNAi-mediated Sororin depletion**

**(A)** Immunoblotting of Sororin in mock- or Sororin-depleted cells sampled at the timing of MS-WGD induction. β-tubulin was detected as a loading control. **(B)** Time-lapse images of H2B-mCherry, EGFP-GCP3, and dCas9-EGFP with C9-1 sgRNA during MS-WGD in mock- or Sororin-depleted cells. Images were taken at 6-min intervals. The timing of starting WGD induction was set as 0 min. White arrows and yellow arrowheads indicate the centrosomes and C9-1 centromeres, respectively. **(C)** Time course of inter-sister chromatid distance in B. Means ± s.d. of at least 14 sister chromatid pairs from at least 7 cells from two independent experiments. Asterisks indicate a statistically significant difference between conditions at the last time point (**$p$ < 0.01, the Welch's t-test). **(D)** Fluorescence microscopy of cells at the entry into the first mitosis after MS-WGD in mock- or Sororin-depleted cells. White arrows and yellow arrowheads indicate the centrosomes and C9-1 centromeres, respectively. **(E)** Histogram of inter-centromere distance in D. Six inter-centromere distances were obtained from the 4 C9-1 foci in an individual cell. At least 258 C9-1 combinations from 43 cells from three independent experiments were analyzed. Asterisks indicate a statistically significant difference between conditions (***$p$ < 0.001, the Welch's t-test). **(F)** Frequency of cells with paired C9-1 centromeres in D. Means ± s.e. of three independent experiments. Asterisks indicate a statistically significant difference between conditions (***$p$ < 0.001, the Brunner-Munzel test). **(G)** Quantification of the angles between each two C9-1 foci with regard to the centroid of the whole-chromosome mass in D. At least 216 C9-1 combinations from at least 36 cells from three independent experiments were analyzed. There is a statistically significant difference between conditions (*$p$ < 0.05, the Welch's t-test). **(H)** Histogram of the distance from each centrosome to its closest C9-1 focus in D. Four centrosome-centromere distances were obtained from an individual cell. At least 172 centrosome-centromere combinations from 43 cells from three independent experiments were analyzed. Asterisks indicate a statistically significant difference between conditions (***$p$ < 0.001, the Welch's t-test). **(I)** Frequency of isolated centrosomes in D. Means ± s.e. of three independent experiments. P-value obtained by the Brunner-Munzel test between conditions is shown on top. **(J)** Frequency of C9-1 nullisomic chromosome segregation during anaphase at the first mitosis after MS-WGD in mock- or Sororin-depleted cells. Means ± s.e. of three independent experiments. At least 102 segregated chromosome masses from at least 34 cells from three independent experiments were analyzed in each polarity. Asterisks indicate statistically significant differences between conditions (***$p$ < 0.001, the Brunner-Munzel test). **(K)** Fates of P0 cells (the first cell cycle) after MS-WGD induction in mock- or Sororin-depleted cells. At least 75 cells from two independent experiments were analyzed. Asterisks indicate a statistically significant difference between conditions (***$p$ < 0.001, the Fisher exact test).

### Supplementary figure 9

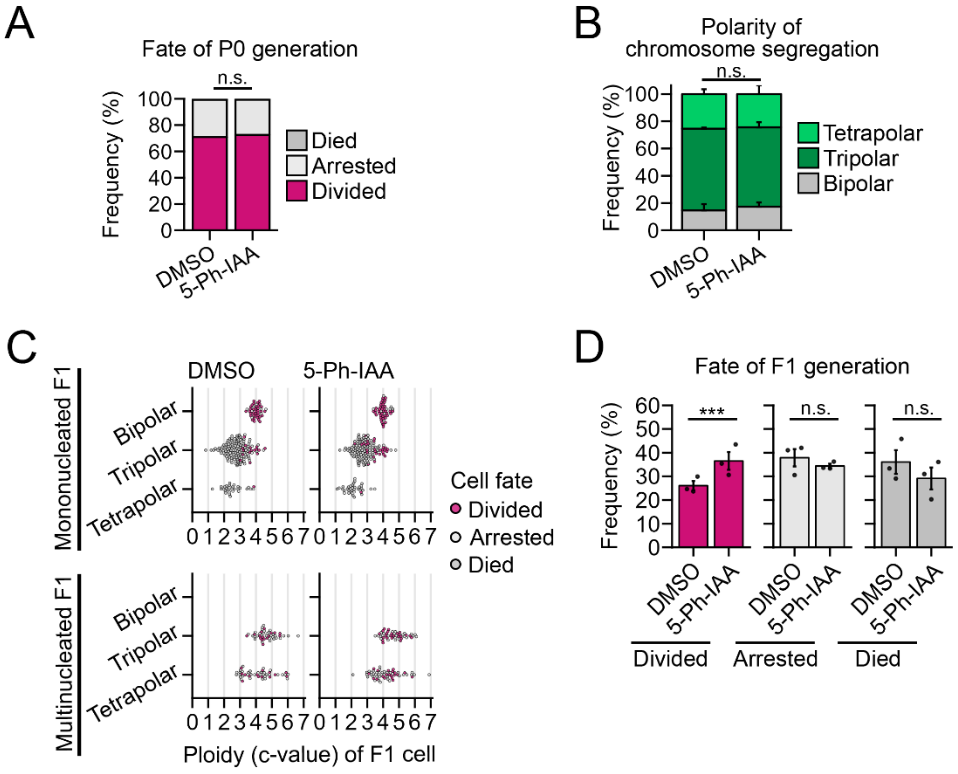

### Supplementary figure 9 (continued)

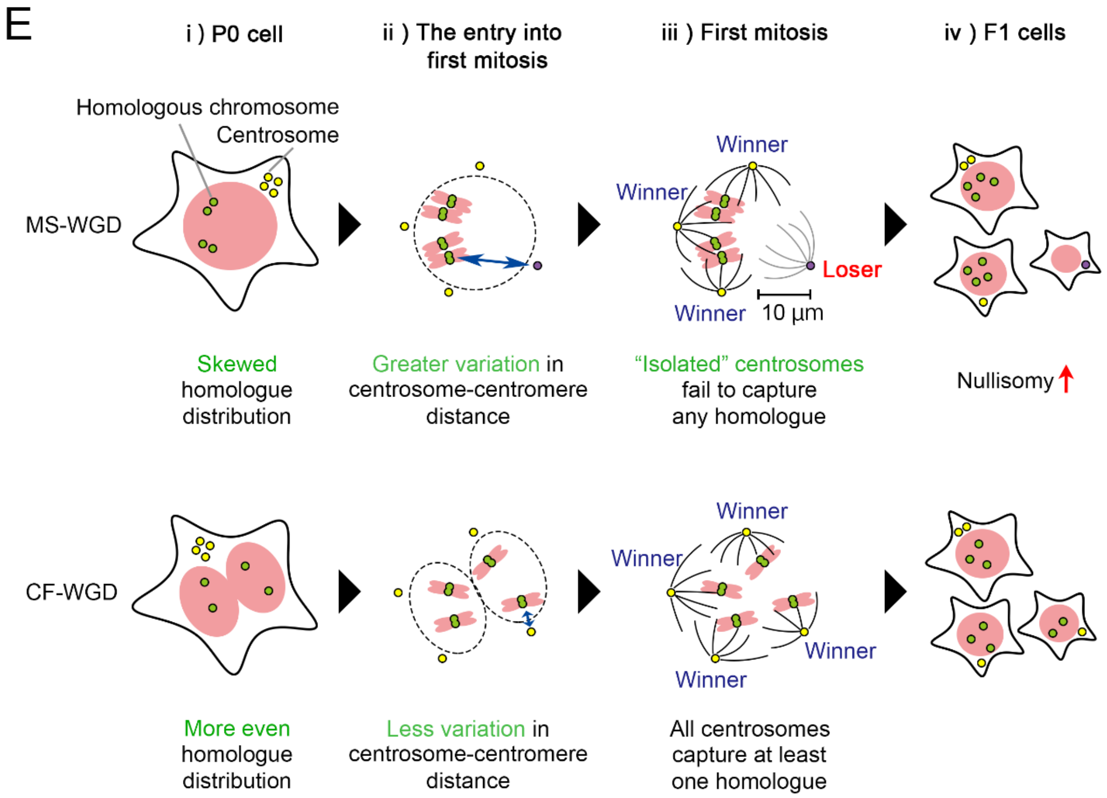

**Figure S9. Artificial induction of sister chromatid separation by transient RAD21-mAC depletion improves F1 cell viability after MS-WGD**

**(A-D)** Fates of P0 cells (A), frequency of chromosome segregation polarity at first mitosis (B), estimated DNA content (c-value) in individual F1 cells (C), or viability of F1 cells (D) after MS-WGD induction in RAD21-mAC cells treated with or without 5-Ph-IAA. Means ± s.e. of three independent experiments in B or D. At least 206 cells (A), 133 cells (B), and 317 cells (C, D) from three independent experiments were analyzed. There is no statistically significant difference between conditions in A or B ($p$ = 0.80 or $p$ = 0.93, respectively, the Fisher exact test). Asterisks indicate a statistically significant difference between conditions in D (***$p < 0.001$, the Brunner-Munzel test). **(E)** A model for the route-dependent difference in post-WGD progenies. While sufficient separation of sister chromatids establishes even distribution of homologous chromosomes in the first cell cycle after CF-WGD, insufficient sister chromatid separation causes more skewed homologue distribution after MS-WGD (i). This difference in homologue arrangement results in greater variation in centrosome-centromere distance upon entry into the first mitosis after MS-WGD than CF-WGD (ii). Presumably because of the limited reach length of centrosomal microtubules, centrosomes apart from any of homologous centromeres further than 10 μm have a high chance of failure in capturing any of these homologues ("Loser" centrosome, iii). These geometric characteristics of post-MS-WGD cells cause frequent nullisomy in F1 cells (iv), limiting the viability of post-WGD progenies compared to CF-WGD.

### Supplementary figure 10

Immunoblotting associated with supplementary figure 2B

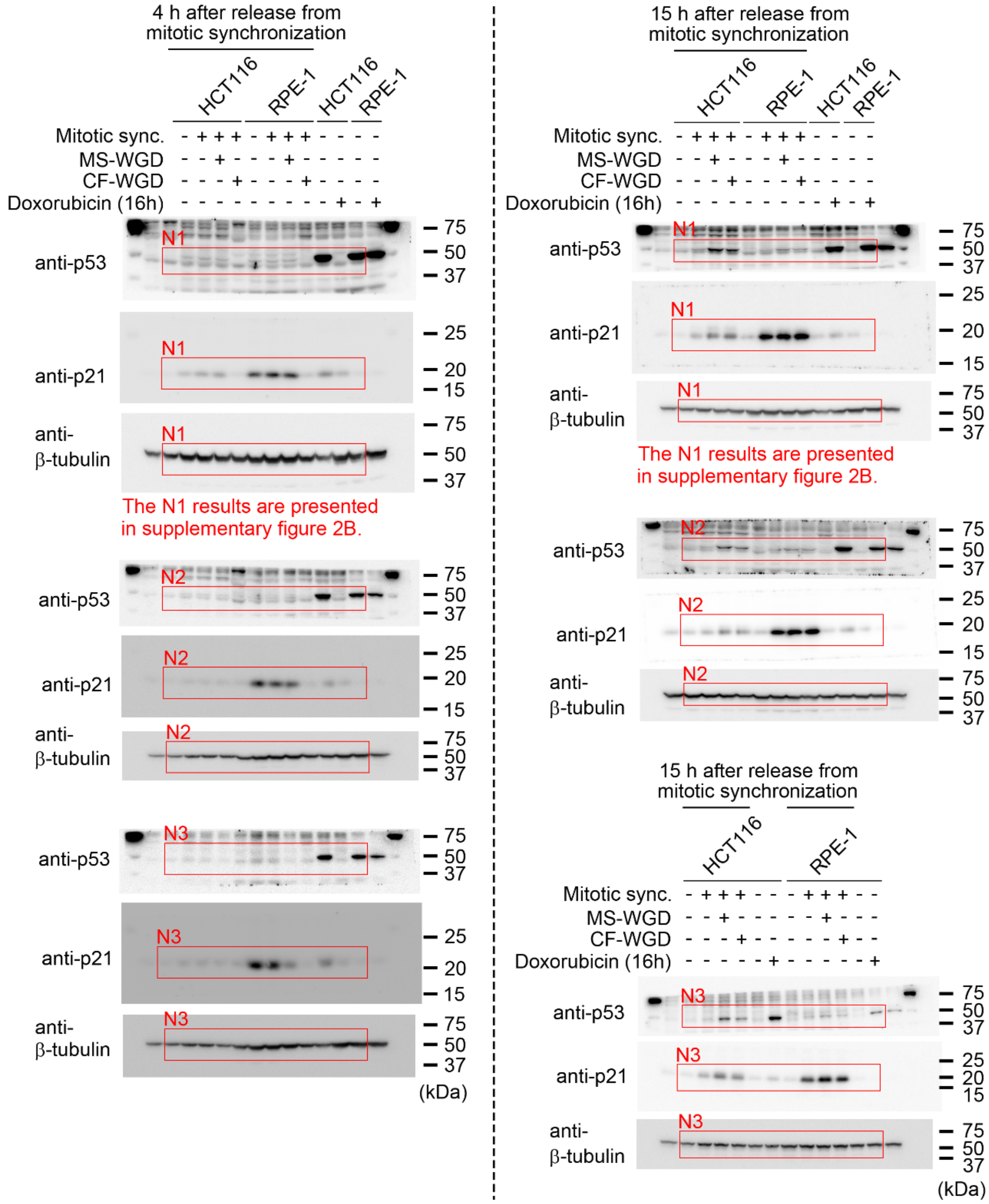

### Supplementary figure 10 (continued)

Immunoblotting associated with supplementary figure 2G

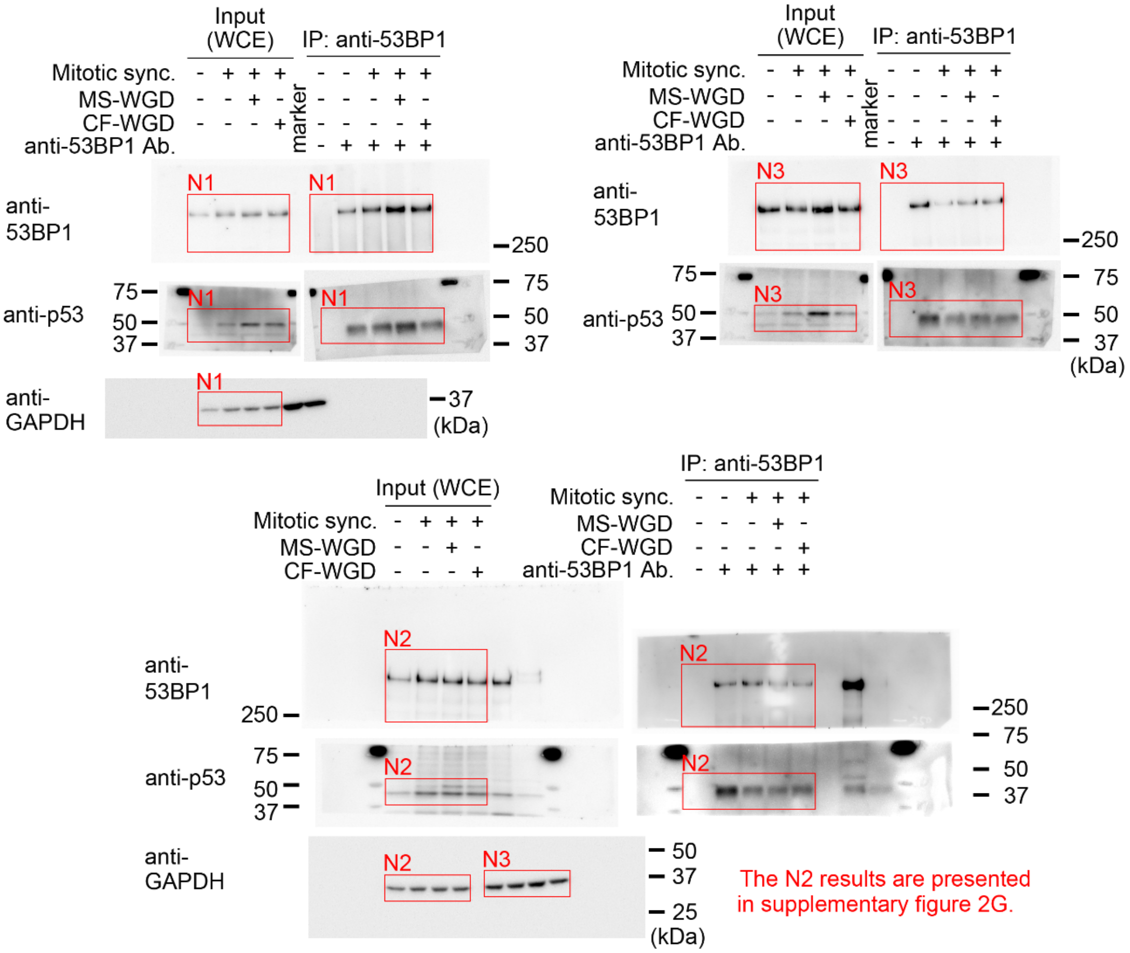

### Supplementary figure 10 (continued)

Immunoblotting associated with supplementary figure 4E

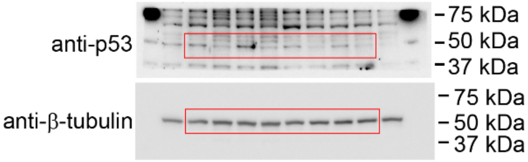


Immunoblotting associated with supplementary figure 7E

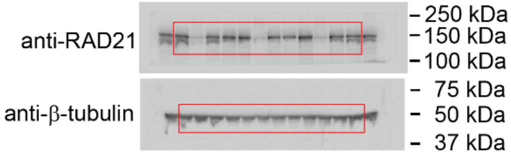


Immunoblotting associated with supplementary figure 8A

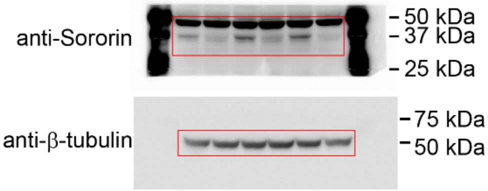

**Figure S10. Original immunoblot membranes used for the analyses**
Uncropped images of all immunoblots are shown, with the approximate indications of the cropped regions.

**Datasets**
Datasets S1-4 were provided as separate supplementary spreadsheets.

**Dataset S1. List of cell lines used in this study**

**Dataset S2. List of plasmids used in this study**

**Dataset S3. List of antibodies and compounds used in this study**

**Dataset S4. Numerical values assigned to the models**

**Movie S1 (separate file). The dynamics of homologous centromeres and the centrosomes during the first mitosis after MS-WGD**
Fluorescence microscopy (maximum z-projections of confocal images) of the first mitosis after MS-WGD. The centrosomes, centromeres of chromosome 9, and chromosomes were labeled by EGFP-EGP3, dCas9-EGFP with specific sgRNA (C9-1), and H2B-mCherry, respectively. Individual centrosomes or centromeres were distinguished by labeling different letters (a-d) or numbers (1-4), respectively. The movie is shown at 600x real-time. Time is indicated as hour: min. Scale bar: 10 μm.

**Supplemental materials (separate files).**

**Supplemental material S1. DNA contents in all colonies analyzed in Figure 1**
Histograms of DNA content distribution in all colonies analyzed for Fig. 1F and G. DNA contents were quantified based on Hoechst33342 signal in each cell within the colonies. Blue lines indicate the estimated G1 DNA content values.

**Supplemental material S2. All microscopic images of the chromosome spreading**
A panel of all microscopic images of the chromosome spreading in Fig. S7I. See the corresponding figure legend for details.

**Supplemental material S3. The ImageJ macro files used for the automated segmentation**
