## Supplementary material for "Sister chromatid separation determines the proliferative properties upon whole-genome duplication via homologous chromosome arrangement": SI appendix: 230811 shading correct_flip_230810colony.ijm

```
d = getDirectory("Choose a Directory");		
newfolder = d + "/shading corrected/";
File.makeDirectory(newfolder);
files = getFileList(d);	
for (i = 0; i < files.length; i++){
	f = d + files[i];
	doMultiFiles(f);
}

function doMultiFiles(f){
	if(!File.isDirectory(f)){
		print(f);
open(f);
		wait(100);
		
		Stack.setSlice(1);
		run("Duplicate...", " ");
		wait(100);
		
		
run("Shading Corrector");
		wait(100);
		
		run("Flip Vertically", "stack");
		run("Flip Horizontally", "stack");
		wait(100);
		

		name = getTitle();
		extension = indexOf(name, ".");
		sub = substring(name, 0, extension);
		newpath = newfolder + sub + "_sc.tif";
		saveAs(".tif", newpath);
		
		close();
		close();
		//run("Close All");
	}
}
```
