## Supplementary material for "Sister chromatid separation determines the proliferative properties upon whole-genome duplication via homologous chromosome arrangement": SI appendix: 250408_ColonyPloidyMeasure_Auto_background_macro1.ijm

```
savefolder = getDirectory("Choose a Directory");		
n = roiManager("count");
run("Set Scale...", "distance=0.6 known=1 unit=µm global");
run("Set Measurements...", "area mean centroid perimeter shape integrated stack display redirect=None decimal=3");

for (i = 0; i < n; i++) {
    roiManager("select", i);
    roiName = RoiManager.getName(i);                 // 配列からROI名を取得
    run("Duplicate...", "title=" + roiName);
    //ColonyRoi = getTitle();  //get colony roi title
    
    newpath = savefolder + roiName + ".tif";
    saveAs(".tif", newpath);
    close();
}
```
