## Supplementary material for "Sister chromatid separation determines the proliferative properties upon whole-genome duplication via homologous chromosome arrangement": SI appendix: 250408_ColonyPloidyMeasure_Auto_background_macro2.ijm

```
showMessage("Select Open Folder");
openDir = getDirectory("Choose a Directory");
saveDir = openDir + "/csv/";
File.makeDirectory(saveDir);

list = getFileList(openDir);

Day = "250405_N5"
Sample = "NR_well2"

for (i=0; i<list.length;i++){
	operation();
};

function operation(){
	open(openDir+list[i]);
    ColonyRoi = getTitle();  //get colony roi title
    
    run("Set Scale...", "distance=0.6 known=1 unit=µm global");
    run("Set Measurements...", "area mean centroid perimeter shape integrated stack display redirect=None decimal=3");
    run("Measure");
    
    saveAs("Results", saveDir + Day + "_" + Sample + "_" +ColonyRoi + ".csv");
    run("Clear Results");
    run("Close All");
}
```
