## Supplementary material for "Sister chromatid separation determines the proliferative properties upon whole-genome duplication via homologous chromosome arrangement": SI appendix: 250408_ColonyPloidyMeasure_Auto_macro2.ijm

```
showMessage("Select Open Folder");
openDir = getDirectory("Choose a Directory");
saveDir = openDir + "/csv/";
File.makeDirectory(saveDir);

list = getFileList(openDir);

Day = "250405_N5"
Sample = "NR_well2"

for (i=0; i<list.length;i++){
	operation();
};

function operation(){
	open(openDir+list[i]);
    ColonyRoi = getTitle();  //get colony roi title
    
    run("Set Scale...", "distance=0.6 known=1 unit=µm global");
    run("Set Measurements...", "area mean centroid perimeter shape integrated stack display redirect=None decimal=3");
    
    run("Command From Macro","command=[de.csbdresden.stardist.StarDist2D],args=['input':'" + ColonyRoi + "', 'modelChoice':'Versatile (fluorescent nuclei)', 'normalizeInput':'true', 'percentileBottom':'1.0', 'percentileTop':'99.8', 'probThresh':'0.5', 'nmsThresh':'0.4', 'outputType':'Both', 'nTiles':'1', 'excludeBoundary':'2', 'roiPosition':'Automatic', 'verbose':'false', 'showCsbdeepProgress':'false', 'showProbAndDist':'false'], process=[false]");
    selectWindow("Label Image");
    close();

    selectWindow(ColonyRoi);
    roiManager("Show All");
    roiManager("measure");
    
    saveAs("Results", saveDir + Day + "_" + Sample + "_" +ColonyRoi + ".csv");
    run("Clear Results");
    run("Close All");
}


//get xy from raw image
/*run("Set Scale...", "distance=0.6 known=1 unit=µm global");
run("Set Measurements...", "centroid stack redirect=None decimal=3");
run("Measure");
X = round(getResult("X",0));
Y = round(getResult("Y",0));
run("Clear Results");
*/
```
