## Supplementary material for "Sister chromatid separation determines the proliferative properties upon whole-genome duplication via homologous chromosome arrangement": SI appendix: README.txt

README


> 230811 shading correct_flip_230810colony.ijm
ImageJ macro to correct shading.
Place your data, acquired using the Grid/Collection Stitching plugin, into a folder. Use this macro first for the acquired image.

> 250408_ColonyPloidyMeasure_Auto_background_macro1.ijm
> 250408_ColonyPloidyMeasure_Auto_background_macro2.ijm
ImageJ macros for measuring the background.
Open your stitched data of a well without cells. Open the ROI manager of each colony. 
Run "250408_ColonyPloidyMeasure_Auto_background_macro1.ijm" for cropping the position of each colony.
Run "250408_ColonyPloidyMeasure_Auto_background_macro2.ijm" for measuring the background intensity of each colony area.

> 250408_ColonyPloidyMeasure_Auto_macro1.ijm
> 250408_ColonyPloidyMeasure_Auto_macro2.ijm
ImageJ macros for measuring the Hoechst intensity of each colony.
Open your stitched data of a well after colony formation. Open the ROI manager of each colony. 
Run "250408_ColonyPloidyMeasure_Auto_macro1.ijm" for cropping the position of each colony.
Run "250408_ColonyPloidyMeasure_Auto_macro2.ijm" for measuring the Hoechst intensity of each nucleus in each colony.

The value from Hoechst intensity was divided by the background intensity to correct for the background.
