## Supplementary figures and images for "Sister chromatid separation determines the proliferative properties upon whole-genome duplication via homologous chromosome arrangement"

### S1_CF_N1_well1.png

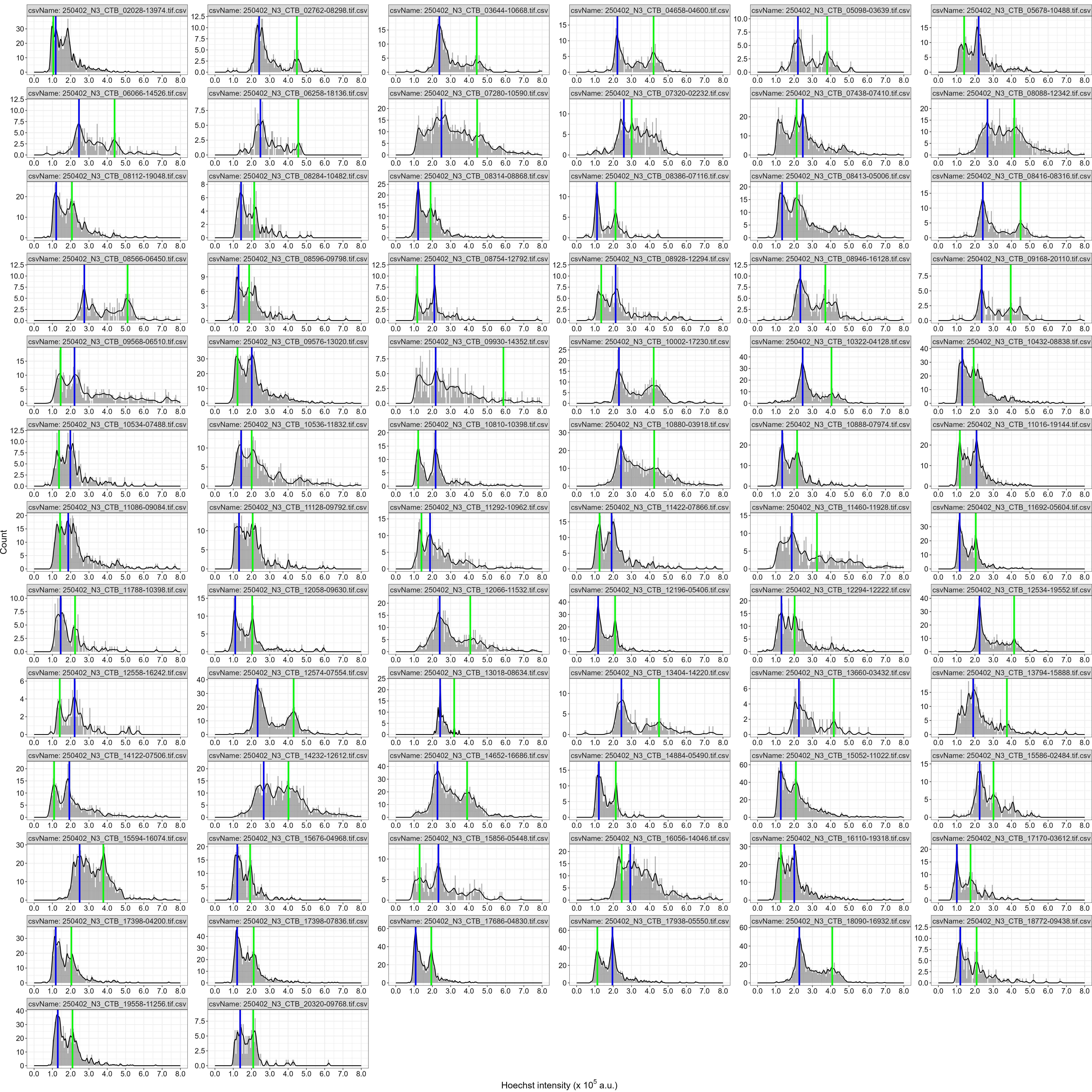

### S1_CF_N1_well2.png

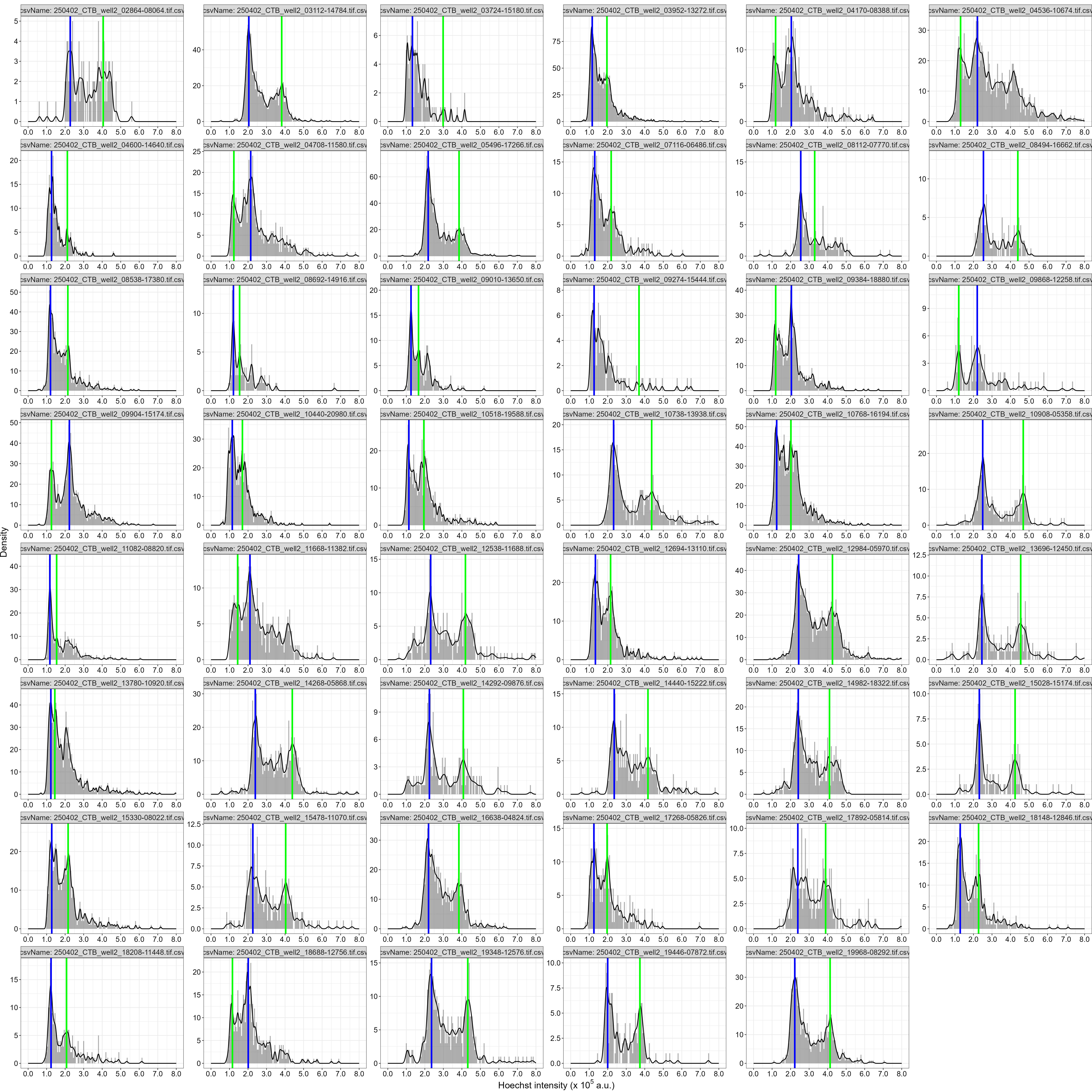

### S1_CF_N2_well1.png

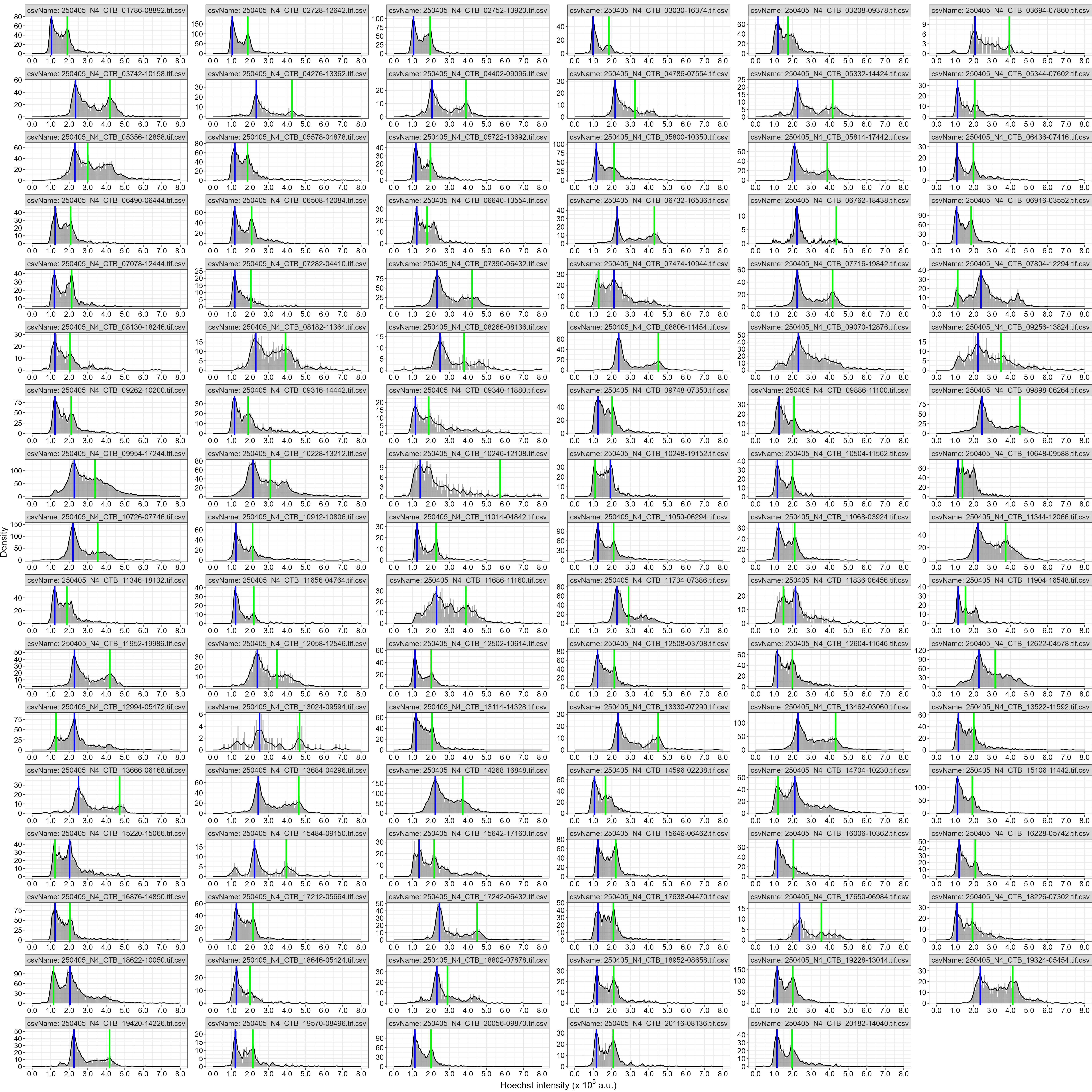

### S1_CF_N2_well2.png

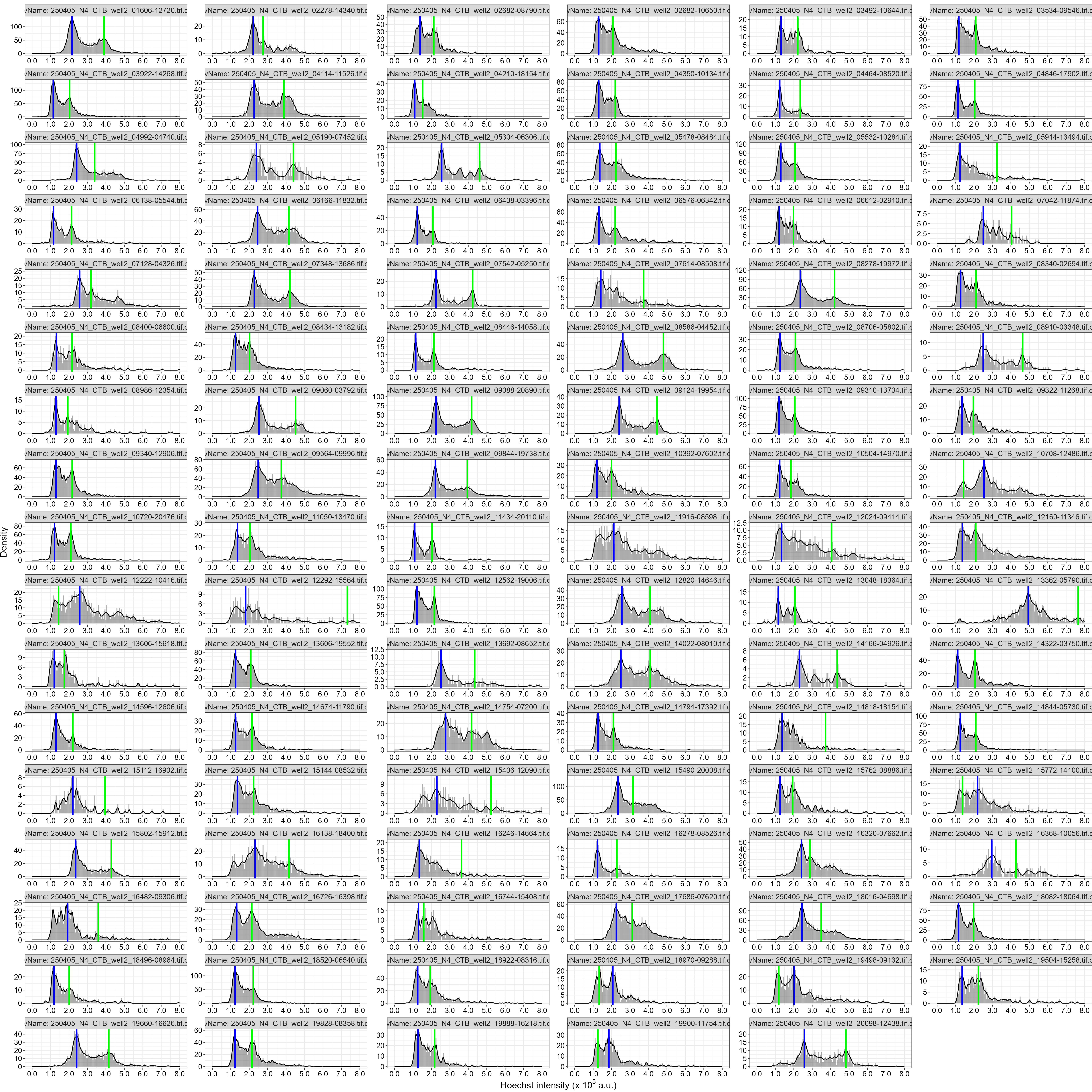

### S1_CF_N3_well1.png

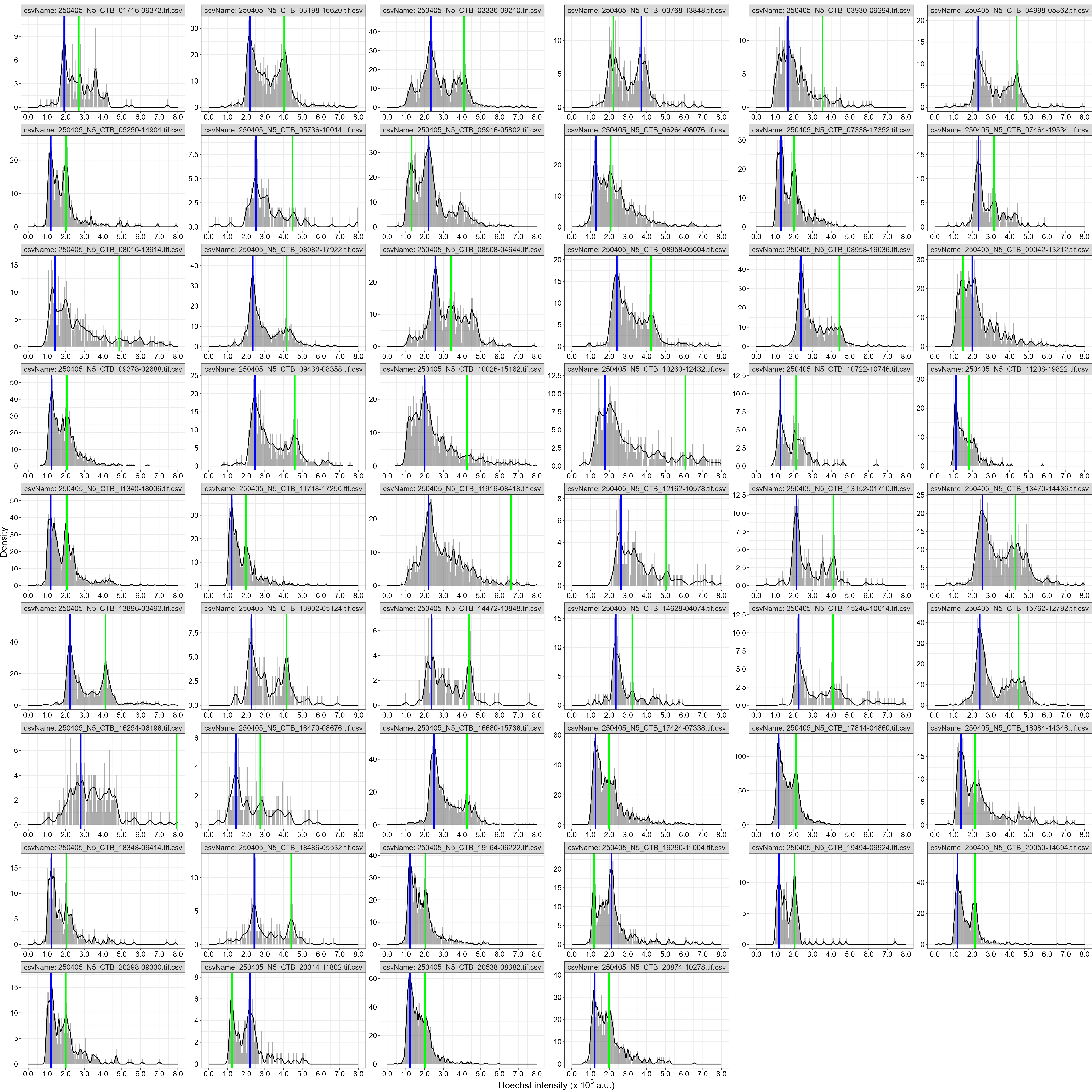

### S1_CF_N3_well2.png

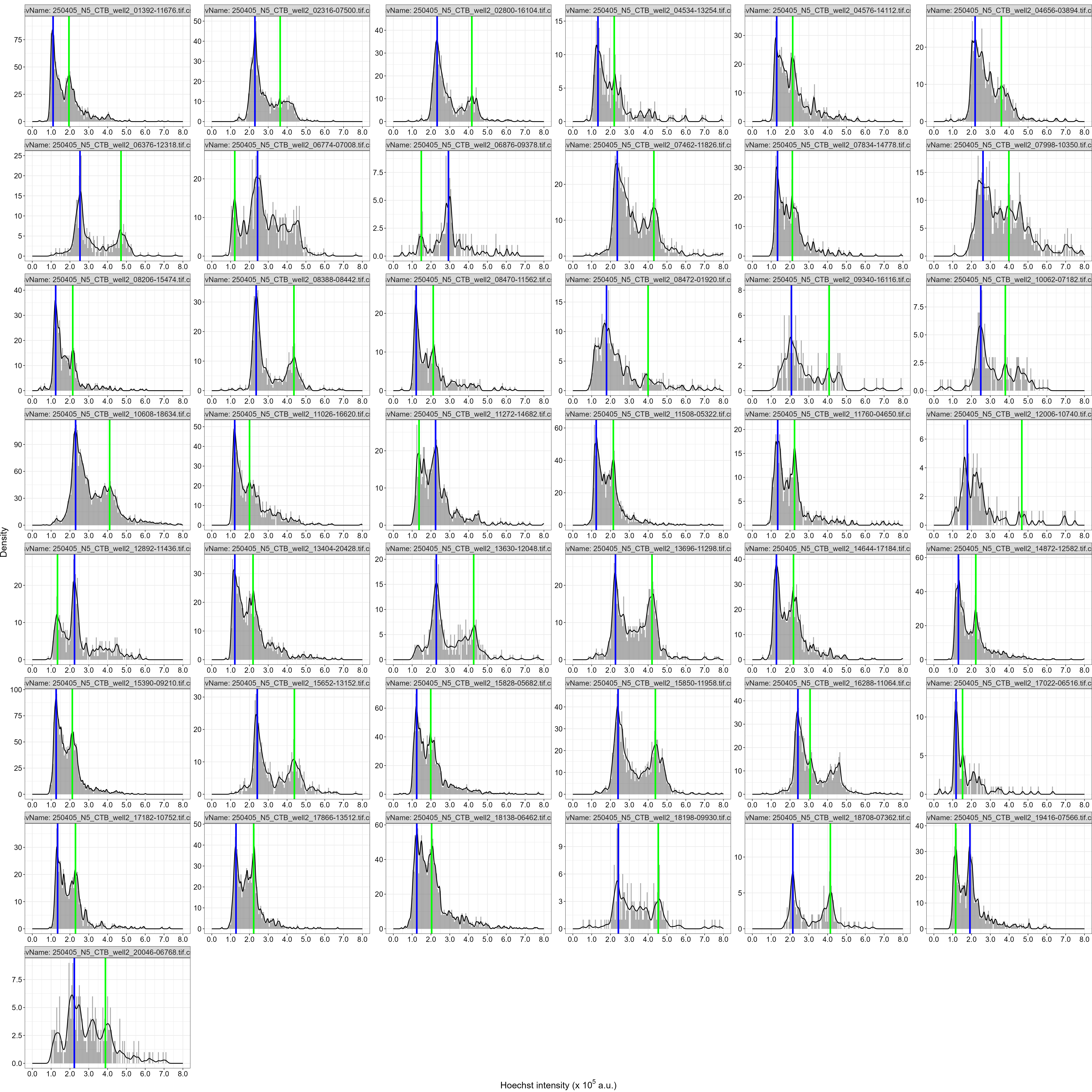

### S1_MS_N1_well1.png

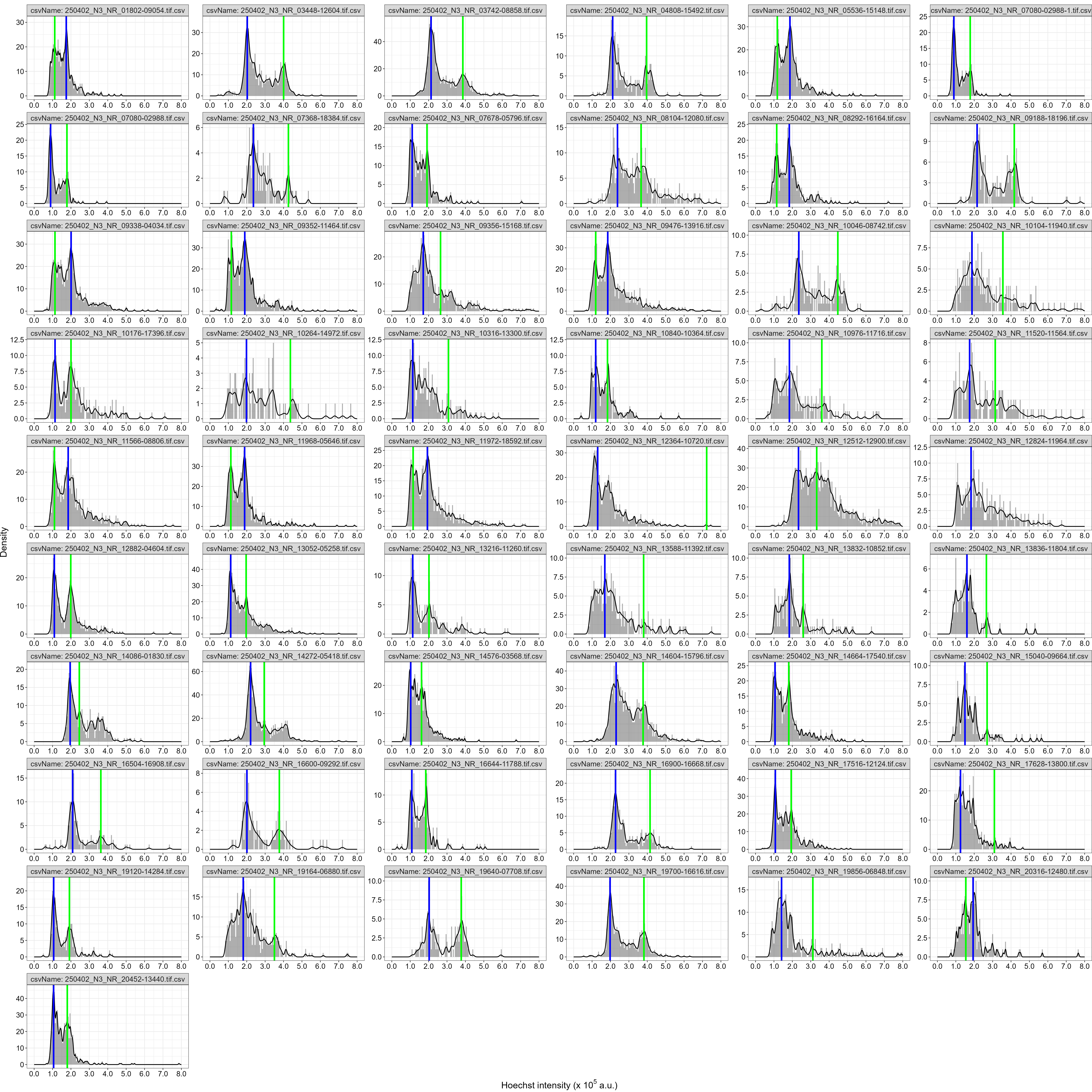

### S1_MS_N1_well2.png

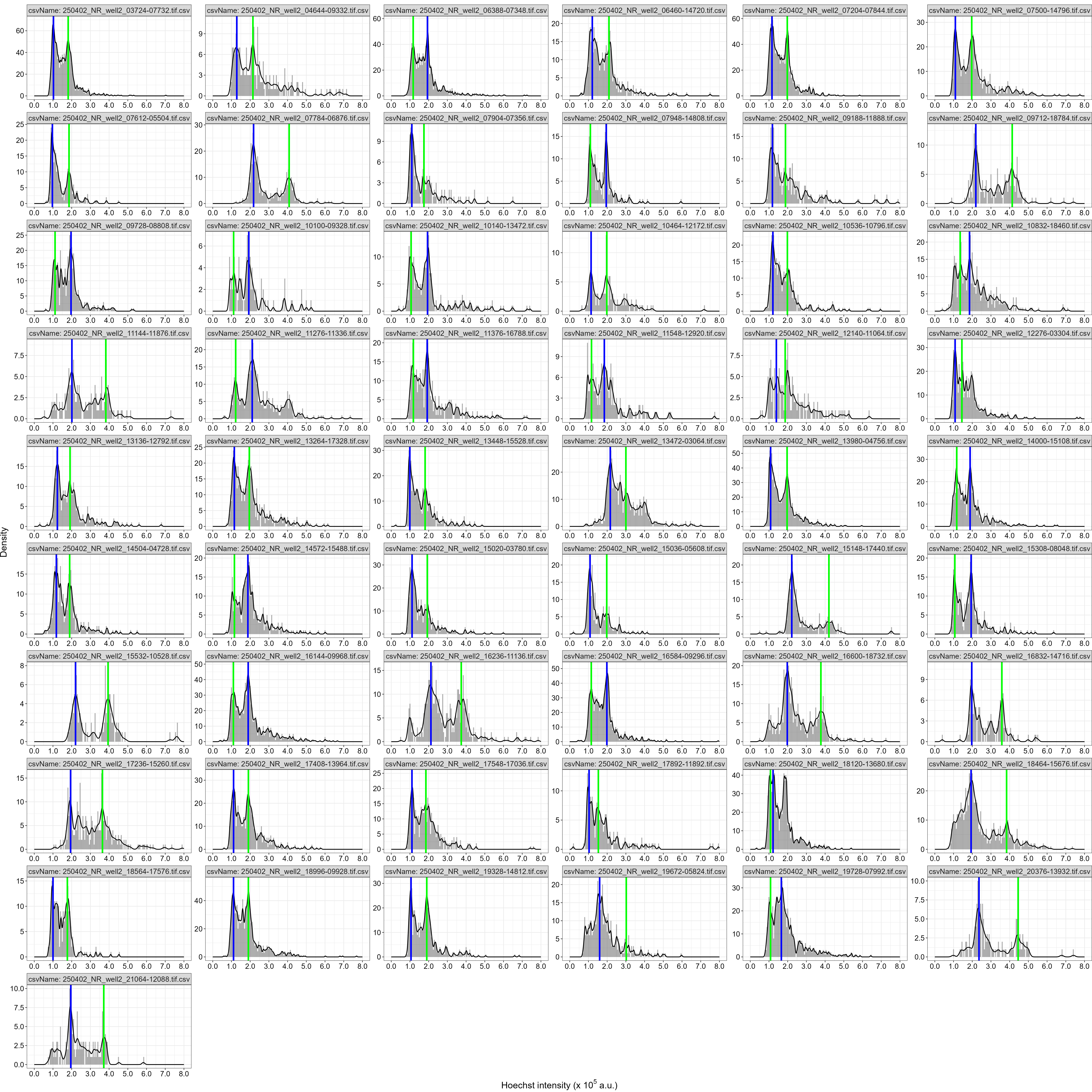

### S1_MS_N2_well1.png

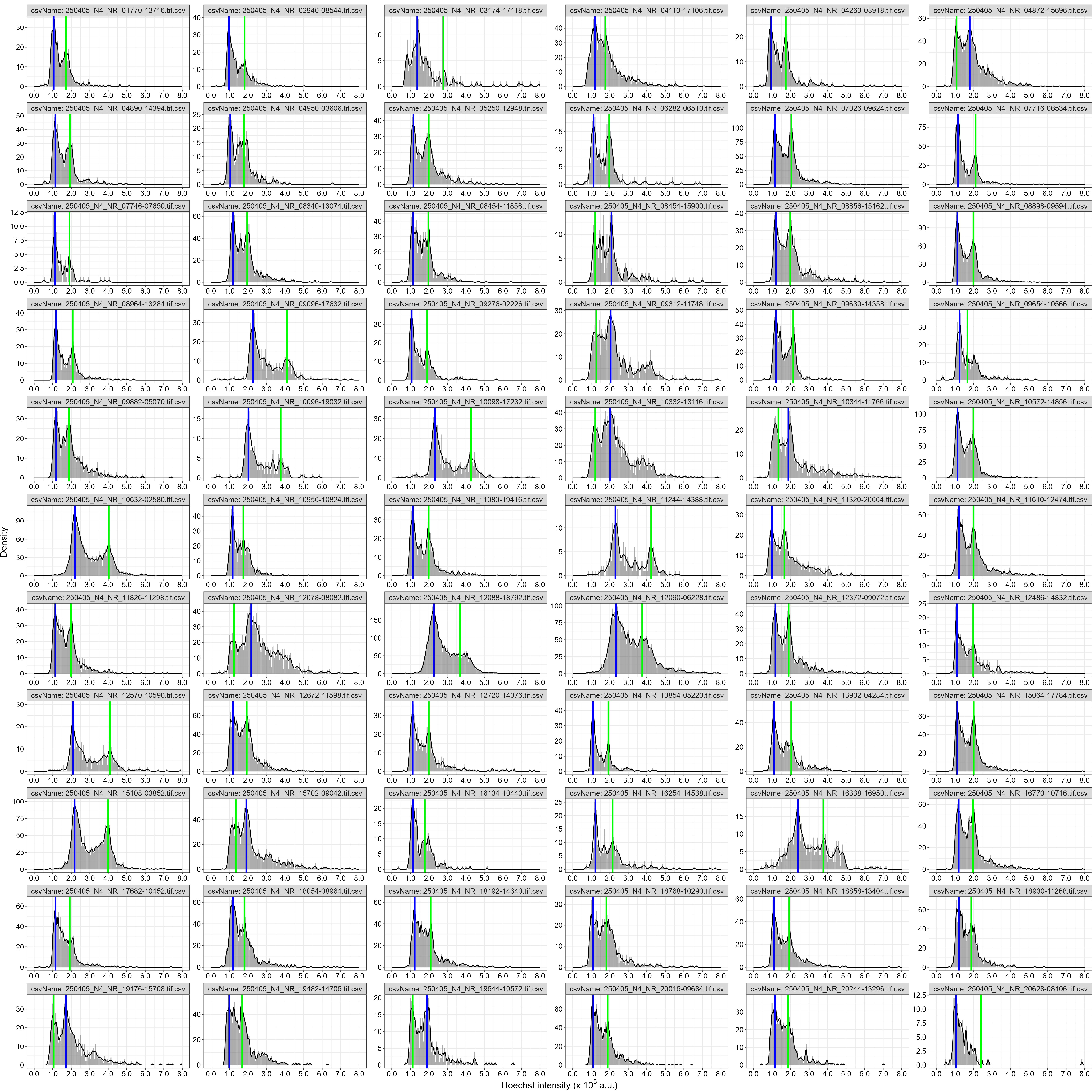

### S1_MS_N2_well2.png

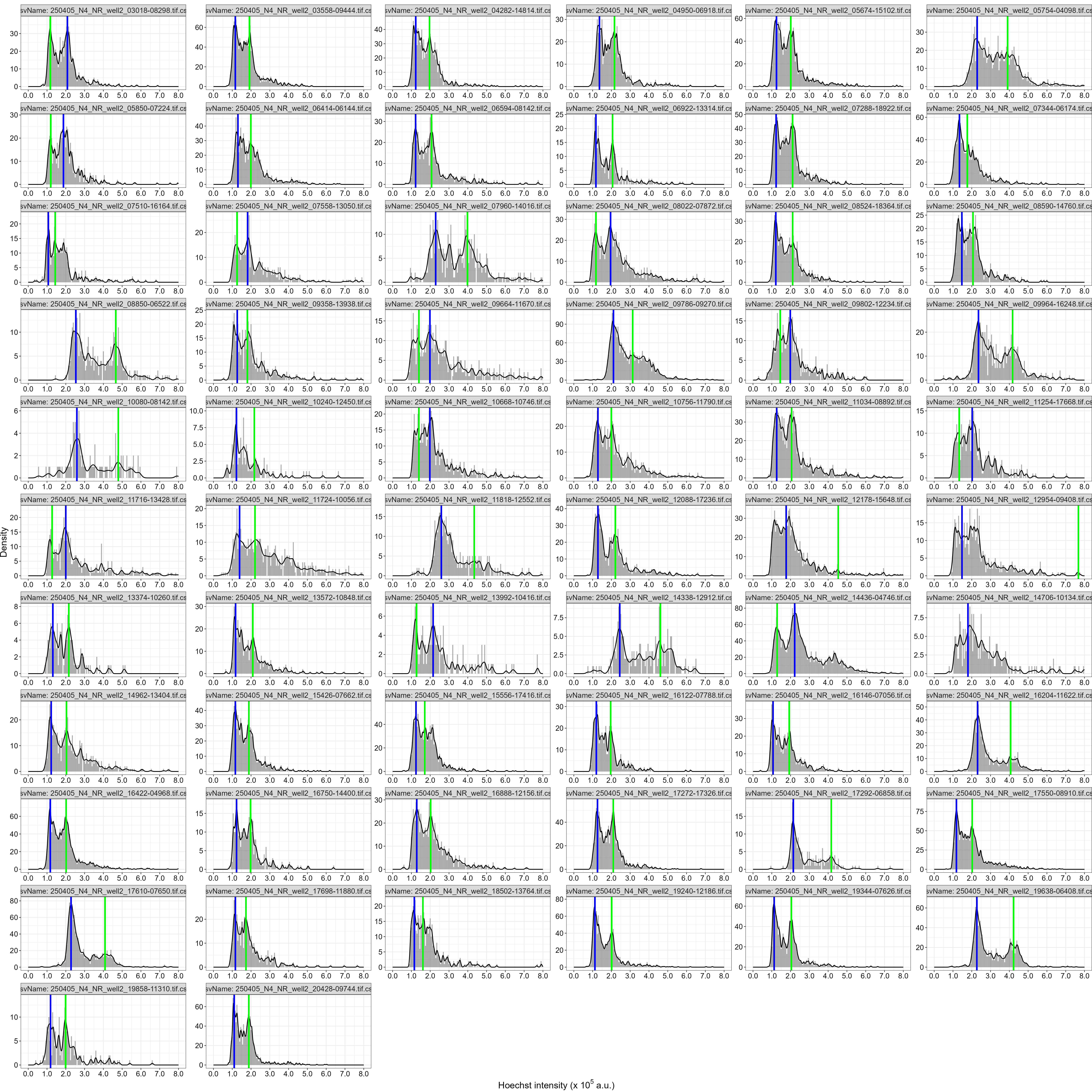

### S1_MS_N3_well1.png

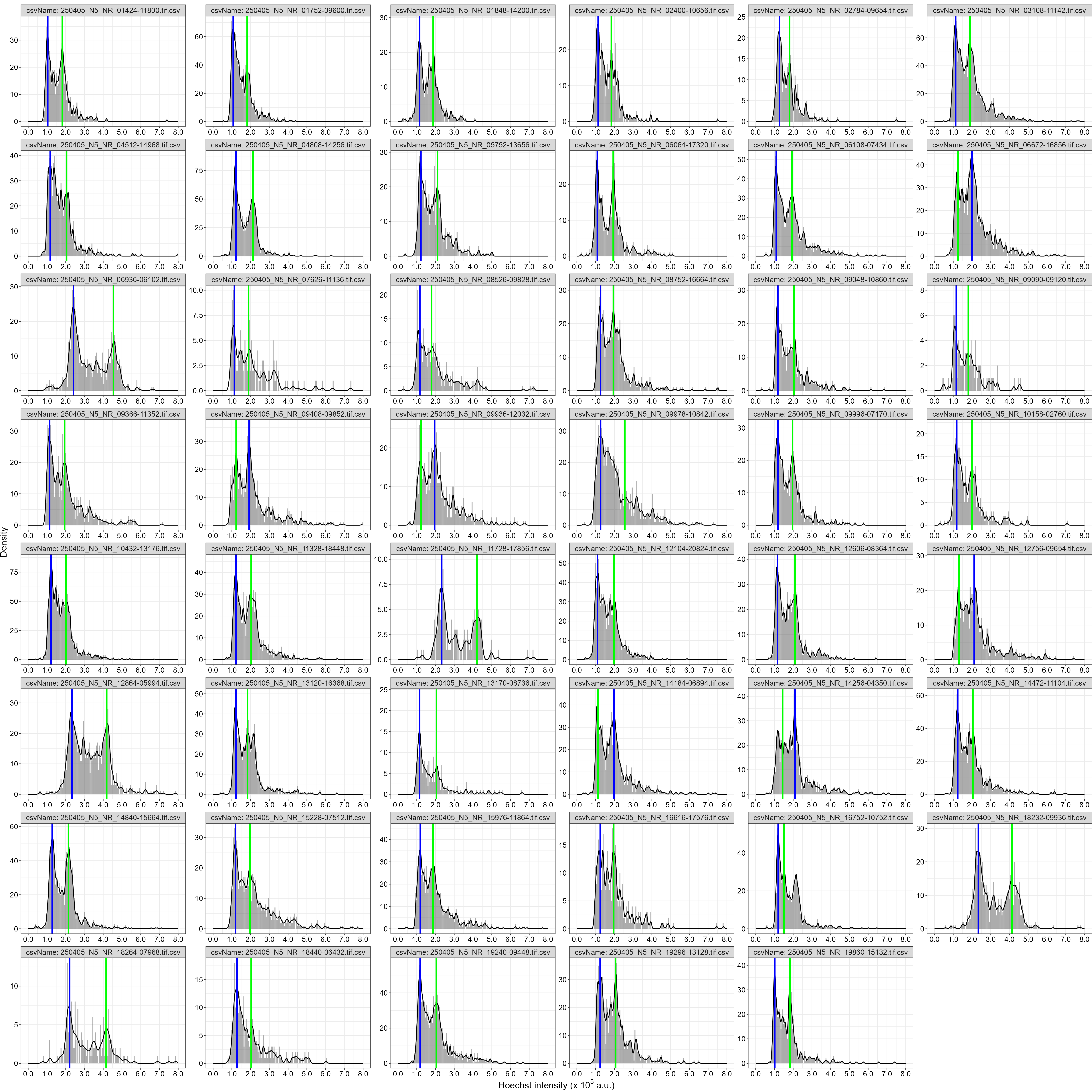

### S1_MS_N3_well2.png

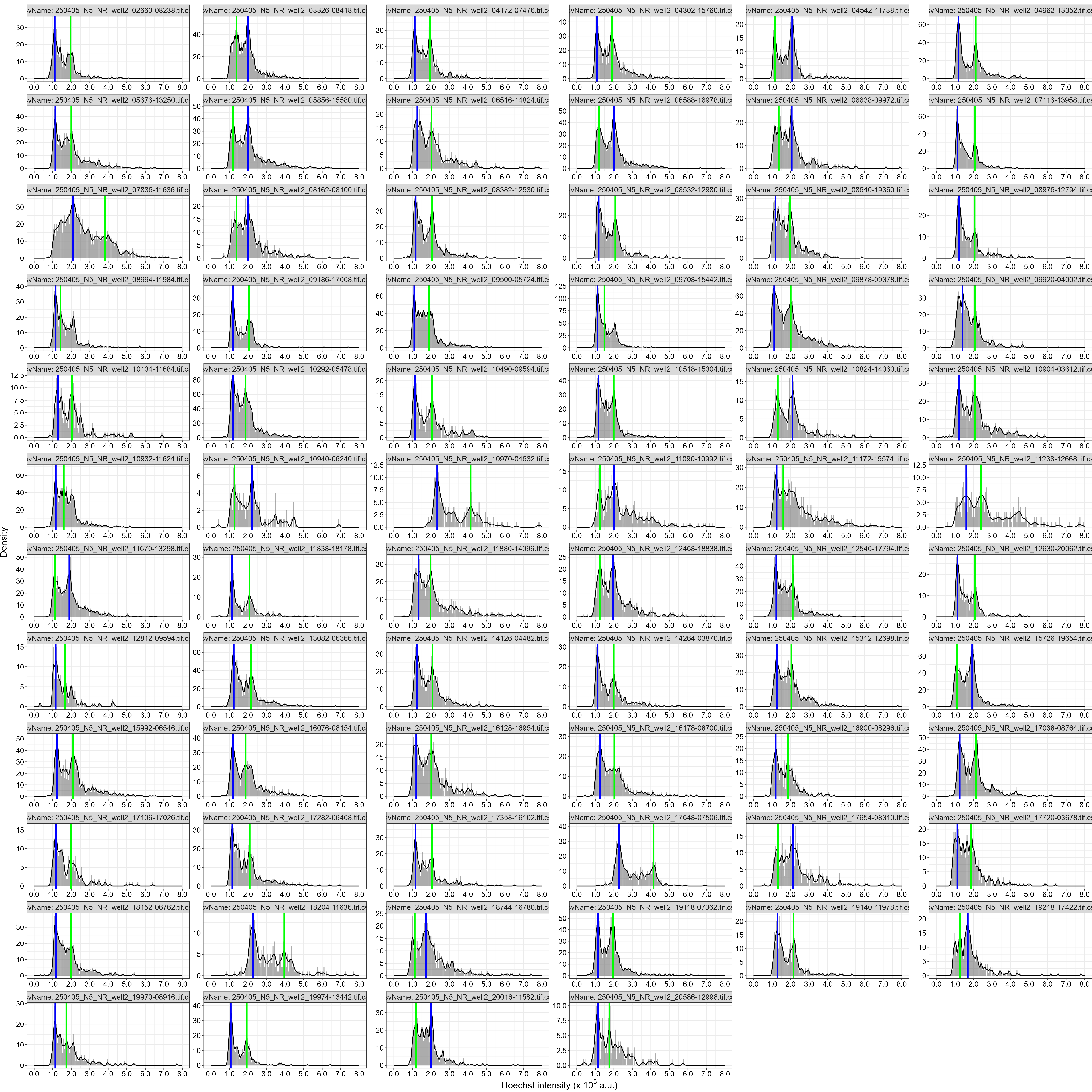

### Supplemental_material_S2_N1_0h_251216.png

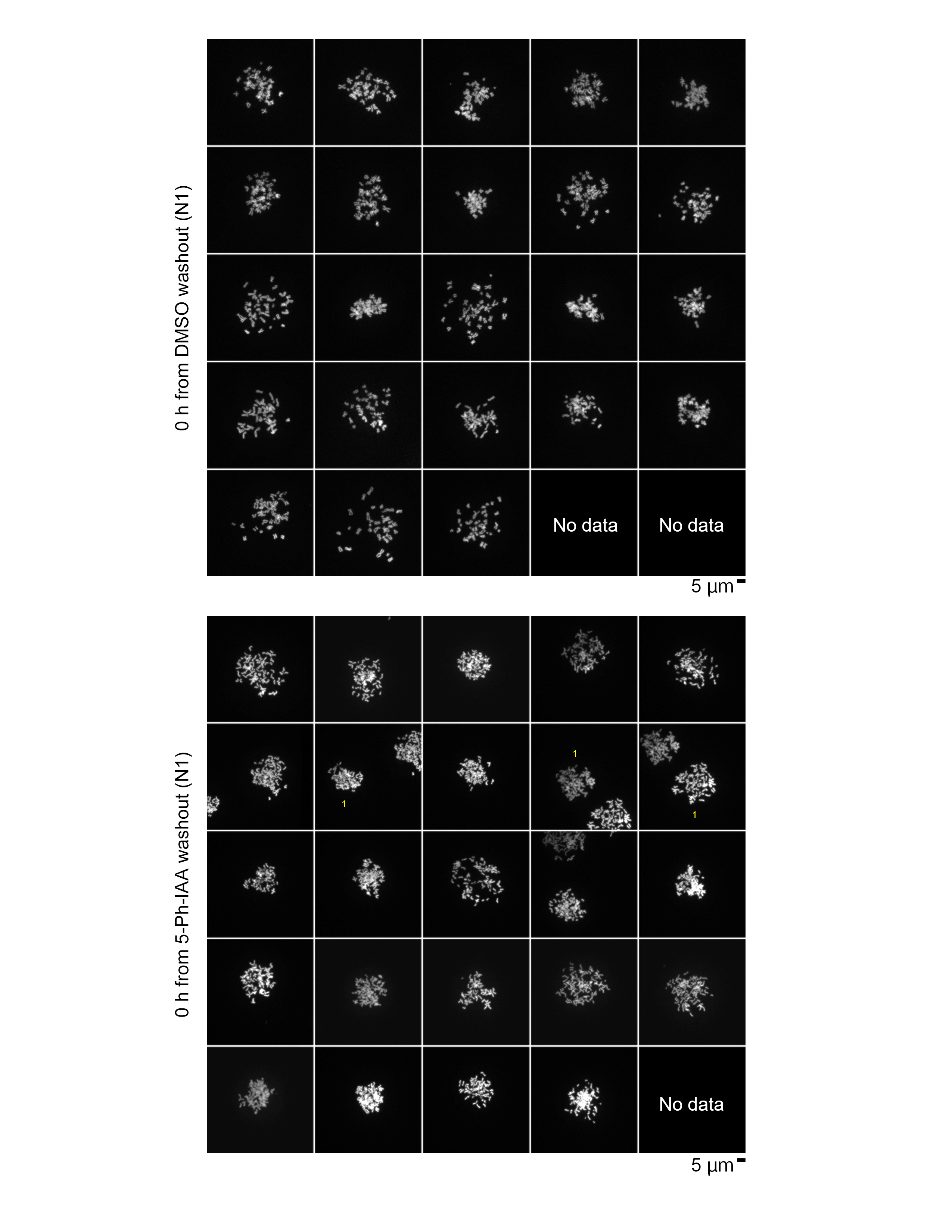

### Supplemental_material_S2_N1_16h_251216.png

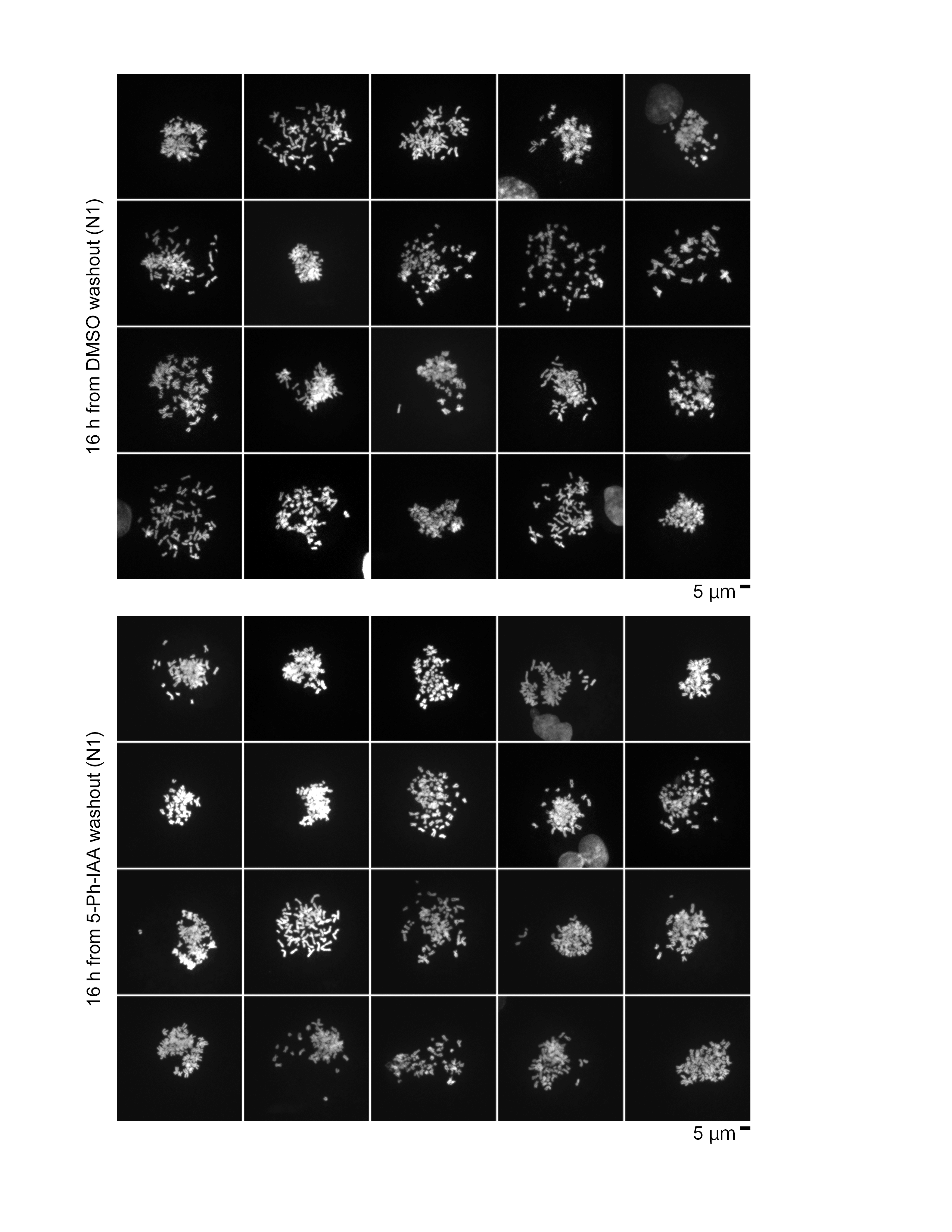

### Supplemental_material_S2_N2_0h_251216.png

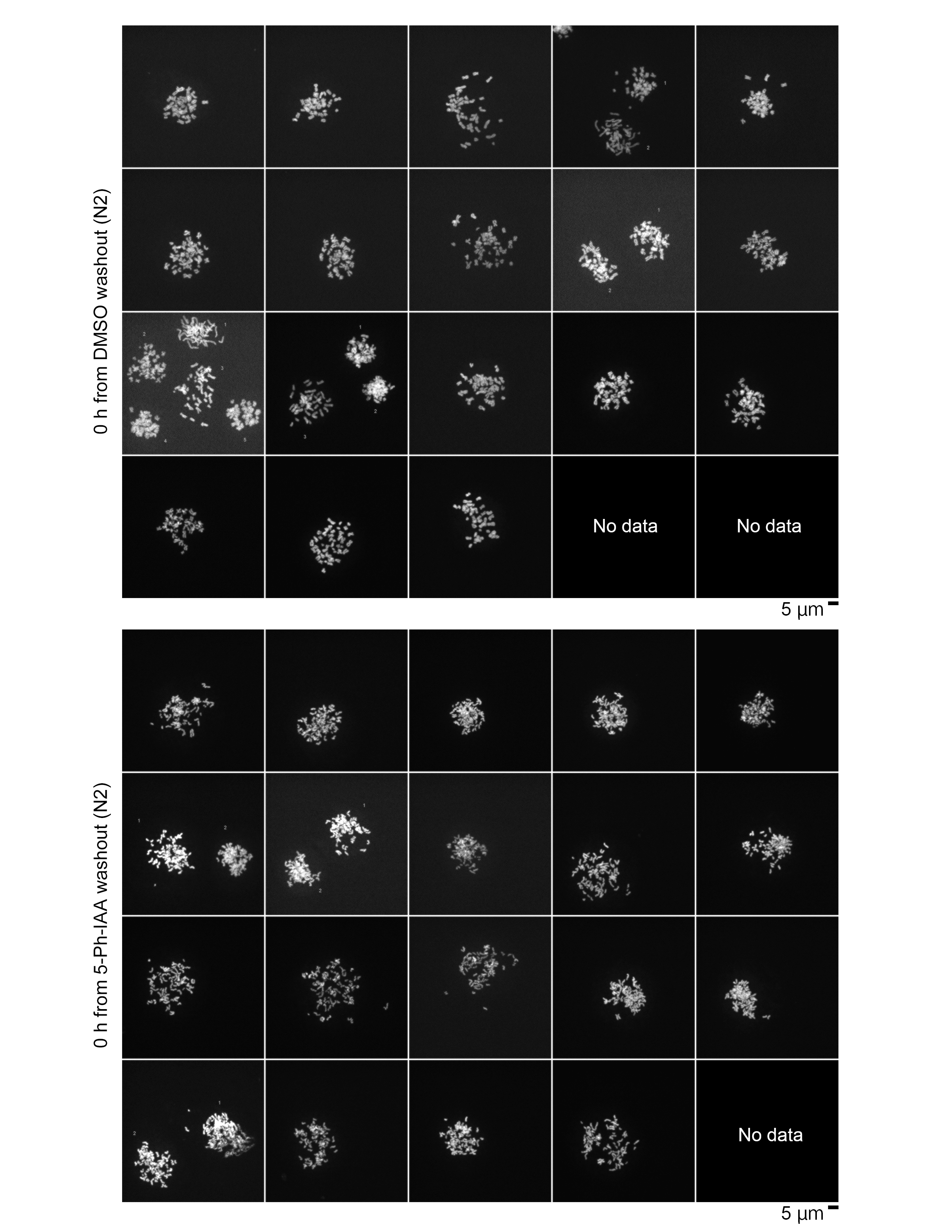

### Supplemental_material_S2_N2_16h_251216.png

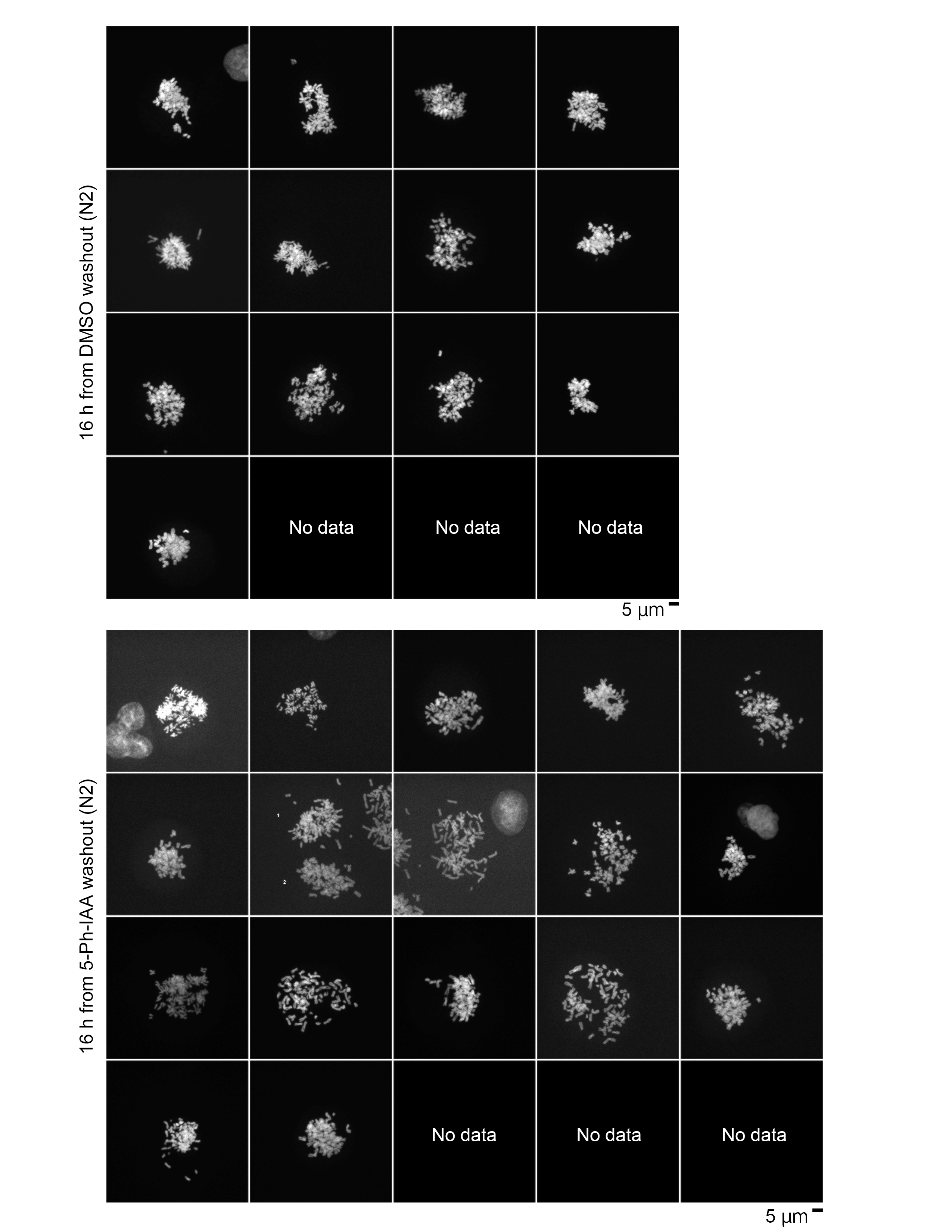

### Supplemental_material_S2_N3_0h_251216.png

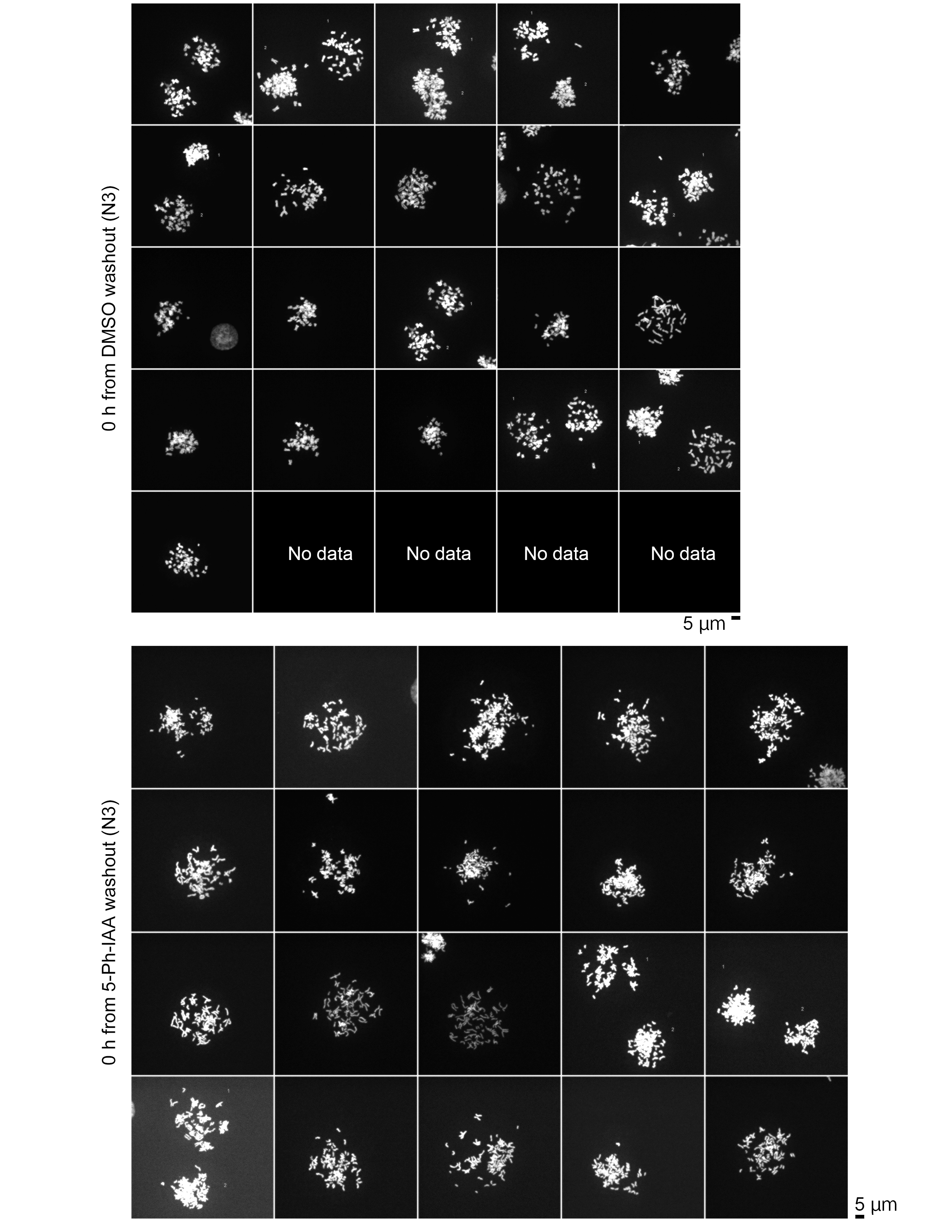

### Supplemental_material_S2_N3_16h_251216.png

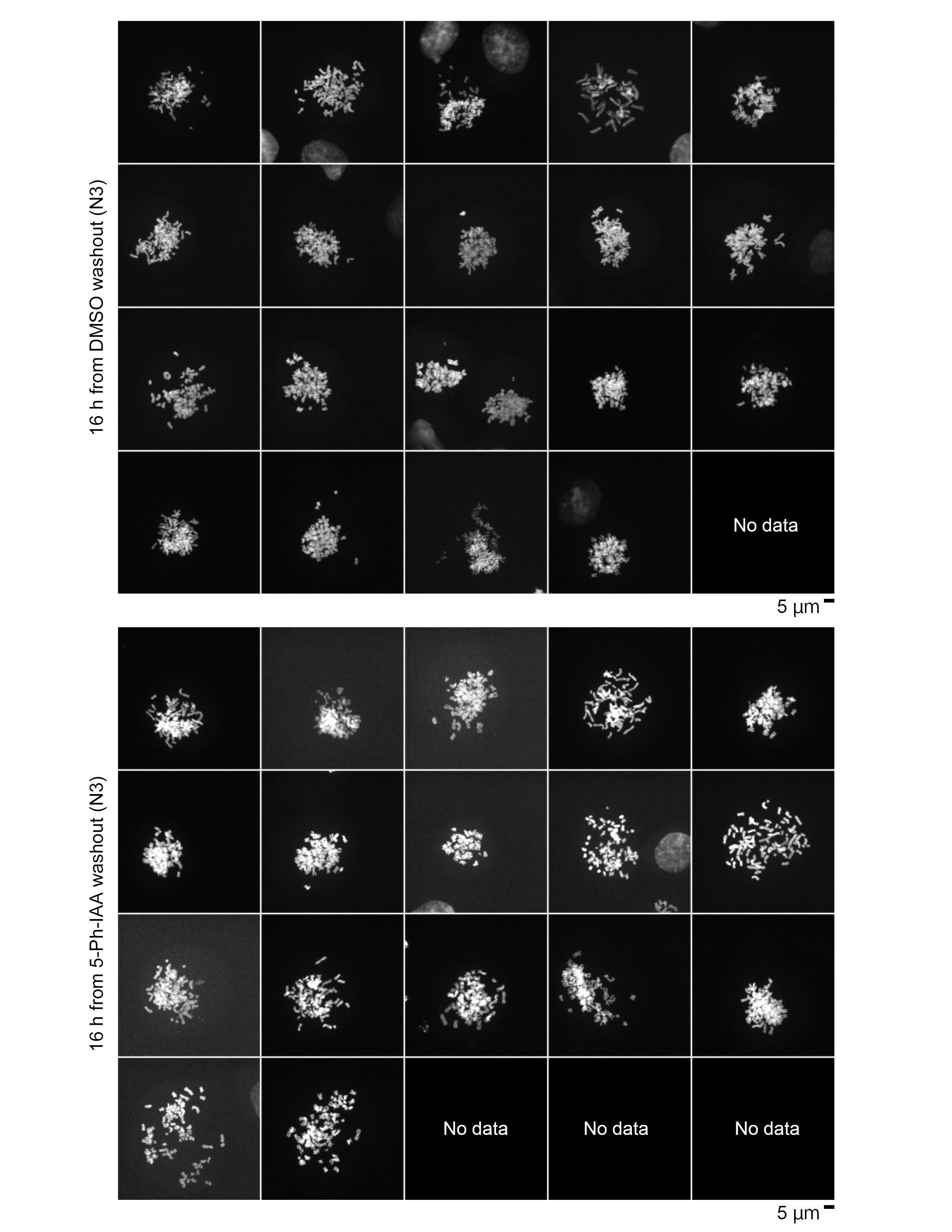
